# A unified rodent atlas reveals the cellular complexity and evolutionary divergence of the dorsal vagal complex

**DOI:** 10.1101/2024.09.19.613879

**Authors:** Cecilia Hes, Abigail J. Tomlinson, Lieke Michielsen, Hunter J. Murdoch, Fatemeh Soltani, Maia Kokoeva, Paul V. Sabatini

## Abstract

The dorsal vagal complex (DVC) is a region in the brainstem comprised of an intricate network of specialized cells responsible for sensing and propagating many appetite-related cues. Understanding the dynamics controlling appetite requires deeply exploring the cell types and transitory states harbored in this brain site. We generated a multi-species DVC cell atlas using single nuclei RNAseq (sn-RNAseq), by curating and harmonizing mouse and rat data, which includes >180,000 cells and 123 cell identities at 5 granularities of cellular resolution. We report unique DVC features such as *Kcnj3* expression in Ca^+^-permeable astrocytes as well as new cell populations like neurons co-expressing *Th* and *Cck*, and a leptin receptor-expressing neuron population in the rat area postrema which is marked by expression of the progenitor marker, *Pdgfra*. In summary, our findings demonstrate a high degree of complexity within the DVC and provide a valuable tool for the study of this metabolic center.

## 1. Introduction

Regulating appetite in response to changing environmental conditions to maintain energy balance requires the intricate interplay of multiple organs. Across the central nervous system, multiple discrete brain regions and cell types control appetite[1]. Within the brainstem, the dorsal vagal complex (DVC), comprising the area postrema (AP), nucleus of the solitary tract (NTS) and dorsal motor nucleus of the vagus (DMV), acts as a key nexus point for metabolic signals, evidenced by the dense expression of many relevant hormone receptors[2]. Separately, the DVC also serves as the primary site of gut-based signals via vagal afferents[3–5]. Functionally, the DVC was long considered a site promoting meal termination[6] but growing evidence suggests it also controls long-term energy balance[7].

Much of the interest in DVC neurons’ role in appetite and energy balance stems from their role as therapeutic targets for obesity and anorexia in cancer cachexia[8, 9]. For many years, most pharmacotherapies failed to produce efficacious and long-lasting weight loss in obesity and increase appetite in cancer cachexia[10–12]. More recently, however, the glucagon-like peptide 1 receptor (GLP1R) agonists are eclipsing the weight loss achieved by previous generations of therapeutics[13]. While expressed across the central nervous system, GLP1R expression is particularly enriched in DVC neurons[14] and it is the DVC that mediates many of the effects of GLP1R agonists[11]. Separately, an emerging strategy for treating cancer cachexia is inhibiting growth differentiation factor 15 (GDF15) signaling in the DVC by blocking the action of GDNF Family Receptor Alpha Like (GFRAL)[15], the receptor for GDF15[16]. Given the clinical significance of the DVC for this spectrum of conditions, understanding the molecular heterogeneity and function of this region is essential.

Traditionally, identification and ensuing study of DVC cell types was limited to known neurotransmitter and receptor-expressing cell types. For instance, the cholecystokinin and monoamine-expressing cells, demarcated by tyrosine hydroxylase (*Th*) expression, were previously shown to be non-overlapping cell populations with unique roles in appetite suppression[17]. More recently, the advent of single-cell mRNA sequencing approaches made unbiased molecular censuses of heterogenous populations possible. Such studies focusing on the mouse have revealed the complexity of the mouse DVC, comprised of dozens of transcriptionally distinct cell types[2, 18, 19]. Moving forward, unifying information from multiple single-nuclei RNA sequencing (snRNA-seq) datasets of the DVC into a reference atlas is needed to more fully understand the cells within the DVC and generate a common and scalable classification for cell identities. Such a reference atlas is also needed to contrast the cell types found within the mouse to other research models, including the rat, which are prominent models for DVC biology[2, 20].

To address this, we generated a comprehensive snRNA-seq based atlas of the mouse and rat DVC. This atlas features mouse and rat labelled cells with increasing granularity and an accompanying computational toolbox including the mouse and rodent atlas label transfer environment using treeArches. The mouse DVC atlas reveals a higher degree of cellular complexity than previously appreciated. The cellular census of the rat DVC exhibits many similarities to the mouse but also rat-specific cells including a novel population of leptin receptor and *Pdgfra*-expressing neurons localized to the AP. Using this unified atlas, we also uncover the cell types that transcriptionally respond to meal consumption which may hold the potential of suppressing appetite without gastrointestinal side effects.

## 2. Materials and methods

### 2.1. Experimental design

To generate a snRNA-Seq based atlas of the mouse dorsal vagal complex (DVC), we isolated the DVC from 30 adult C57BL/6J mice and subjected them to 10x Genomics-based single-nuclei RNA barcoding and sequencing. Mice were either fasted overnight (n=5), fasted overnight and then refed 2 hours prior to euthanasia (n=10), fed ad libitum (n=5), injected with LiCl (n=5) or injected with vehicle (n=5). Conserved cell clusters across these groups were then identified.

### 2.2. Animals

All animals were bred in the Unit for Laboratory Animal Medicine at the University of Michigan and the Research Institute of the McGill University Health Centre. Procedures performed were approved by the University of Michigan Committee on the Use and Care of Animals, in accordance with Association for the Assessment and Approval of Laboratory Animal Care and National Institutes of Health guidelines. Alternatively, procedures were approved by the animal care committee of McGill University.

#### 2.2.1. Mouse tissue for sn-RNA sequencing assay

Wild-type C57BL/6J mice (Jackson laboratories) were given ad libitum access to food (Purina Lab Diet 5001) and water in temperature-controlled (22°C) rooms on a 12-hour light-dark cycle with daily health checks. Mice were euthanized by isoflurane anesthesia and decapitated, then the brain was removed from the skull and aligned in a brain matrix. The cerebellar cortex was removed and the DVC was micro-dissected and flash frozen in liquid nitrogen (**Fig.S1**). To limit any circadian-driven gene expression changes, all animals were euthanized 4 hours following the onset of the light phase. Frozen tissue was stored at -80°C until use.

For assessment of feeding-related gene expression, mice were either ad libitum fed or food was removed at the onset of the dark cycle and the animals were fasted overnight. Animals were then euthanized the next morning under ad libitum fed or fasting conditions. A subset of fasted animals (n=5) were provided food and allowed to refeed for 2 hours prior to euthanasia.

#### 2.2.2. Mouse and rat tissue collection for *in-situ* hybridization and immunofluorescent assays

Animals were provided with ad libitum access to food (Purina Lab Diet 5001) and water in temperature-controlled (22°C) rooms on a 12 h light-dark cycle with daily health status checks. C57BL/6J mice (Jackson laboratories) or Sprague Dawley rats (Jackson laboratoties) were euthanized by isoflurane anesthesia followed by CO_2_ asphyxiation. Animals were then trans-cardially perfused with phosphate buffered saline for 3 minutes followed by 5 minutes perfusion with 10% formalin. Brain tissue was then collected and postfixed for 24 hours in 10% formalin before transfer into 30% sucrose for a minimum of 24 hours. Brains were then sectioned as 30 μm thick, free-floating sections.

#### 2.2.3. Rat tissue collection for snRNA sequencing assays

Wild-type Sprague Dawley rats (Charles River) were provided with ad libitum access to food (Purina Lab Diet 5001) and water in temperature-controlled (22°C) rooms on a 12-hour light-dark cycle with daily health checks. Rats were euthanized by intraperitoneal injections of pentobarbital and decapitated, then the brain was removed from the skull and aligned in a brain matrix. The cerebellar cortex was removed and the DVC was dissected and flash frozen in liquid nitrogen. Frozen tissue was stored at -80°C until use.

### 2.3 Isolation of nuclei from DVC tissue

Frozen tissue was pooled (5 pooled DVC from mice and 2 DVCs from rats) and homogenized in Lysis Buffer (EZ Prep Nuclei Kit, Sigma-Aldrich) with Protector RNAase Inhibitor (Sigma-Aldrich) and filtered through a 30mm MACS strainer (Miltenyi). Filtered samples were centrifuged at 500 rcf for 5 minutes at 4°C, and pelleted nuclei were resuspended in fresh-made wash buffer (10 mM Tris Buffer, pH 8.0, 5 mM KCl, 12.5 mM MgCl2, 1% BSA with RNAse inhibitor) before undergoing a second filtration and centrifugation. The pelleted nuclei were resuspended in wash buffer with propidium iodide (Sigma-Aldrich) and underwent fluorescence-activated cell sorting (FACS) on a MoFlo Astrios Cell Sorter to remove debris. PI+ nuclei were collected and centrifuged at 100 rcf for 5 minutes at 4°C and resuspended in wash buffer to obtain a concentration of 750-1,200 nuclei/μL in preparation for sequencing.

### 2.4. Single-nuclei RNA-sequencing

Library preparation was performed by the Advanced Genomics Core at the University of Michigan. RT mix was added to target approximately 10,000 nuclei recovered per sample and loaded onto the 10× Chromium Controller chip. The Chromium Single Cell 3′ Library and Gel Bead Kit v3, Chromium Chip B Single Cell kit, and Chromium i7 Multiplex Kit were used for subsequent RT, cDNA amplification, and library preparation, as instructed by the manufacturer. Libraries were sequenced on an Illumina NovaSeq 6000 (pair ended with read lengths of 150 nt).

### 2.5. Processing of the snRNA-sequencing files

From the snRNA-seq, we obtained four FASTQ files (two per sequencing lane, containing forward and reverse sequences respectively) per mouse treatment and two files with the forward and reverse sequences for the rat samples. We pre-processed the murine FASTQ files using either CellRanger 5.0.1 with inclusion of introns or CellRanger 7.0.1 with default parameters (which include intronic sequences) to align sequencing reads to the murine genome[21]. We aligned to the *Mus musculus* refdata-gex-mm10-2020-A reference genome. For the rat files, we used CellRanger 3.0.1 with no introns inclusion and aligned using the *Rattus norvegicus* genome assembly Rnor_6.0[21]. We obtained 110,167 and 13,360 cells from the murine and rat experiments, respectively. In addition to these two datasets, we also used the following steps to process the murine Dowsett dataset[18] publicly available as a raw gene expression matrix set of files. We processed the gene expression matrices in R[22] with Seurat v5 and SeuratObject v5 to remove low quality cells[23]. For processing and analysis, R v4.3.1[22] was used. We filtered cells with ≥500 number of genes mapped and >1.22 RNA unique molecular identifiers (UMIs) counts/genes mapped ratio (based on the first 2 percentiles per sample) (**Table S1**). Subsequently, we merged the data per treatment, calculated the median RNA content of clusters at resolution 1.0 and those below the first quartile of the median distribution were removed (**Table S2**). Additionally, we used DoubletFinder v2.0.3 based on the expected doublets rates by 10X Genomics after adjusting for homotypic doublets modeled as the sum of squared annotation frequencies[24] (**Table S3**). On each iteration we processed the data with libraries scaled to 10,000 UMIs per cell and log-normalized. We identified the most variable genes computing a bin Z-score for dispersion based on 20 bins average expression. We regressed UMI counts and used principal component (PC) analysis for dimensionality reduction on to the top 2,000 most variable genes. We used the first 30 PCs for *k*-nearest neighbors clustering and for uniform manifold approximation and projection (UMAP) projections using Seurat v5[23] default parameters. Murine samples integration in a single matrix was done with Harmony[25], after which clustering and UMAP projections were performed using the harmony embeddings instead of PCs.

### 2.6. Mapping of our murine data to existing brain databases

We mapped the resulting murine integrated high quality singlets to the celldex v1.12.0 built-in mouse RNA sequencing reference and other four mouse databases[26–29] (**Table S4**) using SingleR v2.4.1[30] and scRNA-seq v2.16.0[31] packages in R[22]. Since each database has its own cell type nomenclature, in our murine dataset we established a ‘likely-cell type’ to be unanimous among all databases, for example if all databases labeled a cell as ‘neuron’. We initially labeled the cells with conflicting or unassigned label by one or more databases as ‘unknown’. To confirm the cell type of all the cells labeled and to identify the unknown cell identities, we mapped 473 markers from the literature. For this, the average expression per marker gene was obtained on scaled data for each cluster at resolution 1.0 (i.e. 48 clusters) (**Table S5**). We also manually visualized UMAP projections from all these markers in our dataset. Some clusters (e.g. cluster 27) showed in the previous step to contain cells belonging to different cell types, so we subset and reprocessed these clusters using the Harmony embeddings to assign their cell identity based on marker expression at the lowest resolution level that allowed to separate them.

### 2.7. Assigning high resolution clustering-based cell identity in the murine dataset

In the murine dataset, for all non-neuronal clusters at resolution 1.0 gene expression markers were obtained using the FindConservedMarkers function from Seurat v5[23] package in R[22]. The function was run using the MAST algorithm[32] on scaled data with a log fold-change (FC) threshold of 0.25. Only upregulated marker genes detected on 40% of all cells in that cluster, were retained. Since the FindConservedMarkers gives a log2FC output per sample, we calculated the average log2FC across samples and the proportion of samples for which a gene had an adjusted p-value ≤0.05. A marker gene was considered as such if it had an average log2FC>1 across samples and was significant (i.e. had an adjusted p-value ≤0.05) in more than 80% of the samples. Furthermore, neurons were subset, reprocessed and subjected to new clustering through Seurat v5[23]. We obtained the gene markers per cluster at resolution 1.0 as described. We named the clusters based on the known functions of the upregulated/downregulated genes (e.g. myelinating oligodendrocytes), peculiarities of the cell groups (e.g. Ca^+^-permeable astrocytes) or their upregulated gene expression markers if the previous methods were unsuitable (e.g. Ano2 neurons) (**Table S6**). All visualizations, unless otherwise stated, were done using ggplot2 v3.5.0 and ComplexHeatmap v2.18.0 packages in R[22].

### 2.8. Oligodendrocytes trajectory inference

We subset the oligodendrocytes (i.e. pre-myelinating, myelinating intermediate and myelinating) in our murine dataset and placed each cell on an inferred cellular trajectory using their transcriptomic data in R[22]. We used the SCORPIUS v1.0.9 package[33] with default parameters to perform dimensionality reduction by PCA on log-normalized counts and to obtain the trajectory image, with a random seed of 1,000. We additionally performed the analysis on the Harmony embeddings from the original data integration.

### 2.9. Gene expression files format interconversions

To convert Seurat (‘RDS’) DVC datasets to ‘h5ad’ format and vice versa, we used reticulate v1.35.0[34] and anndata 0.7.5.6[35] packages in R[22] with import of scipy, scanpy, numpy and anndata modules from Python v3.9[36]. To convert .txt, .csv and .tsv datasets (i.e. the Ludwig dataset[2], the spinal cord dataset[37] and the cortex/hippocampus database[38]), we converted those to ‘h5ad’ format in Python v3.9[36].

### 2.10. Murine DVC hierarchy construction

We converted our murine labeled count matrix to h5ad. We used treeArches[39] in Python v3.9[36] to create a manual tree using the three layers of cellular granularity in our database. We reprocessed the samples using the treeArches pipeline[39] to normalize the count data (counts normalized per cell and then log-normalized). After multiple tests, we determined that the 2,500 most variable genes was the optimum parameter for the integration of other datasets based on our murine data, therefore we identified the top 2,500 most variable genes and integrated the samples using scVI and treeArches[39, 40]. This reference latent space obtained after integration was used to generate the UMAP embeddings. This sample integration was done to ensure that inter-sample variations were removed for the cell identity steps. We trained our manual tree based on the cell identity layer-3 labels using default parameters.

We subset the Ludwig dataset[2] to match the initial 2,500 most variable genes from our datasets and performed surgery to incorporate labels from the Ludwig dataset[2]. The two layers of granularity established by Ludwig[2] were combined in the variable “identity_layer3” to contain the highest granularity of cell identity for neurons and non-neuronal cell types. Next, we normalized the count data as specified for our murine dataset, and mapped the Ludwig dataset on the reference latent space using scArches. We then used the learn_tree function with default parameters for hierarchical progressive learning from scHPL v1.0.5[41] which yielded a hierarchy of harmonized cell labels from both datasets. We printed all trees using matplotlib v3.8.2[42] and scHPL v1.0.5[41]. All steps performed and scripts used in this phase are available in the GitHub repository: https://github.com/LabSabatini/DVC_cell_atlas.

#### 2.10.1. Corroboration of the validity of our murine cell hierarchy using treeArches prediction modality

Using treeArches[39] and its dependencies in Python v3.9[36] we compared the original cell labels in our and Ludwig datasets with the resulting prediction of the label for each cell using our learned hierarchy. We used the predict_labels function from scHPL v1.0.5[41] with default parameters, and the latent representation obtained when we constructed our murine hierarchy. We compared the original and the predicted labels through a heatmap visualization. Next, we predicted the labels of new data, a murine dataset by Dowsett which comprises the NTS, a part of the DVC[18]. We processed the Dowsett data similarly to the Ludwig dataset and obtained the query latent representation. We then used the predict_labels function from scHPL v1.0.5[41] with default parameters to transfer the labels in our hierarchy to the Dowsett dataset based on gene expression similarity among the 2,500 initial most variable genes. We then compared the resulting labels from this prediction to the manual mapping of our cell identities based on maker expression (**Table S7**) using UMAP embeddings in R[22].

### 2.11. Labeling of our rat DVC dataset

We processed our rat dataset as previously described to obtain high-quality singlets. Based on the three layers of cell identity that we created to label the murine data from our experiments, we calculated the average expression of the top 5 marker genes for each of the layer-3 identities (except duplicated genes and mouse-specific genes) on every rat cluster to identify those that were equivalent and could be labeled as the murine data (**Tables S6** and **S8**). Since this dataset is smaller than the mouse data, we decided to use clusters at resolution of 2.0 to capture more heterogeneity and delineate better the non-neuronal cell populations. Clusters 27 and 29 were further subset and reclustered since we found them to contain a mix of endothelial cells, mural cells, tanycytes and fibroblasts based on marker expression. Microglia and mixed neurons were sub-clustered as well. The lowest resolution level that allowed to separate populations was used to label the cells. Neurons were also subset and we mapped the murine neuronal classes markers to resolution 2.0 markers which were manually curated to delineate better the different populations within them. For all cell identities, and because we only applied one treatment to the rats (i.e. fed ad libitum), we used the FindMarkers function from Seurat v5[23] package in R[22] to obtain gene expression markers of each rat cell population. The function was run using the MAST algorithm[32] with same parameters used for the murine data (**Table S9**).

### 2.12. Construction of the rodent DVC hierarchy

We then integrated our rat labeled dataset to the murine samples in Python v3.9[36] using the initial top 2,500 most variable genes initially defined in our mouse data. Incorporation of rat labels was done as previously described for the Ludwig dataset using the learn_tree function for hierarchical progressive learning from scHPL v1.0.5[41]. The cell populations that were not rejected but placed at the bottom of the tree were moved manually to their belonging parent branches. For example, the ‘smooth_muscle_rat’ label was moved to be a children node of the mural cells node. If the parent branch did not exist, we created it. For example, the ‘immunity_related_rat’ label belonged to a new rat neuronal class, therefore, we incorporated this parent node within the neurons node and then we moved the ‘immunity_related_rat’ label to be a children node. After manual curation, we re-trained the tree. We printed all trees using matplotlib v3.8.2[42] and scHPL v1.0.5[41]. All steps performed and scripts used in this phase are available at the GitHub repository https://github.com/LabSabatini/DVC_cell_atlas for reproducibility.

### 2.13. Mapping gene expression in other brain-site snRNA-seq datasets

We downloaded labeled datasets from six brain regions different from the DVC: the HypoMap, spinal cord, cortex, hippocampus, pons and forebrain databases[37, 38, 43, 44]. For the HypoMap data, we subset the database by randomly selecting cells belonging to 32 included hypothalamic samples. We processed the datasets in Seurat v5[23] as previously described and obtained Harmony[25] embeddings to obtain UMAP projections. The labels used in the visualization are the original curated labels provided on each dataset. Neurons were further subset from the processed datasets and reprocessed as previously described. The other datasets were processed in the same manner after subsetting astrocytes. The pons and forebrain data was downloaded using scRNAseq v2.16.0 packages in R through SingleR v2.4.1[30].

### 2.14. Differential gene expression analysis in murine treatments

We subset the murine dataset neurons and glial cells at layer 2 granularity of identity and obtained differential gene expression for pair-wise comparison of treatments (i.e. refed versus ad libitum fed, refed versus fasted and LiCl versus vehicle injected mice). We used the FindMarkers function from Seurat v5[23] package in R[22] using the MAST algorithm[32] on log-normalized counts with a minimal log fold-change threshold of 0.1 and default parameters. The number of cells per treatment can be found in **Table S10**.

### 2.15. Immunofluorescent staining

Free-floating brain sections from mouse and rat were washed with PBS, three times for five minutes. The sections were then blocked for one hour in PBS containing 0.1% Triton X-100 and 3% normal donkey serum (Fisher Scientific). The sections were incubated overnight at room temperature with Goat anti-PDGFRA (Invitrogen, CAT#, 1:200), HuC/HuD (Invitrogen, A-21271, 1:30). The following day, sections were washed and incubated for two hours with Alexflour488 and Cy3 conjugated secondary antibody (Jackson immunoresearch, 1:500). Tissues were then washed three times in PBS before being mounted onto glass slides, covered in mounting media (Fluoromount-G, Southern Biotechnology) and coverslipped. Imaging was performed using an Olympus BX61 microscope and a Zeiss LSM780-NLO laser scanning confocal microscope equipped with IR-OPO lasers at the Molecular Imaging Platform, Research Institute of the McGill University Health Centre (RI-MUHC), Montreal, CA.

### 2.16. *In-situ* hybridization and imaging

Brain sections containing the DVC from four wild-type C57BL/6J mice obtained as described above, were fixed on glass slides and stored at -20°C for no more than 36 hours. We followed the manufacturer’s ACD 323100 user manual for the RNAscope® Multiplex Fluorescent Reagent Kit v2 Assay for fixed frozen tissue samples. Briefly, we rinsed the slides with 1X PBS and incubated them at room temperature for 10 minutes after adding ∼1-2 drops of H_2_O_2_ per section. We removed the H_2_O_2_ from slides and rinsed them twice with distilled water (diH_2_O). Next, we submerged the slides in 200 mL of hot diH_2_O for 10 seconds followed by 200mL of RNAscope® 1X Target Retrieval Reagent for 5 minutes, using a steamer with lid. After briefly transferring the slides to room temperature diH_2_O, we submerged them in 100% ethanol for 3 minutes and allowed them to air dry. We applied approximately 1-2 drops of RNAscope® Protease III to each section and incubated them at 40°C for 30 minutes in a HybEZ^TM^ oven with diH_2_O wet paper in the tray. After rinsing the slides with diH_2_O, we hybridized the probes at 40°C for 2 hours.

From this point on, we performed all steps by incubating the slides at 40°C in the HybEZ^TM^ oven with humid tray followed by rinse using RNAscope® 1X Wash Buffer. We applied approximately 1-2 drops of RNAscope® reagents AMP1, AMP2, AMP3 and then intercalated HRP channel (15 minutes), fluorophore (30 minutes) and HRP blocker (15 minutes) for each channel present in the probe mix. The mix of probes used for the *Th*/*Cck* co-expression assay was RNAscope® Mm-Th-C2 (317621-C2) and Mm-Cck-C3 (402271-C3) diluted in probe diluent as specified in the user manual. For *Hcrt* we used RNAscope® Mm-Hcrt-C2 (490461-C2) and for *Agrp*, Mm-Agrp (400711), in a dilution as specified in the user manual. The used fluorophores were Cy3 for *Cck* and *Hcrt* (1:1,500), and Cy5 for *Th* and *Agrp* (1:1,000) diluted in RNAscope® TSA buffer. DAPI dye (1:1,000) was applied at the end of the assay, and the samples were stored for ∼48 hours at -4°C, protected from light, before imaging. We imaged the sections using a Zeiss LSM780-NLO laser scanning confocal microscope with IR-OPO lasers at the Molecular Imaging Platform at the RI-MUHC, Montreal, CA.

## 3. Results

### 3.1. The murine dorsal vagal complex cell classes

To generate a *de novo* snRNA-seq atlas of the mouse DVC, we isolated the DVC from 30 adult C57BL/6J mice and subjected them to 10X Genomics-based snRNA barcoding and sequencing (**Fig.1A**). From the CellRanger pre-processing pipeline[21] we obtained 110,167 cells of which 99,740 were retained after processing. Cell identities were initially mapped to 5 databases from different brain regions and then curated manually using 473 cell identity markers (**Tables S4-S5**). We distinguished clusters of neurons and glial, vascular and connective tissue cells, a first layer of cell identity (**Fig.1B; Table S9**). We further describe a second layer of cellular resolution, which are subgroups of the first layer that contain the cell classes (**Fig.1C; Table S7**).

**Figure 1:**
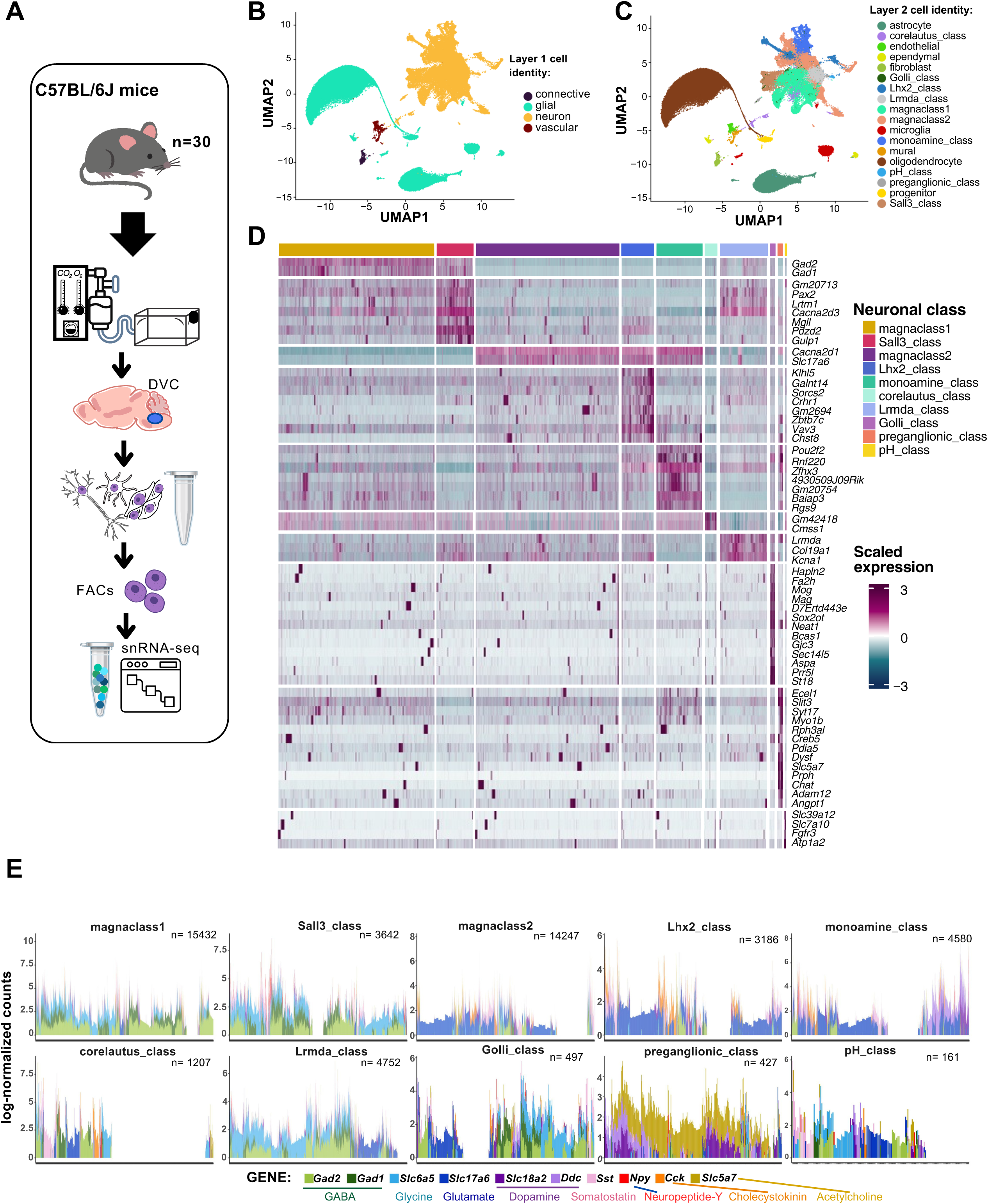
Two layers of cell identity for the murine DVC cells from snRNA-seq data. **A.** Adult C57BL/6J mice were subjected to 10X Genomics-based snRNA barcoding and sequencing after dissection of their DVC (n=30). **B.** Labeled UMAP plot of the mouse DVC (cells n=99,740) at the lowest resolution (layer 1) including 4 cell identities, and **C.** the cell class (layer 2) including 18. **D.** Heatmap of the main marker genes (average log2FC expression >80 percentiles of upregulated genes) for each neuronal class of the mouse DVC (MAST algorithm; neurons n= 48,131; adj. p<0.05). Each column is one cell and each row is one marker gene. **E.** Stacked barplot of log-normalized counts of genes for 10 transcripts (i.e. *Gad2*, *Gad1*, *Slc6a5*, *Slc17a6*, *Slc18a2*, *Addc*, *Sst*, *Npy*, *Cck* and *Slc5a7*) related to transmission/release of 8 neurotransmitters and neuropeptides, by neuronal cell class. Each bar represents one neuron (n=48,131). snRNA-seq=single-nuclei RNA-sequencing; DVC=dorsal vagal complex; FACs=fluorescence-activated cell sorting; UMAP=uniform manifold approximation and projection; FC=fold-change; adj.=adjusted; GABA=gamma-aminobutyric acid

Among the ten neuronal cell classes identified at layer 2 resolution, we found two major groups which we called ‘magnaclass1’ and ‘magnaclass2’ (**Fig.S2**). Magnaclass1 neurons predominately express genes required for inhibitory neurotransmission (e.g. *Gad1*, *Gad2*) and share transcriptomic markers with the Sall3-class of neurons, whereas magnaclass2 neurons express mostly excitatory transcripts (e.g. the glutamate transporter VGLUT2) also found in the Lhx2-class and monoamine-class (**Fig.1D-E; Fig.S2**). However, as described in other brain areas[45, 46], evidence of co-transmission/co-release of multiple primary neurotransmitters was found in at least 55% of neurons regardless of their cell class (**Fig.1D-E; Table S11**). We also identified 8 transcriptionally distinct neural classes in addition to the magnaclasses. Among these 8 classes we found two neural classes expressing traditionally non-neuronal genes; the Golli and pH-related classes. The Golli-class neurons are a relatively small cluster, comprising ∼1% of all neurons. Interestingly, Golli-class neurons express multiple myelin-associated genes such as *Mag* and *Mog*[47–49] normally expressed by oligodendrocytes (**Fig.1D; Table S6**). Additionally, we identified a class of neurons with genes normally upregulated in astrocytes (e.g. *Slc6a11*) and related to clearance and uptake of glutamate, gamma-aminobutyric acid (GABA), Ca^+^ and bicarbonate [50, 51], which we named ‘pH-related class’ (**Table S6**). Similar to Golli-class neurons, pH-related neurons were a relatively rare cluster (<1% of all neurons). While the corelautus-class neurons express very few genes at higher levels, among them are *Cdk8*, *Lars2*, *Gabra6* and *Fat2*. Expression of *Cdk8* is largely absent from the DVC but is enriched within the DVC-adjacent hypoglossal nucleus (**Fig.S3**) suggesting these cells are hypoglossal neurons. We also identified the Lhx2-Class which was enriched for the Parvalbumin (*Pvalb*) gene whose expression is largely excluded from the DVC. As *Pvalb* is highly expressed in the neighbouring cuneate nucleus (**Fig.S3**), we posit this class may belong to this anatomically adjacent area which was captured in our dissection. When interrogating snRNA-seq data from the hypothalamus[43] (**Fig.S4**), another brain site important for energy balance, we could not find clusters sharing all markers with our neuronal classes, indicating DVC-specific neuronal programs (**Fig.S4C**).

### 3.2. The murine DVC cells at their highest resolution

To better define the heterogeneity of the mouse DVC, we further assigned a clustering-based third layer of cellular resolution resulting in fifty cell identities, of which 15 are non-neuronal (**Fig.2A**). At this resolution we differentiated between two major groups of astrocytes. Differential expression in synapse-related factors revealed a group of astrocytes with high expression of the inwardly rectifying GIRK1 (*Kcnj3*) channel[52] and lack of expression of the AMPAr subunit GluA2 (*Gria2*), additionally rendering them K^+^/Na^+^/Ca^+^-permeable[53, 54] (**Fig.2B**). We speculate these astrocytes are specific to the DVC as we could not find *Kcnj3*+; *Gria2*-astrocytes in the hypothalamus, forebrain, cortex or hippocampus[38, 43, 44] (**Fig.S4D; Fig.S5**). Furthermore, although some regions close to the DVC such as pons and the spinal cord show some co-expression of *Kcnj3* and *Gfap*, the patterns are not to the same extent as observed in the DVC [44] (**Fig.S5F-G**). In addition, we distinguished a spectrum of DVC mature oligodendrocytes, including a group resembling the intermediate oligodendrocytes described in the other brain sites that decrease in number with age[37, 55] (**Fig.2C**). This gradient of expression of markers including *Fyn*, *Opalin* and *Anln* in oligodendrocytes (**Fig.2C**), do not respond to a developmental trajectory (**Fig.S6**). Although ‘myelinating-intermediate’ and ‘myelinating’ oligodendrocytes include cells with overlapping gene expression programs, there is no evidence that each oligodendrocyte will follow a single trajectory (**Fig.S6**). Furthermore, to properly separate some cell identities, we performed sub-clustering in three DVC clusters (i.e. clusters 23, 26 and 27) (**Fig.S7A**). This permitted us to label pre-myelinating oligodendrocytes, endothelial and mural cells, and distinguish two fibroblast sub-types (**Fig.S7**). One of these subtypes expresses high *Stk39* previously described in fibroblast stress-response[56] and so was termed ‘s-fibroblasts’ for stress-associated fibroblasts (**Fig.S7B**), which we failed to detect in the hypothalamus (**Fig.S4E**). Additionally, we excluded the possibility of monocytes in our microglial data (**Fig.S8A**), which had no evidence of microglial activation as we mapped minimal levels of the activation genes *Cxcl10*, *Cd5*, *Cxcl9* and *Zbp1*[57], therefore we posit that our DVC microglial cells are in a basal state (**Fig.S8B**). To further evaluate possible activation states in microglia, we performed sub-clustering and evaluated their marker genes (**Fig.S8C**). We have identified a group of microglia expressing oligodendrocyte markers and resembling a subpopulation of hippocampal microglia in an Alzheimer’s disease model which is disease-protective and synaptic-function-supporting[58], and a cortical/subcortical phagocytic subpopulation in a multiple sclerosis model[59]. A second group expresses *Ndrg2* and receptors for GABA and glutamate which have been related to microglia that may induce neuronal damage[60, 61] (**Fig.S8D**).

**Figure 2:**
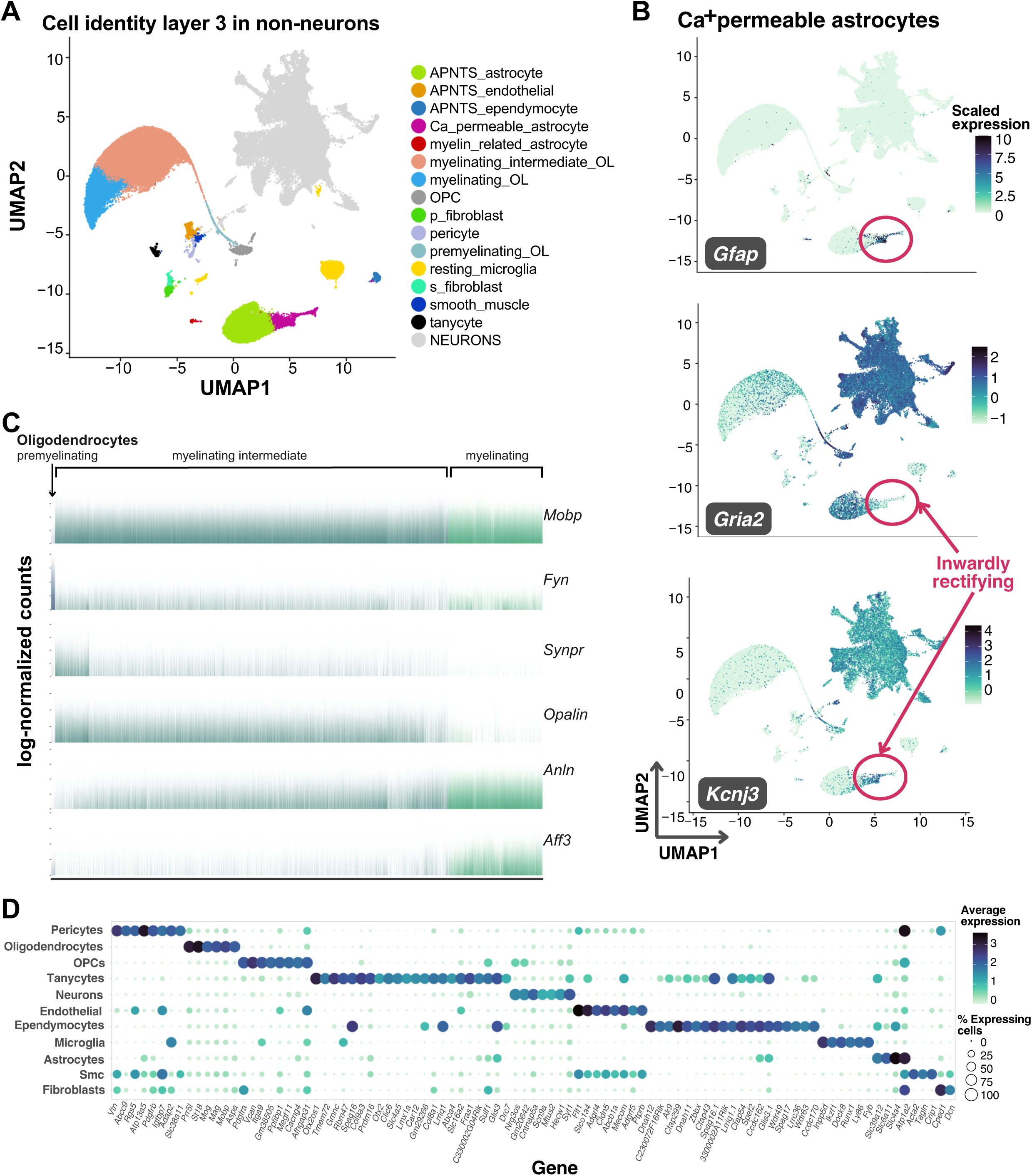
Third layer of non-neuronal murine DVC cell identities. **A.** Labeled UMAP plot of the mouse DVC with the 15 non-neuronal cell identities shown (total cells n=99,740; non-neuronal cells n=51,609). **B**. UMAP plot showing scaled expression of *Gfap*, *Gria2* and *Kcnj3* genes in Ca^+^-permeable astrocytes (circled; total cells n=99,740; Ca^+^-permeable astrocytes n=1,719). **C**. Barplot of log-normalized expression of six genes in oligodendrocytes (premyelinating n=267; myelinating intermediate n=25,692; myelinating n= 6,170). Only premyelinating oligodendrocytes lack expression of *Mobp*, involved in myelin formation. Reduction of *Synpr* and *Opalin* expression is observed as *Anln* and *Aff3* increase in myelinating oligodendrocytes. **D.** Balloon plot of the main cell types of non-neuronal cells and neurons showing average log-normalized counts of their marker genes (MAST algorithm; adj. p<0.05). Markers shown are upregulated genes in >80% of cells per group with average log2FC>4, or upregulated in >70% of cells with average log2FC>8. DVC=dorsal vagal complex; UMAP=uniform manifold approximation and projection; APNTS=Area postrema and nucleus of the solitary tract; OL=oligodendrocyte; OPC=oligodendrocyte precursor cell; Smc=smooth muscle cells

We also obtained the DVC gene expression markers of the main cell types in the central nervous system and neurons (**Fig.2D**). To more thoroughly classify DVC neural types, we subset and re-clustered the 10 neural classes from layer 2 which resulted in 35 neuronal identities as a third layer of granularity (**Fig.3A; Table S6**). These identities include ‘preganglionic’ or *Chat*-expressing neurons and a large number of neurons with unspecific markers which we called ‘mixed neurons’ (**Table S6**). Of the mixed neuron classes, two are largely excitatory (Mixed neurons 3 and 4) and two largely inhibitory, however, cells in the four mixed neuron classes share similar transcriptional profiles with many other neural classes. When we performed sub-clustering on mixed neurons, we could distinguish a total of 10 subtypes from the original 4 mixed neuronal identities (**Fig.S9**); however, transcriptional differences between these were subtle. Interestingly, a mixed neurons-like group of cells was previously described in the murine hypothalamus[26], suggesting multiple brain sites may contain neurons which lack highly differentiable transcriptional programs from snRNA-sequencing data.

**Figure 3:**
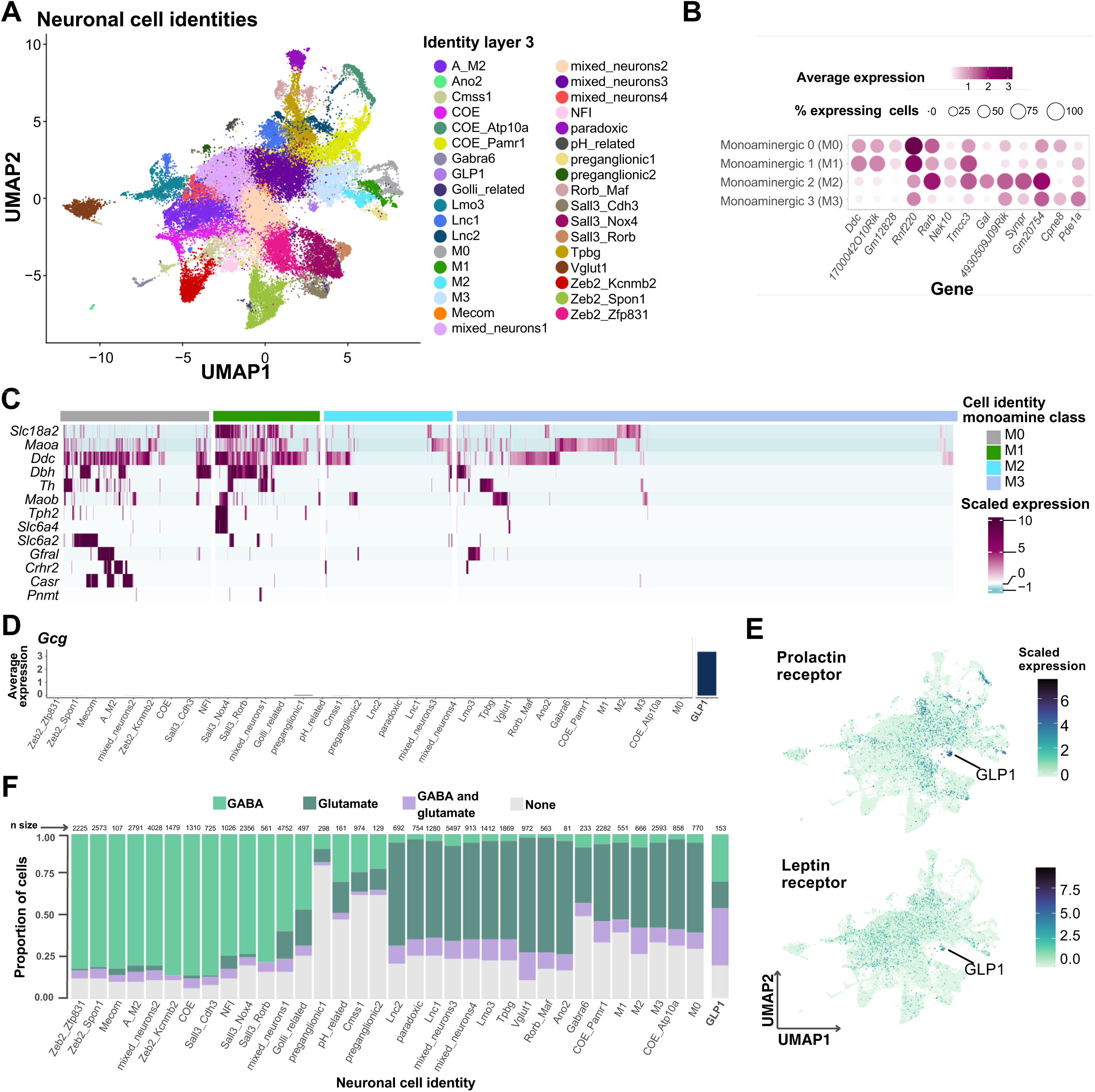
Neuronal populations of the murine DVC. **A**. Labeled UMAP plot with the neuronal layer 3 cell identities (neurons n=48,131). **B**. Monoamine class balloon plot showing average log-normalized expression of marker genes for each cell identity (MAST algorithm; M0 cells n=770, M1 cells n=551, M2 cells n=666 and M3 cells n=2,593; adj. p<0.05), and **C.** heatmap of expression of monoamine related genes. In the heatmap, each column represents one cell and each row, one gene. **D**. Barplot of *Gcg* average log-normalized expression by neuronal cell identity. E. UMAP plot showing the expression of leptin receptor and prolactin receptor genes in neurons. The GLP1 cluster is highlighted (GLP1 cells n=153). **F**. Stacked barplot of the proportion of cells expressing one or more GABA-related genes (i.e. *Slc32a1*, *Gad1*, *Gad2*) and glutamate-related genes (i.e. *Slc17a6*, *Slc17a7*). The proportion of cells per cell group co-expressing GABA and glutamate associated genes are shown in purple. Only cells with log-normalized counts >0 were considered to express the genes. DVC=dorsal vagal complex; UMAP=uniform manifold approximation and projection; Res.=resolution; GABA=gamma-aminobutyric acid

Additionally, we split the monoamine-class into four layer-3 cell identities (i.e. M0, M1, M2 and M3) with specific cell markers (**Fig.3B; Table S6**). M0 and M1 neurons express genes for enzymes synthesizing norepinephrine and serotonin, but M0 expresses minimal levels of the transporter VMAT2 (*Slc18a2*) (**Fig.3C; Table S6**). Due to the low proportion of cells expressing *Th* and *Tph2* in M2 and M3 neurons, we consider them monoamine-modulators as they seem to synthesize trace amines (**Fig.3C; Table S6**). Of note, the receptor for GDF15, GFRAL, is expressed heavily by a subset of M0 neurons, some of which co-express the calcium sensing receptor (*Casr*) gene (**Fig.3C**).

We named ‘GLP1’ a neuronal cluster with high expression of the pre-proglucagon gene (*Gcg*) (**Fig.3D; Table S6**), which also expresses high levels of both the prolactin and leptin receptors[62] (**Fig.3E**). Additionally, these cells have the highest proportion of DVC neuronal co-expression of glutamate- and GABA-related genes (**Fig.3F**); mainly *Gad2* with considerably lower expression of *Slc32a1*.

In our analysis we found a subset of monoaminergic neurons that express the cholecystokinin (*Cck*) gene (**Fig.4A**). While the *Cck* and *Th*-expressing DVC neurons are considered to be non-overlapping cell types with distinct physiologies[17, 63], but we quantified >10% of the *Th*-expressing neurons also expressing *Cck* in our snRNA-seq data. We further confirmed co-expression of *Th* and *Cck* results by *in-situ* hybridization (**Fig.4B**). The majority of this overlap occurs in cells belonging to the COE-Atp10a and monoamine class (i.e. M0, M3) cell identities (**Fig.4A**).

**Figure 4:**
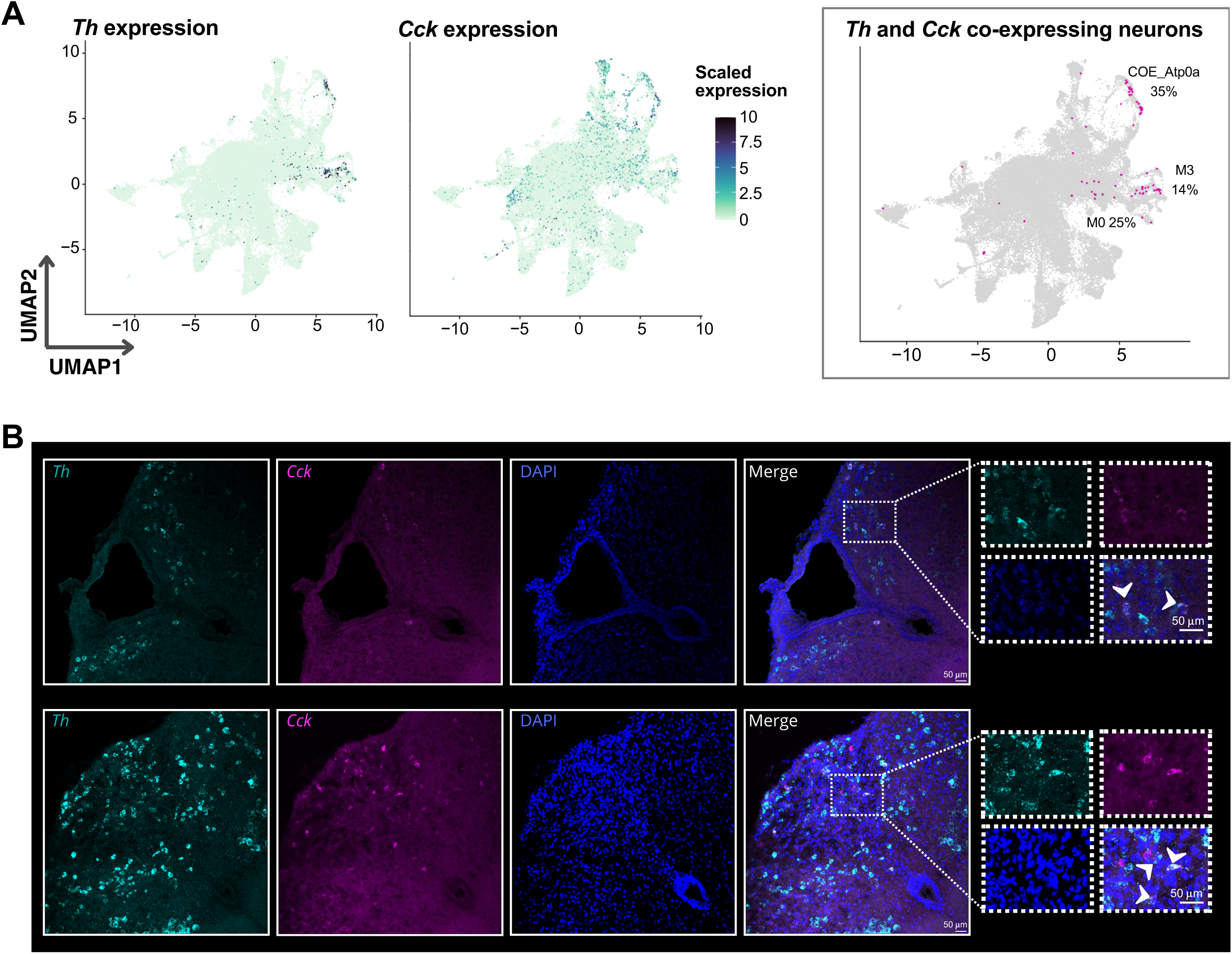
*Th* and *Cck* co-expression in the DVC. **A.** UMAP plot of neuronal *Th* and *Cck* scaled expression and cells with co-expression of both genes (neurons n=48,131; *Th*-expressing neurons n=764; *Cck*-expressing neurons n=3,821; *Th*/*Cck* co-expressing neurons n=80). On the right plot, cells highlighted in magenta are the co-expressing neurons. The three neuronal cell identities in which the majority of nuclear co-expression of *Th* and *Cck* mRNA is found, as well as the percentage of total co-expressing neurons, are shown. **B.** *In-situ* hybridization of *Cck* and *Th* mRNA in coronal mouse DVC sections corresponding to -7.2mm and -7.56mm relative to bregma showing overlap of *Th*/*Cck*. Some of the cells with overlapping signal are highlighted with an arrow in the enhanced merged image. DVC=dorsal vagal complex; UMAP=uniform manifold approximation and projection

### 3.3. The murine DVC cell atlas

To generate a comprehensive murine DVC database from multiple datasets, we utilized treeArches[39] to harmonize our snRNA-seq DVC labels with that from a publication by Ludwig and collaborators (i.e. the ‘Ludwig dataset’)[2] (**Fig.5A-B**). After building our initial cell hierarchy (**Fig.S10**), we integrated our dataset with the Ludwig dataset, giving a total of 171,868 cells (**Fig.5**B). Next, the labeled cell identities from the Ludwig dataset[2] were incorporated into our tree based on the similarity between the transcriptomic programs of our labeled identities and the Ludwig-labeled identities using progressive learning through scHPL[41] (**Fig.5A**). Some of the Ludwig dataset identities failed to be incorporated into our tree, like the oligodendrocyte precursor cells (OPCs), which contain a mix of OPCs and pre-myelinating oligodendrocytes (**Fig.S11A**). The mix of cell programs makes it impossible for treeArches to correctly add one branch of the Ludwig OPCs to the tree as ideally it would be added as a subpopulation of both the OPCs and pre-myelinating oligodendrocytes, which is not biologically possible. The resultant murine hierarchy has a fourth layer of cellular resolution as some of Ludwig identities, mainly neurons, are subgroups of our cell identities (**Fig.S11B**).

**Figure 5:**
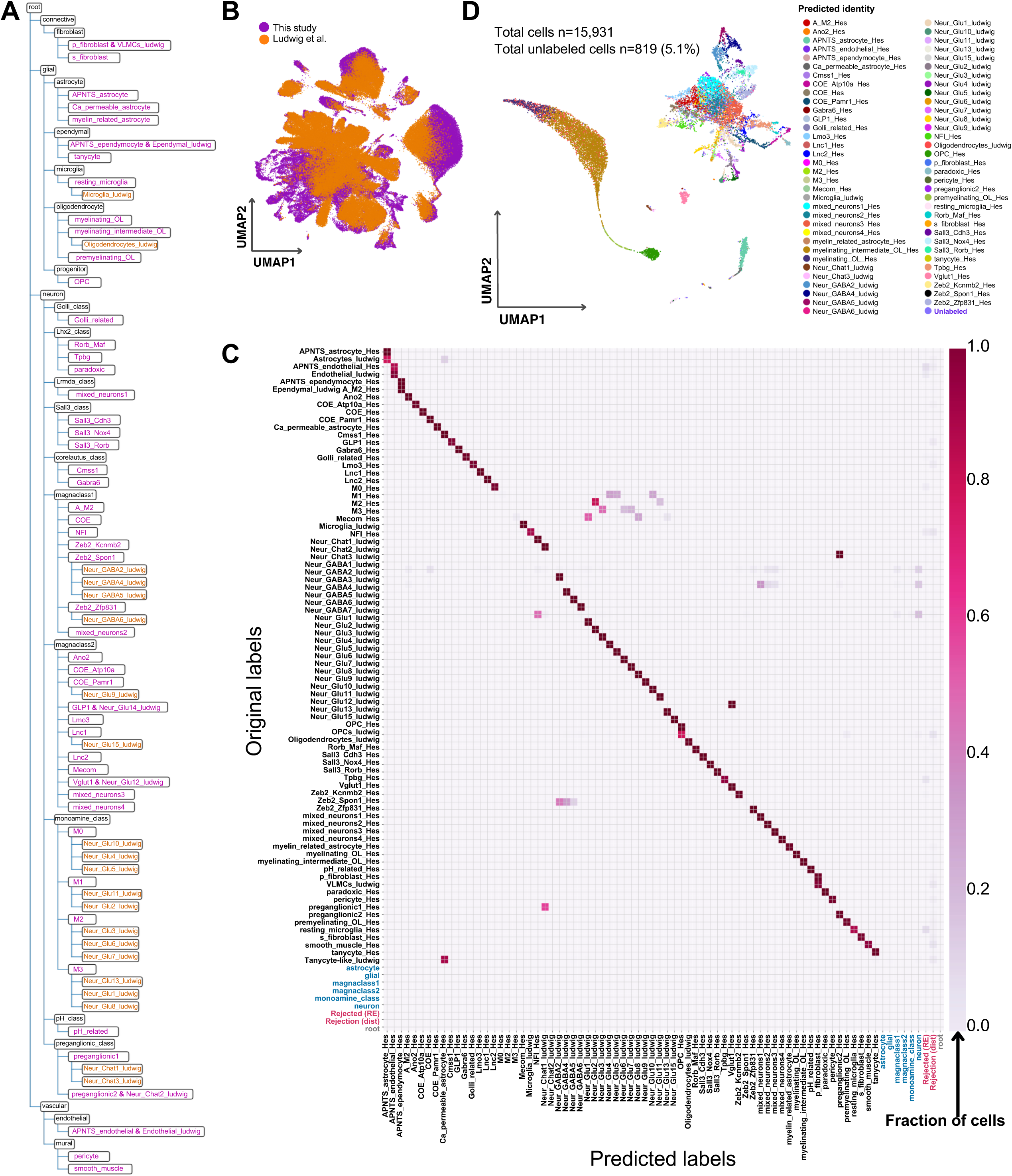
The murine DVC cell hierarchy. **A**. Dendrogram of the harmonized hierarchy which incorporates cells from Ludwig and our datasets. Orange cell identities are a fourth layer of cell identity resolution obtained as some of the Ludwig dataset identities are subgroups of our original cell identities at their highest resolution. Magenta labels represent layer 3 of cellular granularity. These two layers are considered high-resolution. **B.** UMAP plot of the integration between our and Ludwig datasets using treeArches (cells n=171,868). **C**. Pairwise heatmap showing the proportion of cells originally labeled in this study and in Ludwig dataset (y-axis), predicted to belong to each identity group (x-axis) by treeArches using our learned harmonized hierarchy. In blue are the labels considered non-specific for a high-resolution cell identity (n=2,324; 1.3%). In pink are the cell labels for rejected cells, therefore not assigned any identity by treeArches (n=3,108; 1.8%). **D**. UMAP plot of the Dowsett dataset labeled through treeArches using the learned harmonized hierarchy representation from our murine DVC atlas. The ‘unlabeled’ cells include those rejected by the algorithm and thus without an assigned identity, and those with an unspecific layer-3/4 label. For example, some were labeled ‘neuron’ or ‘monoamine-class’ but without a high-resolution cell identity. UMAP=uniform manifold approximation and projection; APNTS=Area postrema and nucleus of the solitary tract; Lnc=long non-coding; OL=oligodendrocyte; OPC=oligodendrocyte precursor cell; Neur=neuronal; COE=collier/Olf1/EBF transcription factor

We confirmed the validity of our cell hierarchy using three methods. First, by using the learned model to predict the labels of the cells we used to develop them (i.e. this study and the Ludwig dataset), for which we obtained highly concordant cell labeling (**Fig.5C**). Secondly, we used the learned model to predict the labels of another publicly available dataset from Dowsett and collaborators (i.e. the ‘Dowsett dataset’)[18] (**Fig.5D**). Using this approach, we were able to successfully label 95% of these cells at high resolution (i.e. layer 3 or 4 of cell identity granularity); further validating our combined DVC labels. Finally, we manually mapped the marker genes from Ludwig[2] and Dowsett[18] neuronal clusters in our dataset and confirmed the sub-groups of neurons among our identities pointed out by treeArches (**Fig.S11B**).

### 3.4. An integrated rodent DVC cell hierarchy

To identify rat DVC cell clusters and subclusters and contrast these with the mouse DVC atlas, we generated a *de novo* ad libitum-fed rat dataset (cells n=12,167) (**Fig.6A**) and labeled it by mapping the top 5 gene expression markers of each of our murine layer-3 cell identities (**Table S8**). After manual curation at three granularities (**Fig.6B-C; Table S9**), we obtained 52 DVC rat identities at high cellular resolution (**Fig.6C**) including a small group of ‘unspecific’ cells (n=35) that we could not confidently label because its top gene expression markers have unknown functions and none of our known identities’ genes clearly marked this population (**Fig.6B-C; Table S8**).

**Figure 6:**
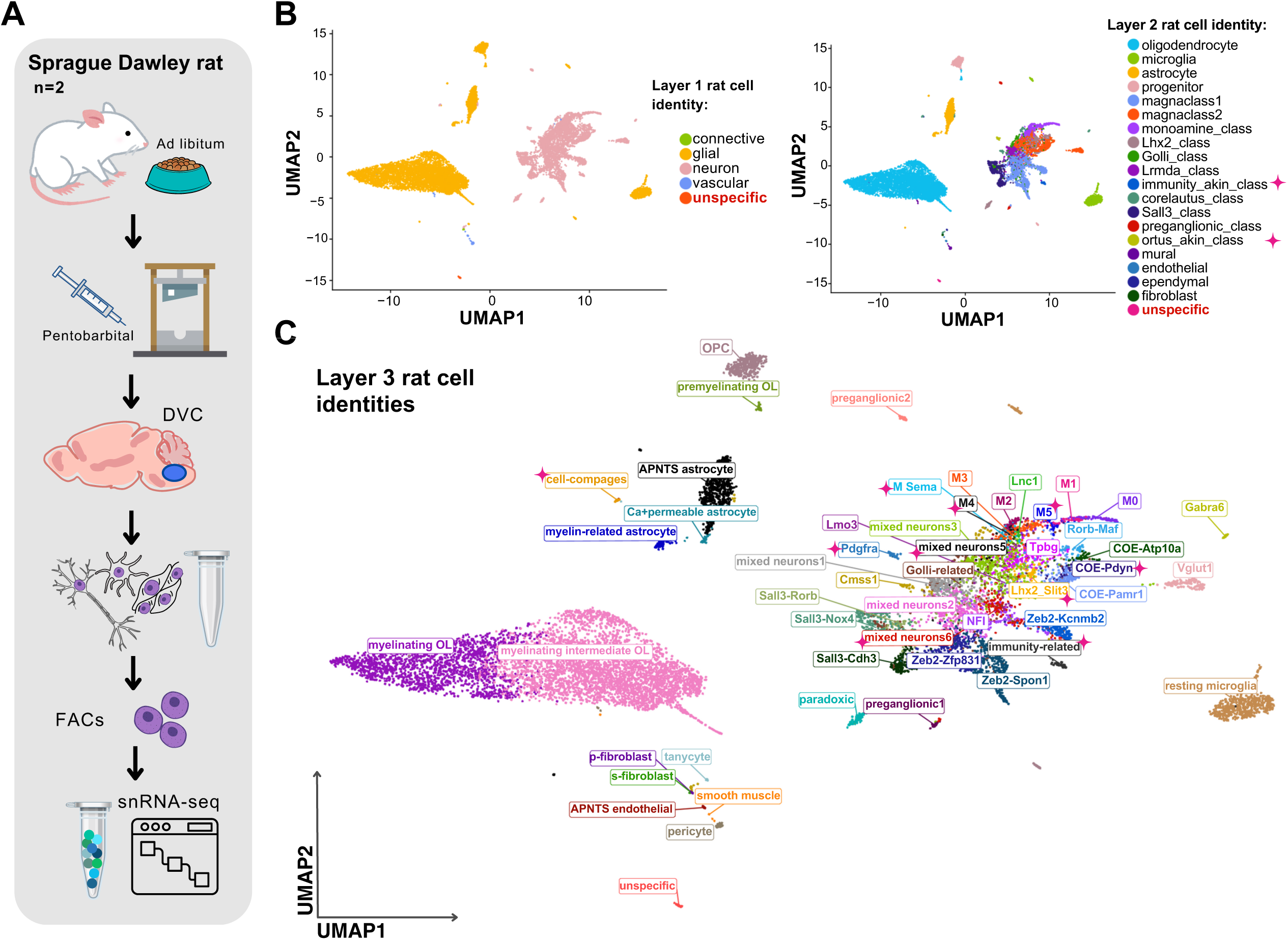
The snRNA-seq derived cell identities for the rat DVC. **A.** Schematic of the pipeline for snRNA-seq of the rat DVC. **B**. Labeled UMAP plots of the low resolution (i.e. layers 1 and 2 of granularity) and **C**. high resolution (i.e. layer 3) cell identities in our rat DVC dataset (cells n=12,167). We labeled four layer-1, nineteen layer-2 and fifty-two layer-3 cell identities. Those labels novel to this dataset and not present in the murine data are highlighted with a pink ♦ symbol. We found a small cluster (cells n=35) that we could not corroborate its identity at any layer of cellular granularity that we named ‘unspecific’. snRNA-seq=single-nuclei RNA-sequencing; DVC=dorsal vagal complex; FACs=fluorescence-activated cell sorting; UMAP=uniform manifold approximation and projection; APNTS=Area postrema and nucleus of the solitary tract; Lnc=long non-coding; OL=oligodendrocyte; OPC=cursor cell; Neur=neuronal; COE=collier/Olf1/EBF transcription factors

We established 10, layer-3 neuronal identities not previously found in the murine data (**Fig.6C; Fig.S12A; Table S9**). Notably, two of those did not belong to any of the existing murine neuronal classes (i.e. layer-2 of cellular granularity), therefore we created two new layer-2 classes (**Fig.6B**). One of these, the immunity-related neurons, have high *Csf1r* expression, as well as cystatin C and *Inpp5d* which are commonly associated with immunity-related processes in microglia and with immunomodulation in cancer[64, 65] (**Fig.7; Table S10**). The second neural class identified in rat but not mouse DVC were the ‘Pdgfra neurons’ (**Fig.7**). These neurons express *Pdgfra*, *Arhgap31* and *Itga9*, known markers for progenitor cells for which we named this cell class ‘ortus-akin’ (Latin: ‘origin’) because they are similar to cells giving origin to other cells[66] (**Fig.7A; Table S10**). Interestingly, this neuronal class not only expressed *Pdgfra*, but also other OPC signature genes **(Fig.7A**). Mapping the expression levels of metabolic-associated receptors revealed that a subset (∼15%) of both immunity-akin and ortus-akin express detectable *Gfral* mRNA but this is likely an underrepresentation due to the challenges in detecting lowly expressed transcripts such as *Gfral*. Prolactin receptor and leptin receptor (**Fig.7B**) were also found to be expresses by these neurons, although they seem to have a varied neurotransmitter profile (**Fig.S12B**). To confirm the presence of *Pdgfra*-expressing neurons in the rat DVC, we performed immunostaining in the mouse and rat brainstem. Within the mouse, we found PDGFRA-immunoreactivity(IR) in the two morphologically distinct cells; matching OPC and blood vessel-associated fibroblast morphologies(**Fig.7C**). However, within the rat, we identified PDGFRA-IR in cells with three morphologies, one of which resembled neurons with an enlarged cell body and protruding cellular processes (**Fig.7C**). To determine if these PDGFRA-IR cells were neurons, we co-stained with the neural marker, HuC/D and noted several PDGFRA and HuC/D copositive cells (**Fig.7D**); confirming our sequencing data that a PDGFRA-expressing neural class exists in the rat DVC and localizing these cells to the area postrema.

**Figure 7:**
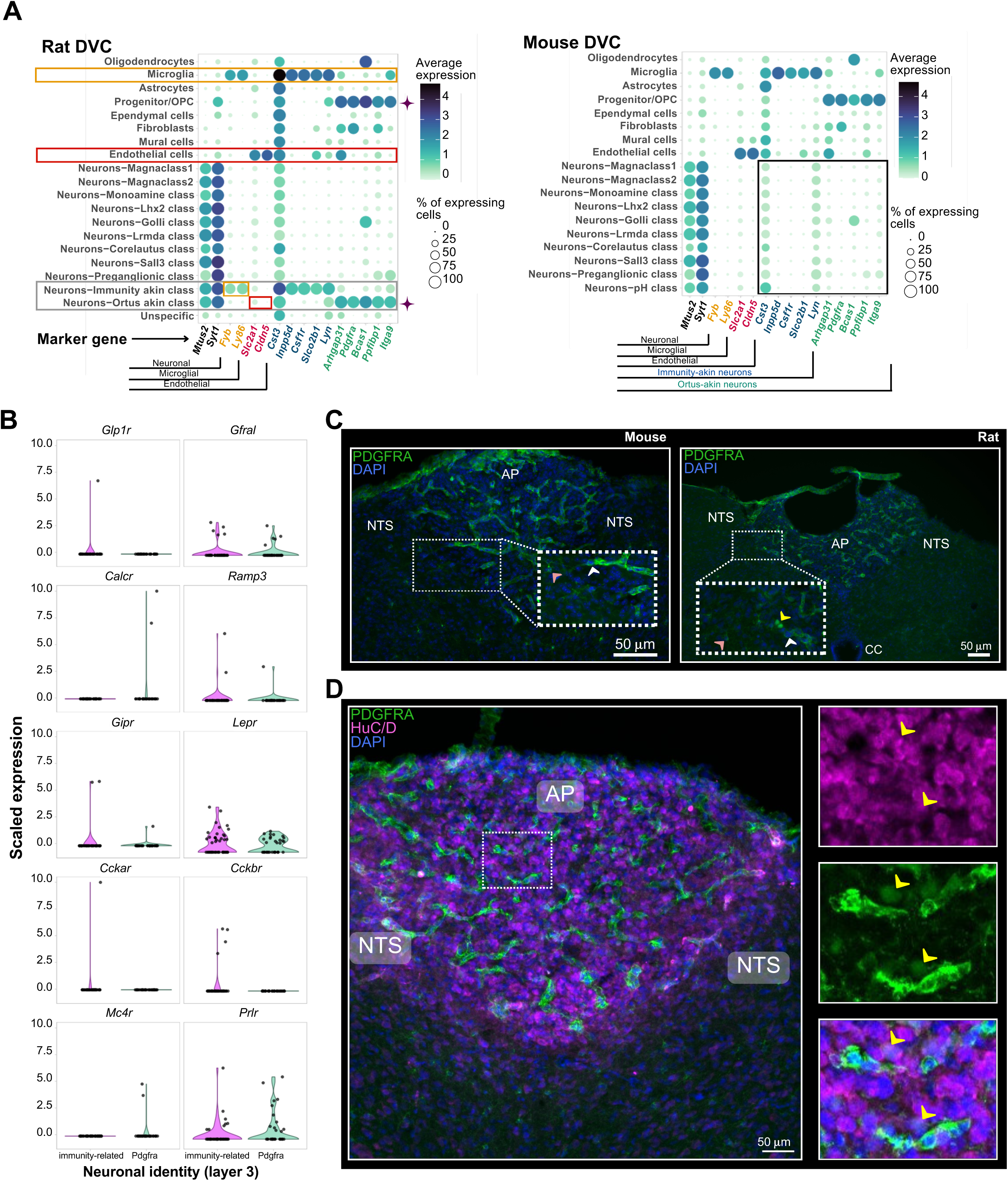
Two novel neuronal classes specific to the rat DVC. **A.** Balloon plots comparing the expression of the marker genes of the two novel neuronal classes found in rats (cells n=12,167): immunity-akin and the ortus-akin classes (framed in gray). Expression is shown across the layer-2 rat cell identities. Based on the expression of two DVC neuronal markers *Mtus2* and *Syt1*, we corroborated those cell classes contain neurons, and not microglial or vascular/endothelial cells (In orange and red, respectively). The immunity-akin class (n=48 neurons) shares high expression of *Cst3*, *Inpp5d*, *Csf1r*, *Slco2b1* and *Lyn* with microglial cells. Meanwhile, the ortus-akin class (n=34 neurons) shows higher overlap of gene expression markers (i.e. *Arhgap31*, *Pdgfra*, *Bcas1*, *Ppfibp1* and *Itga9*) with OPCs (highlighted with a ♦ symbol). Mouse cell identities (cells n=99,740) do not show these expression patterns (framed in black). Average expression was calculated using log-normalized counts. **B**. Violin plots of scaled expression of 10 genes (i.e. *Glp1r, Gfral, Calcr, Ramp3, Gipr, Lepr, Cckar, Cckbr,Mc4r* and *Prlr*) coding for metabolism-associated receptors in the Pdgfra and immunity-related rat neurons (Pdgfra neurons n=34; immunity-related neurons n=48). Pdgfra neurons are the only layer-3 identity within the ortus-akin class. An overlapping dotplot shows one dot per cell. **C**. Immunofluorescent detection of PDGFRA in mouse and rat DVC. Pink arrowheads point to cells with OPC morphology, white arrowheads point to cells with fibroblast morphology and the yellow arrowhead points to cells with neural morphology. **D.** Co-staining for PDGFRA and HuC/D in rat DVC with high magnification (right images) of area postrema PDGFRA-expressing cells . Yellow arrowheads indicate colocalization of HuC/D and PDGFRA. DVC=dorsal vagal complex; OPC=oligodendrocyte precursor cell; AP=area postrema; NTS=nucleus of the solitary tract; PDGFRA= Platelet-derived growth factor receptor A; CC=central canal

Other differences found between mouse and rat DVC expression includes increased expression of the leptin receptor in neurons as well as cholecystokinin (**Fig. S13**).

To construct an integrated rodent DVC cell atlas, we incorporated the labeled rat dataset into our hierarchy using treeArches[39] (**Fig.8**). The rat data was projected onto the mouse datasets (**Fig.8A**) and the 42 high resolution rat cell identities were appended to the tree (**Fig.8B; Fig.S13**). The final hierarchy has 123 labels of which 99 are high resolution according to our cellular granularity (i.e. layers 3 to 5) (**Fig.8B**). In addition it has 20 layer-2 identities and 7 cell identities that are novel from the rat dataset (**Fig.8B**).

**Figure 8:**
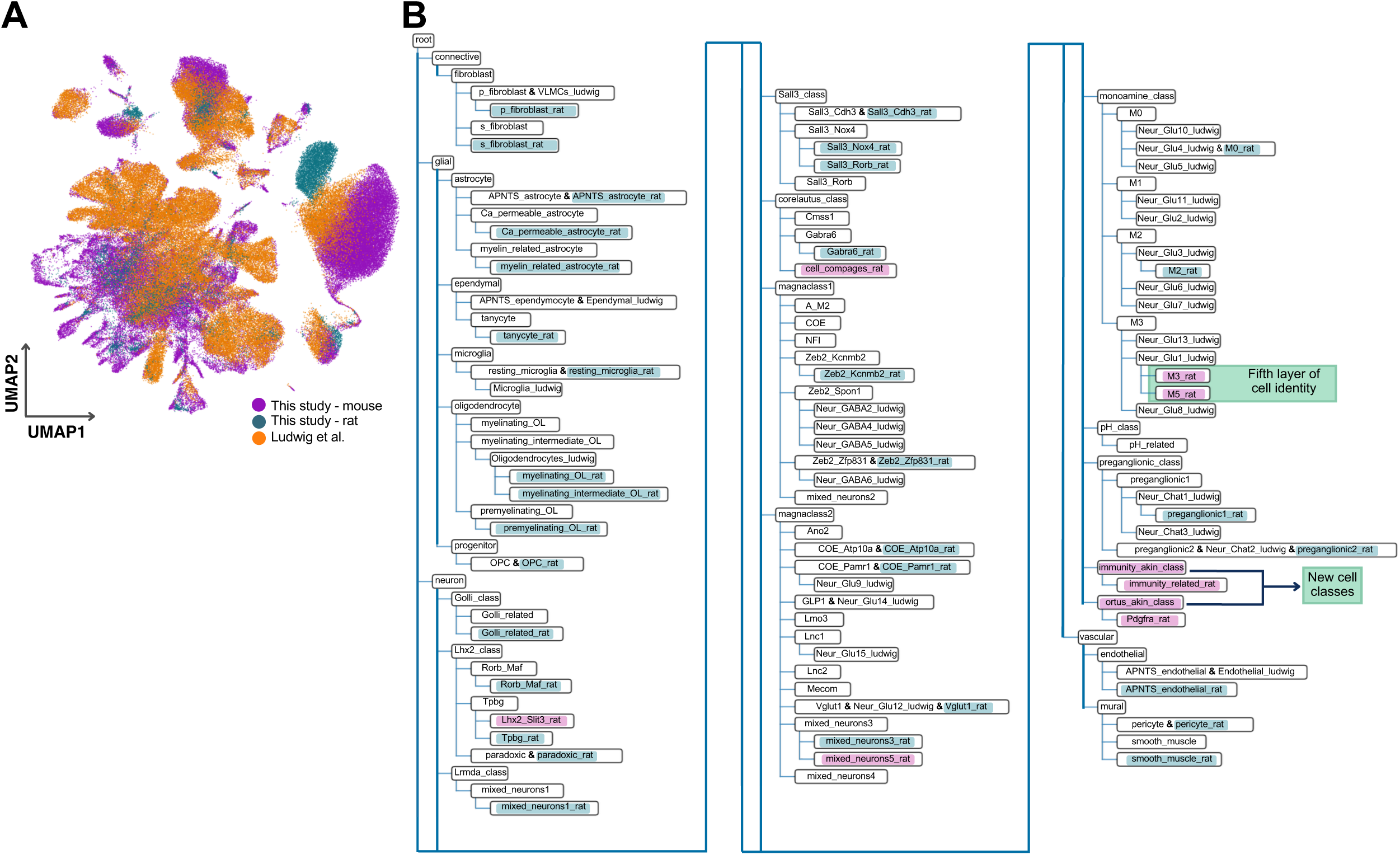
The rodent DVC cell hierarchy. **A.** UMAP plot of the integration between the murine hierarchy and our rat dataset using treeArches (cells n=184,035). **B.** Dendrogram of the harmonized hierarchy of our labeled rat dataset and the murine DVC hierarchy. We highlight the incorporated rat cell identities (blue and magenta). Those in magenta are the novel rat identities established by us. The immunity-akin and the ortus-akin classes were not found in the murine datasets. The M3 and M5 rat identities are subgroups of a Ludwig layer 4 cell identity, therefore yields a fifth layer within the monoamine neuronal class. UMAP=uniform manifold approximation and projection; DVC=dorsal vagal complex; APNTS=Area postrema and nucleus of the solitary tract; Lnc=long non-coding; OL=oligodendrocyte; OPC=oligodendrocyte precursor cell; Neur=neuronal; COE=collier/Olf1/EBF transcription factors

### 3.5. *Hcrt* and *Agrp* expression in the DVC

Since *Agrp*-expressing cells have been recently described within the DVC[67], we mapped this and other neuropeptide genes in DVC neurons. Surprisingly, we found *Hcrt* and *Agrp* mRNA in neurons in our dataset (**Fig.S14A**). We then performed *in-situ* hybridizations for these transcripts in both the mouse hypothalamus and DVC. While both *Hcrt* and *Agrp* were readily detectable in the hypothalamus, we failed to observe signal within the DVC (**Fig.S14B**). Since we also found expression of *Hcrt* and *Agrp* in neurons of the Dowsett and Ludwig datasets[2, 18] (**Fig.S14C**), we then mapped the sequenced reads from these transcripts and confirmed that most of them mapped to intronic and non-coding regions of these genes (**Fig.S14D**). Furthermore, our rat DVC dataset was pre-processed excluding introns from the reads mapping, and the *Hcrt* and *Agrp* expression levels observed is minimal (**Fig.S14E**), which is in agreement with the non-coding location of the reads in mouse DVC. These data explain absence of signal from our *in-situ* hybridizations and suggests DVC cells do not produce HCRT or AGRP neuropeptides.

### 3.6. Meal-related transcriptional programs in the murine DVC

As the DVC acts as a primary node for meal-related signals[68], we wanted to define which of our cell identities responded to meal consumption and thus performed differential gene expression analysis between DVC cell types from mice that were euthanized under fasting conditions, with ad libitum access to food or two hours post-refeeding following an overnight fast (**Fig.1A**; **Fig.9**). This analysis revealed widespread transcriptional changes across nearly every neural cell identity, the exceptions being corelautus, Golli class and pH classes (**Fig.9A**). Additionally, the size of the transcriptional changes were relatively small, with approximately 90% of the differentially expressed genes having increases or decreases between 0.5 and 1.0 log2 fold-change, with only ∼10% of the gene changes being ≥1 log2 fold-change between conditions (**Fig.9A**). Of the responding neurons, magnaclasses 1 and 2, which represent large populations of predominately inhibitory and excitatory neurons (**Fig.S2**), showed the greatest number of differentially expressed genes following refeeding for both comparisons refed versus fasting, and refed versus ad libitum food access (**Fig.9A**). To understand how refeeding transcriptionally alters inhibitory compared with excitatory DVC neurons, we compared differentially expressed genes between these two classes across conditions. Intriguingly, despite being in transcriptionally distinct classes, we found meal consumption altered many of the same genes (**Fig.9B-D**), suggesting these cells may be receiving similar postprandial signals which are propagated via these cells in unique manners. This is supported by the similar receptor expression profile of magnaclass 1 and 2 (**Fig.S15**). One notable exception is *Cckbr* which is upregulated in magnaclass 2 and downregulated in magnaclass 1 in refeeding when compared with ad libitum fed mice(**Table S12**). We also note that oligodendrocytes contained the highest number of differentially expressed genes in refeeding (**Fig.9A**). While the roles of oligodendrocytes in refeeding responses within the DVC have yet to be determined and the biologic significance of these changes are presently unclear, DVC oligodendrocytes also have robust transcriptional responses in response to fasting[18]; suggesting these cells are sensitive to both positive and negative energy balance associated-cues and may have important roles in regulating energy homeostasis.

**Figure 9.**
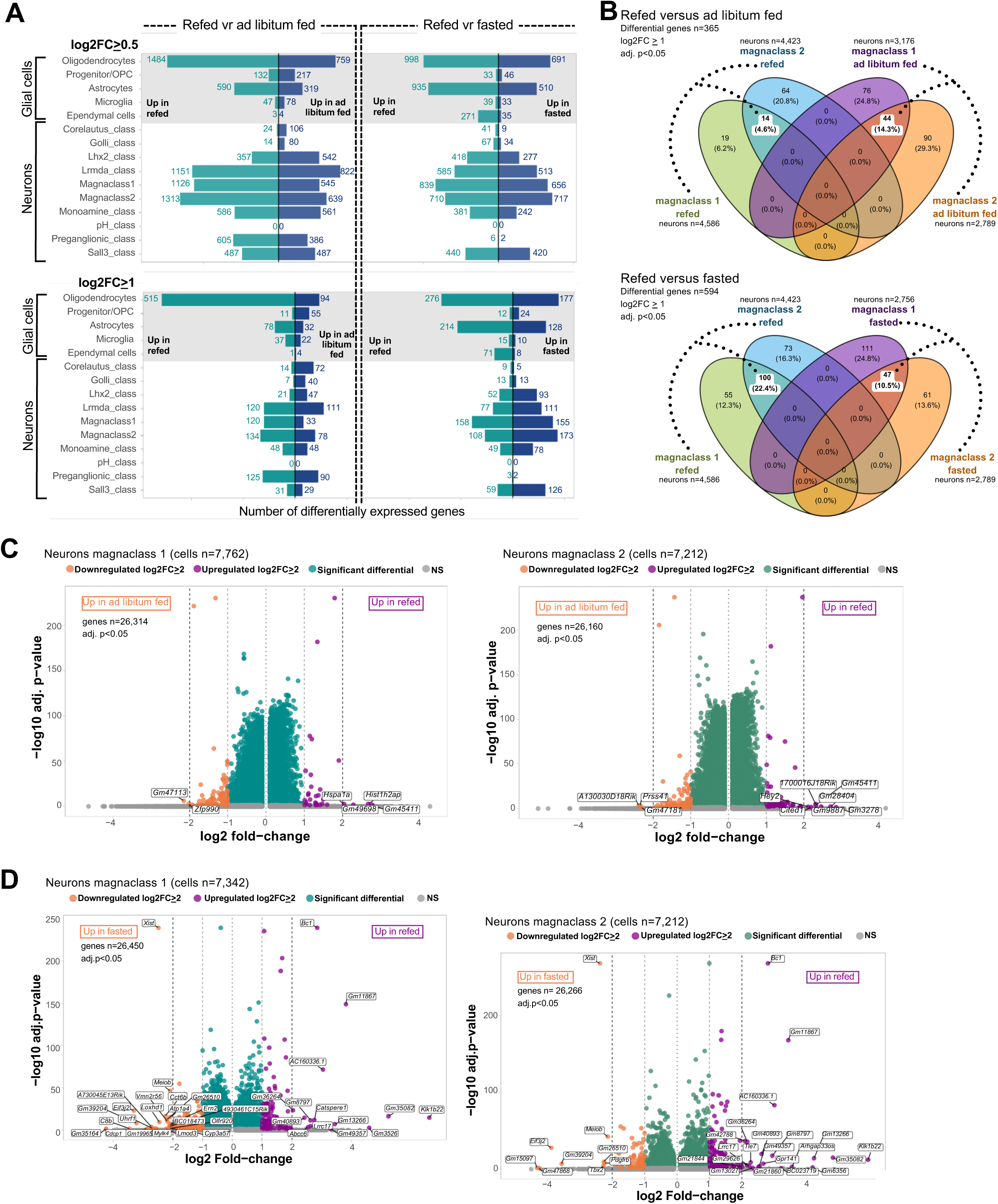
Meal-related transcriptional changes in the mouse DVC. **A.** Horizontal barplots of the number of differential genes between refed and ad libitum fed, and refed and fasted mice in glial cells and neurons per layer-2 cell identity (MAST algorithm on log-normalized counts; adj. p<0.05; refed neurons n=14,396; refed glial cells n=18,019; ad libitum fed neurons n= 9,885; ad libitum fed glial cells n=4,610; fasted neurons n=8,829; fasted glial cells n=12,496). **B.** Venn Diagrams of the differentially expressed genes in magnaclass 1 and magnaclass 2 neurons between refed and ad libitum fed, and between refed and fasted treatments (MAST algorithm on log-normalized counts). The number of overlapping log2FC≥1 upregulated genes between the two magnaclasses per treatment are highlighted. The treatment-induced changes in only one cell class are shown as non-overlapping. The percentage is based on the total differential genes surveyed per comparison. **C.** Volcano plots of the differential genes in neurons belonging to the magnaclass 1 and the magnaclass 2 between refed and ad libitum fed, and **D.** refed and fasted treatments (MAST algorithm on log-normalized counts; adj. p<0.05). Each point represents one gene. Only genes upregulated or downregulated with a log2FC≥2 are labeled. Since we considered a minimum log fold-change of 0.1 between treatments per cell group, those genes with very low variance (i.e. log fold-change<0.1) were excluded from the differential expression analysis, therefore a variable number of genes are shown per comparison per identity, in the volcano plots. DVC=dorsal vagal complex; vr=versus; OPC=oligodendrocyte precursor cell; FC=fold-change; adj.=adjusted; NS=non-significant

## 4. Discussion

The recent success of DVC-based therapies[13, 69] has moved DVC biology to the forefront of weight control therapies and there is now clear clinical impetus to fully define DVC cell types and their functions. Previous studies detailing DVC cell types describe a high degree of heterogeneity[2, 18, 19], however, these studies lacked an in-depth analysis of neurotransmitter profiles of neurons, any cross-species evaluation of DVC cell types, or an analysis of how meal consumption transcriptionally regulates these cells. We therefore developed *de novo* mouse and rat DVC cell atlases to address these points.

To more fully describe the cellular complexity within the DVC, we combined a pipeline of cell label transfer using brain existing published datasets and thorough curation of cell identities, which resulted in a 4-layer mouse cell hierarchy and a 5-layer rodent cell hierarchy of the DVC. Within this hierarchy we found several unique features of DVC glial cells. First, we found Ca^+^-permeable astrocytes in the DVC in which *Gfap* is almost exclusively expressed and they coexpress *Kcnj3* (GIRK1 potassium channel). These cells largely lack *Gria2*, consistent with findings in Bergmann glial cells in which colocalization of *Gria2* and GFAP was found to alter neuron-glial interactions[70]. The heterogeneity of astrocytes across, and even within, spatially distinct CNS sites is becoming increasingly apparent[71] and our finding of DVC-specific astrocyte transcriptional profiles further supports this. When we compared Ca^+^-permeable DVC astrocytes to hypothalamic astrocytes[43], we found the same expression pattern with minimal overlap between *Gria2* and *Gfap.* However, DVC astrocytes seem to have some unique properties when compared with hypothalamic, cortical and hippocampal astrocytes. Principally, DVC astrocytes express the GIRK1 channel, likely rendering them less excitable[72]. As astrocytes have well established roles in maintaining the blood brain barrier function[73], and the pattern of GFAP in the DVC is largely restricted to the boundary separating the AP and NTS, it is possible these DVC-specific astrocytes have specialized functions in regulating blood brain barrier permeability.

Additionally, while we identified hypothalamic fibroblasts which express high *Aldh1a1* similar to the DVC p-fibroblasts, we failed to find a group equivalent to s-fibroblasts. Other cell populations seem to be consistent to what has been found in other brain sites, like the intermediate oligodendrocytes that are found enriched in the hypothalamus and the corpus callosum in mice[55]. In summary, these findings show that while certain DVC glial cells share identities with glial cells from other CNS sites, there are also DVC-specific glial cells.

Within the mouse neural classes, we found overlap between *Cck* and *Th*-expressing cells. As these cells have previously been described as non-overlapping, this was unexpected[17]. Our *in-situ* hybridization confirmed the presence of *Cck*/*Th* positive cells within the NTS and AP. We believe the discrepancy may be due to detection methods as these cells were initially defined as non-overlapping using immunofluorescent detection of TH protein and a *Cck* genetic reporter[17]. Functional studies of *Cck-* and *Th*-expressing neurons within the NTS demonstrate these cells have somewhat distinct roles; while both reduce appetite, *Cck* neurons are strongly aversive whereas *Th*-labelled neurons are non-aversive[74]. While no functional characterization of *Cck* and *Th* neurons of the AP has been performed, we posit that as some *Th*-expressing AP neurons also express *Gfral* and *Casr*; both having potently aversive ligands (e.g. GDF15, deoxynivalenol)[16, 75]; they are likely to mediate aversive anorexia. The role of dual *Cck/Th* positive neurons remains to be determined.

While AGRP expression has long been considered to be hypothalamic-exclusive, a recent report describes *Agrp*-expressing cells within the mouse DVC[67], i n part from the detection of Agrp mRNA in scRNA-Seq data. Similarly, we also detected transcripts for both *Agrp* and *Hcrt* in our snRNA-Seq data but failed to detect any transcripts within the DVC by *in-situ* hybridizations. One possible explanation for the discrepancy between the snRNA-Seq data and our *in-situ* hybridizations data is that the snRNA-Seq mRNA reads for*Agrp* and *Hcrt* gene overwhelmingly mapped to non-coding regions, of the respective genes. Combined, these data suggests to us that neither AGRP or HCRT neuropeptides are produced by DVC cells in rodents.

We show the presence of neurons with seemingly mixed programs in the DVC, which we called ‘mixed neurons’. Similar groups have been described in the hypothalamus[26] suggesting that some neurons may have hybrid-functionalities. These mixed neurons share many transcriptomic markers expression with many other cell identities, and may require the use of other technologies to fully identify them anatomically in the brain.

To perform a cross-species analysis, we generated a rat-based DVC atlas and compared it with the mouse DVC. Our rat dataset was smaller than the mouse data, it is possible some of the of the new neuronal cell identities (e.g. M_sema, M4) may contain cells belonging to multiple cell types, as treeArches did not appended these identities to the tree. Despite this, we corroborated two new cell types in the rat DVC that are not present in mice the *Pdgfa*-expressing (ortus-akin class) and the immunity-related neurons*. Pdgfra* is a well described marker gene for OPCs and fibroblasts within the central nervous system[66] but, to our knowledge, has not been associated with neurons. Notably, although the PDGFRA immuno-signal in HuC/D+ neurons is clearly discernible, it is less intense than in OPCs and fibroblasts, in agreement with expression levels from our snRNA-Seq data. Since it was previously demonstrated that OPCs can give rise to neurons in the adult brain[76] and there is evidence of neurogenesis in the AP following amylin administration in rats[77], it is possible that these PDGFRA+/HuCD+ cells in the AP represent a transient stage of PDGFRA+ OPCs differentiating into neurons.

Regardless of their maturity, we speculate that the Pdgfra neurons (ortus-akin class) have roles in appetite regulation as they express leptin receptor and *Gfral*. Interestingly, while multiple studies have failed to detect *Lepr* via Cre-based reporters[62, 78] or leptin-induced phosphorylated STAT3 in the AP of mice[79], *Lepr* and leptin-induced phosphorylated STAT3 are detectable within the rat AP[80–82]. We believe one explanation for this species discrepancy in *Lepr* expression is the absence of the ortus class neurons in mice. Future studies are needed to characterize these cells and determine whether their presence is found in other mammals including humans.

DVC-targeting obesity therapeutics, including amylin and GLP1 mimetics, produce clinically relevant weight loss but are associated with multiple gastrointestinal adverse events and there is a need to develop new obesity therapies that can reduce appetite without unwanted gastrointestinal side effects[83]. Indeed, the DVC cells that control appetite suppression and nausea are separable[11, 74], making targeting such cells an attractive target for pharmacotherapy. As meal consumption is generally not associated with gastrointestinal distress, we sought to identify refeeding responsive cells within the mouse DVC. From this analysis, we found nearly all neural cell identities are transcriptionally altered by meal consumption. We also found the magnitude of transcriptional changes was relatively small, regardless of the cell class examined. While relatively small, the scale of the transcriptional changes in response to refeeding are similar to many of the gene expression changes observed in *Agrp*-neurons following 16 hours food deprivation[43]. Regardless of whether this is a technical limitation of snRNA-Seq methods or accurately reflects the scale of transcriptional changes in the DVC, utilizing transcriptional responses to refeeding to identify neural classes as targets for pharmacotherapy is unlikely to yield specific and targetable pathways.

In general, one limitation in our study is that it is based on nuclear RNA which may reflect a fraction of biological changes. Using nuclear isolation facilitates generating to have single-neuron data[84] since obtaining viable whole neurons (including all their processes) is very challenging. Previous studies on the relationship between nuclear and cellular RNA content in brain show that nuclear sequencing reads may map to intronic regions of genes in higher proportion[85]. Although this does not appear to impact gene expression analysis for neurons in the current pipelines like the ones we used for our mouse data[85], it is possible that some biological features such as microglial activation state, are not fully recovered using nuclear RNA[86]. Although we performed sub-clustering in the mouse microgia and assessment of the resulting marker genes for each group, we did not find evidence of activation possibly due to this matter. Separately, for meal consumption, we were unable to analyze responses in individual mice as pooling DVCs from individual mice was the most amenable option to maximize nuclei for in our pipeline. As we pooled DVCs from both sexes, this limited our ability to detect sexual dimorphic cell class proportions and gene expression levels.

## 5. Conclusion

As the amount of snRNA-Seq data performed on the DVC continues to grow, there is a need for a common, scalable label set that can be applied to new studies and integrate new sequencing data. To develop such a label set, we started with a *de novo* DVC atlas, we generated three layers of cellular granularity for both neural and non-neuronal cell types. Then, using the treeArches pipeline[39] we harmonized our snRNA-Seq labels with the Ludwig dataset[2] and increased our cellular granularity from three to four layers of resolution. Manually curating and harmonizing our dataset enhanced the ability of our cell atlas to capture the complexity of the DVC neurons since not all groups seem to contain easily delineable subgroups of cells. For example, eleven of the labels incorporated into our cell hierarchy in this pipeline belong to the monoaminergic neuronal class, which seem to encompass multiple subgroups of specialized cells. We further constructed the rodent DVC cell atlas which resulted in a fifth layer of cell identity to the monoaminergic M3 neurons. Although the number of identities increases as layers are more specific, this gives this unified atlas greater adaptability to different research needs. Additionally, our labels can be accurately harmonized with new databases regardless of size and new DVC snRNA-Seq data can be added to the tree to generate new branches. Thus, our unified rodent DVC atlas is highly amenable to future research elucidating new biologic and therapeutic insights into DVC cell types.

## Abbreviations

3v: third ventricle
adj.: adjusted
AMPAr: alpha-amino-3-hydroxy-5-methyl-4-isoxazolepropionic acid receptor
AP: area postrema
APNTS: Area postrema and nucleus of the solitary tract
ARH: arcuate nucleus of the hypothalamus
BAM: binary alignment map
CC: central canal
COE: collier/Olf1/EBF transcription factor
diH_2_O: distilled water
DMV: dorsal motor nucleus of the vagus
DVC: dorsal vagal complex
FACS: fluorescence-activated cell sorting
FC: fold-change
GABA: gamma-aminobutyric acid
GDF15: growth differentiation factor 15
GIRK1: G protein-coupled inwardly rectifying potassium channel 1
GLP1R: glucagon-like peptide 1 receptor
GFRAL: GDNF family receptor alpha-like
IR: immunoreactivity
LH: lateral hypothalamus
Lnc: long non-coding
ME: median eminence
Neur: neuronal
NG: *Ng2* glial cell
NS: non-significant
NTS: nucleus of the solitary tract
OL: oligodendrocyte
OPC: oligodendrocyte precursor cell
PC: principal component
PDGFRA: Platelet-derived growth factor receptor A
Res.: resolution
RNA-seq: RNA sequencing
smc: smooth muscle cells
snRNA-seq: single-nuclei RNA sequencing
UMAP: uniform manifold approximation and projection
UMI: unique molecular identifiers
v: version
VGLUT2: vesicular glutamate transporter-2
vr: versus

## 6. Acknowledgements

We would like to thank authors of the Ludwig and the Dowsett publications for making publicly available their datasets. We thank M. Myers Jr. for the valuable help provided; S. Hebert and S. Bailey for their insights and suggestions in regards to the bioinformatic analysis; members of the Kokoeva lab for their insights on the *in-situ* hybridization and immunohistochemistry pipelines; S. Feng and M. Fu for technical assistance for images acquisition at the Molecular Imaging Platform at the RI-MUHC, and A. Duensing for assistance with relevant file transfer.

## 7. Author contributions

Conceptualization: CH and PS

Methodology: CH, AB, LM, MK and PS

Sample collection: AB and PS

Software: CH, LM and HJM

Data analysis: CH, LM, MK and PS

Data curation: CH and HJM

Validation of findings: CH, FS, HJM and PS

Imaging: CH and PS

Visualization: CH, MK and PS

Supervision: CH and PS

Writing—original draft: CH and PS —review and editing: CH, AB, LM, FS, HJM, MK and PS

Funding acquisition: MK and PS

## 10. Data statement

All data needed to evaluate the conclusions in the paper are present in the paper and/or the Supplementary Materials. FASTQ files have been deposited in the NCBI Sequence Read Archive (Accession: PRJNA1161292). In addition, our murine and rat datasets have been made available for visualization and analysis through the Broad Institute Single Cell Portal (https://singlecell.broadinstitute.org/single_cell/study/SCP2773). The labels from our cell atlas can be transferred to new datasets through treeArches using the available hosted Jupyter Notebook in Google Colab for the mouse data (https://colab.research.google.com/drive/1v983gcQxIwBN-4lGx18JXPweZoKqMbwX#scrollTo=SJTZppZWa-63), and for the rodent data (https://colab.research.google.com/drive/10tAmfRosMscuBak6wHX80TgdEBsHeWQB#scrollTo=941ab5c1-8910-4e67-a632-4d18601d45b4). Original code used for constructing the murine and rodent cell hierarchies and predicting labels using treeArches are available in the GitHub repository https://github.com/LabSabatini/DVC_cell_atlas.

## Supplementary Material

<u>Figure S1: DVC-centric dissection protocol</u>

**A.** Following euthanasia, the brain is removed from the mouse and briefly rinsed with PBS. To better visualize the brainstem, the cerebellum is removed with fine forceps. The brain is then placed within a mouse brain matrix with 1mm spacing between razor guides. In this example blades will be placed in the 12th (cut 1) and 14th (cut 2) razor guide (these sites are shown). Using fresh razors make two clean cuts in a single, smooth motion. The bottom showns the reference images of the mouse brain (from the Allen Brain Institute reference 3D mouse atlas) visualizing the location of brainstem and the AP/NTS. In these reference images, the cerebellum is not visualized. **B.** Resultant brainstem tissue from the first two cuts is then further cut twice more (cuts 3 &4) to keep the middle of the tissue where the DVC is located. This tissue is cut once more (cut 5) to generate a DVC-centric dissection. Reference images of the mouse brain (Coronal sections from the Allen Brain Institute reference atlas) displaying the rostral and caudal sides of tissue collected from cuts 1&2 and the location of cuts 3-5, are shown.

<u>Figure S2: Mapping of *Gad2* and VGLUT2 in neurons</u>

UMAP plot of the murine neurons (n=48,131) showing scaled log-normalized expression of **A.** *Gad2*, a gene related to GABA synthesis and transmission, **B**. and VGLUT2, a glutamate transporter. Cells with higher expression of *Gad2* belong to the neuronal magnaclass1, while cells with higher expression of VGLUT2 (*Slc17a6*) to the magnaclass2 (both highlighted in yellow in adjacent plots). However, the expression is not mutually exclusive among both classes, nor every neuron belonging to each class express the respective transcript.

UMAP=uniform manifold approximation and projection; GABA=gamma-aminobutyric acid; VGLUT2=vesicular glutamate transporter-2

<u>Figure S3: Anatomical mapping of the DVC mouse layer-3 identities</u>

Schematic diagram of the spatial localization for the 35 neuronal identities at layer 3 of resolution in the mouse DVC. Coronal sections from the mouse brain in the Allen Brain Atlas were used to manually map the top marker genes of each (MAST algorithm on log-normalized counts; adj. p<0.05) to the DVC.

AP=Area Postrema; NTS= Nucleus of the solitary trac; DMV= Dorsal motor nucleus of the vagus; CC= central canal; 12N= hypoglossal nucleus; Cu=Cuneate nucleus.

<u>Figure S4: Mapping of DVC populations in a murine hypothalamic dataset</u>

UMAP plots of a randomly selected subset of the HypoMap database (cells n=76,260; samples n=32) labeled **A**. by sample and **B**. by original curated cell class. **C.** Balloon plot showing average expression of the top 5 upregulated gene markers of each neuronal DVC class (MAST algorithm; adj. p<0.05) in the neurons of the HypoMap subset (n=45,679). We selected the neuronal curated class cells from our initial HypoMap subset and reprocessed them, including clustering through *k*-nearest neighbors algorithm using the first 30 PCs and Harmony reduction embeddings. Resulting clusters at resolution 0.5 are shown. Average expression in the HypoMap neurons was calculated using log-normalized counts. **D.** Set of UMAP plots showing scaled expression of three genes (i.e. *Gfap*, *Gria2* and *Kcnj3*) which have a specific pattern of expression in murine DVC Ca^+^-permeable astrocytes. The astrocytic population is circled in purple (astrocytes n=7,629) and while *Gfap* and *Gria2* expression is in agreement to what we observe in Ca^+^-permeable astrocytes, in the hypothalamic astrocytes *Kcnj3* is not expressed. E. Set of UMAP plots of expression of *Aldh1a1* and *Stk39* which are markers of p-fibroblasts and s-fibroblasts in the DVC. The hypothalamic fibroblasts are circled in magenta (fibroblasts n=300).

DVC=dorsal vagal complex; UMAP=uniform manifold approximation and projection; AMPAr=α-amino-3-hydroxy-5-methyl-4-isoxazolepropionic acid receptor; GIRK1=G protein-coupled inwardly rectifying potassium channel 1; OPC=Oligodendrocyte precursor cell; NG=*Ng2* glial cell

<u>Figure S5: Mapping of DVC Ca^+^-permeable astrocytes in other murine brain sites</u>

UMAP plots of a cortical/hippocampal astrocytes dataset by **A.** sample, **B**. brain area and **C**. original astrocyte type (cell n=1,811). Set of UMAP plots showing scaled expression of three genes (i.e. *Gfap, Gria2* and *Kcnj3*) which have a specific pattern of expression in murine DVC Ca^+^-permeable astrocytes in **D.** cortical/hippocampal, **E.** forebrain, **F.** pons and **G**. spinal cord astrocytes. There is some overlap in *Gfap* and *Gria2* expression in cortex, hippocampus and forebrain, with almost no expression of *Kcnj3*. This is different from what we observe in DVC Ca^+^-permeable astrocytes. In pons and the spinal cord, there is some expression of *Kcnj3*, but the *Gfap* and *Kcnj3* co-expression is rare.

DVC=dorsal vagal complex; UMAP=uniform manifold approximation and projection

<u>Figure S6: Trajectory inference of the oligodendrocyte lineage</u>

Dotplot of the two principal components obtained through dimensionality reduction using the main 50 principal components analysis on log-normalized gene expression data in oligodendrocytes (cells n=32,129) **A**. merged and **B**. by sample, and on Harmony embeddings in oligodendrocytes **C.** merged and **D**. by sample. Cellular trajectory was inferred using SCORPIUS also on normalized gene expression data (**A**. and **B.)** and on the Harmony embeddings (**C.** and **D**.), and drawn across the data points (black line). Each data point represents one cell.

OL=oligodendrocyte

<u>Figure S7: Murine DVC clusters</u>

**A**. UMAP plot of the mouse DVC clusters at resolution 1 (cells n=99,740). Cells from mice exposed to 5 food and aversion-related treatments were integrated using the Harmony algorithm in R and the resulting embeddings were used to calculate the clusters and UMAP projections. We subset and re-clustered three clusters containing oligodendrocytes, vascular cells and fibroblasts, respectively (highlighted). **B.** Violin plots of the two types of fibroblasts in cluster 26, distinguished by *Skt39* (stress-response ‘s-fibroblasts’) and *Aldh1a1* (perivascular ‘p-fibroblasts’) expression. Each data point represents one cell (s-fibroblasts n=244; p-fibroblasts n=559). **C**. UMAP plot of subclusters from clusters 23 (oligodendrocytes) and **D.** 27 (vascular cells) using the Harmony embeddings. Gene expression markers for cell populations were mapped (**Table S5**) and the smallest resolution that allowed to separate them was further used to label the cells. Cells in subclusters 1 and 2 from cluster 23 were labeled ‘premyelinating_OL’ whilst subclusters 0 and 3 were labeled ‘myelinating_intermediate_OL’. In cluster 27, subcluster 7 was labeled ‘pericyte’, subcluster 4 was labeled ‘smooth_muscle’ and the rest of the clusters were considered to be endothelial cells.

DVC=dorsal vagal complex; Res.=resolution; UMAP=uniform manifold approximation and projection; OL=oligodendrocyte; smc=smooth muscle cell.

<u>Figure S8: Mouse microglial activation states</u>

**A.** UMAP plot showing the mapping in the mouse DVC microglia (n=2,787).of monocyte (*Emilin2* and *Gda*) and microglial (*P2ry12* and *Fcrls*) known differential markers, corroborating the absence of monocytes in our dataset**. B.** UMAP plot of the microglia from the mouse DVC showing scaled log-normalized expression of *Cd5*, *Zbp1*, *Cxcl9* and *Cxcl10*. Microglia was subset and re-processed using the harmony integration embeddings to obtain the UMAP coordinates. **C.** Sub-clusters at resolution 0.2 of these microglial cells was performed and the **D.** top marker genes are shown in a balloon plot with average log-normalized counts (MAST algorithm; adj. p<0.05) along with astrocytes and oligodendrocytes (microglia n=2,787, oligodendrocytes n= 32,129, astrocytes n=11,693). Cluster C3 shares high expression of markers with oligodendrocytes (in purple) while cluster C5 shares markers with astrocytes (in red). Microglial markers are highlighted in orange.

UMAP=uniform manifold approximation and projection; DVC=dorsal vagal complex

<u>Figure S9: Mixed neurons sub-clustering</u>

UMAP plots with the mixed neurons of the mouse DVC (mixed neurons n=15,190) showing their **A.** original layer-3 identity and **B.** final sub-cluster. **C**. Heatmap of the top 5 marker genes (MAST algorithm; adj. p<0.05) expression for each sub-cluster in all neurons (i.e. mixed neurons and all other neuronal identities, neurons n= 48,131). Each column is one cell and each row is one marker gene. Some marker genes were duplicated for more than one sub-cluster (e.g. *Nrxn3* is marker of sub-clusters 2, 3, 8 and 10; *Pcdh9* is marker of sub-clusters 6 and 8) therefore these genes are only shown once in the plot and highlighted in red.

UMAP=uniform manifold approximation and projection; DVC=dorsal vagal complex

<u>Figure S10: Initial murine cell hierarchy using our samples</u>

**A**. UMAP plot of the integration of our samples (based on the five mouse treatments administered to mice prior to euthanasia; see Methods) using treeArches. There were two cohorts of refed mice. **B**. UMAP plot of the fifty layer-3 cell identities obtained using the treeArches pipeline. **C**. Dendrogram showing the initial hierarchy of the three layers of cell identity established by us in this study.

UMAP=uniform manifold approximation and projection; inj.=injected; APNTS=Area postrema and nucleus of the solitary tract; Lnc=long non-coding; OL=oligodendrocyte; OPC=oligodendrocyte precursor cell; COE=collier/Olf1/EBF transcription factors

<u>Figure S11: Incorporation of the Ludwig dataset to our initial hierarchy using treeArches</u>

**A.** Barplot of log-normalized expression of OPC markers (i.e. *Pdgfra*, *Cspg4*, *C1ql1*) and premyelinating OL markers (i.e. *Bmp4*, *Fyn*) in the OPCs-labeled cells from the Ludwig dataset (total cells n=1,770; cells expressing >1 marker gene n=1,104). Cells labeled as OPCs in the Ludwig dataset are a mix of OPC and premyelinating OL. Some cells lack expression of the five marker genes used in this plot, they are represented on the left side with an empty bar and those expressing *Fyn*/*Bmp4* but not expressing the OPC markers are likely premyelinating OLs. Cells were ordered using hierarchical clustering based on these five gene expression data through the Ward’s method in R. We only indicate the ‘likely cell identity’ for cells with expression of the five marker genes. **B**. UMAP showing the manual mapping of Ludwig and Dowsett neuronal clusters and identities in neurons from this study. We used the gene expression markers for all clusters from Ludwig and Dowsett to identify which of our neurons shared identity with those described in both publications. Only groups and clusters successfully mapped manually are shown. The Ludwig identities are in black, Dowsett identities are in blue and our identities are in purple. Most of neuronal clusters in both publicly available datasets, are subgroups of our cell identities. OPC=oligodendrocyte precursor cell; OL=oligodendrocyte; UMAP=uniform manifold approximation and projection; APNTS=Area postrema and nucleus of the solitary tract; Lnc=long non-coding; COE=collier/Olf1/EBF transcription factors

<u>Figure S12: Rat DVC neurons</u>

**A.** Labeled UMAP plot with the neuronal layer 3 cell identities in our rat dataset (neurons n=4,721). Neurons were subset and re-processed to obtain the UMAP coordinates. **B.** Violin plots of scaled expression of 12 genes (i.e. *Gad2, Gad1, Slc6a5, Slc17a7, Slc17a6, Th, Ddc, Nos1, Sst, Npy, Cck* and *Slc5a7*) related to transmission/release of GABA, glutamate, monoamines, nitric oxide, somatostatin, neuropeptide-Y, cholecystokinin and acetylcholine in the Pdgfra and immunity-related rat neurons (Pdgfra neurons n=34; immunity-related neurons n=48). An overlapping dotplot shows one data point representing each cell. DVC=dorsal vagal complex; UMAP=uniform manifold approximation and projection; COE=collier/Olf1/EBF transcription factors; Lnc=long non-coding; GABA=gamma-aminobutyric acid

<u>Figure S13: Receptors and neuropeptides expression in mouse and rat neurons</u>

Balloon plot of average nuclear mRNA for a selected list of receptors and neuropeptides related to metabolism in mouse and rat neurons (rat n= 4,721, mouse n=48,131). Expression values represent average log-normalized counts for each gene. All cell identities at layer 3 for each species are shown.

<u>Figure S14: Dendrogram after incorporation of the rat dataset to the murine hierarchy Dendrogram (i.e. tree) resulting from incorporating our rat dataset (cells n=12,167) to the existing murine DVC cell hierarchy using treeArches. The rat cell populations in red that did not show similarity with the previously existing murine identities are appended at the bottom by treeArches (shown in red). DVC=dorsal vagal complex</u>

<u>Figure S15: Expression of *Agrp* and *Hcrt* in DVC neurons</u>

**A.** UMAP plots of the neurons (n=48,131) from our murine DVC dataset showing scaled expression of *Agrp* and *Hcrt*. **B**. *In-situ* hybridization images of *Agrp* and *Hcrt* in the DVC show no exonic mRNA. We included tissue sections of the arcuate nucleus of the hypothalamus and in the lateral hypothalamus as controls, and *Agrp* and *Hcrt* were present in these sites, respectively. For each reaction, tissue from the same mouse was used through two different sections, one including the DVC and one including the hypothalamic region. **C.** UMAP plots of the Dowsett (n=8,288) and the Ludwig datasets neurons (n=49,392) showing scaled expression of *Agrp* and *Hcrt*. **D.** Genome visualization of reads coverage in the *Agrp* and *Hcrt* regions of the *Mus musculus* refdata-gex-mm10-2020-A reference from two of our mouse sample sequencing replicates. Resulting BAM files from the original murine DVC FASTQ files were obtained after processing. We visualized the reads from these BAM files that mapped to the *Agrp* and *Hcrt* genes using IGV v2.16.1[87]. The coverage of these genes (*Agrp* region: chr8:105,566,180–105,582,268; *Hcrt* region: chr11:100,761,693–100,762,931) are shown in gray. The majority of the reads mapped to the intronic regions of *Agrp*, and to the last part of the 3’-end of exon 2 and the 3’-untranslated region of *Hcrt*. **E.** UMAP plot of rat DVC neurons (n=4,721) showing scaled expression of *Agrp* and *Hcrt*. For all UMAP projections, neurons from each dataset were subset based on our labeling for our murine and rat datasets, the treeArches resulting labeling for the Dowsett dataset, and the original labeling from Ludwig and collaborators. Then the neurons were processed using Seurat[23] and integrated using Harmony[25], to yield the final UMAP embeddings.

DVC=dorsal vagal complex; UMAP=uniform manifold approximation and projection; AP=area postrema; NTS=nucleus of the solitary tract; ARH=arcuate nucleus of the hypothalamus; 3v=third ventricle; ME=median eminence; LH=lateral hypothalamus. BAM=binary alignment map

<u>Figure S16: Expression of metabolism-associated receptors in the DVC magnaclasses</u>

Violin plots of scaled expression of 10 genes (i.e. *Glp1r, Gfral, Calcr, Ramp3, Gipr, Lepr, Cckar, Cckbr, Mc4r* and *Prlr*) coding for metabolism-associated receptors in the magnaclass1 and magnaclass2 neurons of the mouse DVC in fasted (magnaclass1 n=2,756; magnaclass2 n=2,789) and refed (magnaclass1 n=4,586; magnaclass2 n=4,423). DVC=dorsal vagal complex

<u>Table S1: Filtering of our and Dowsett datasets during processing</u>

Compilation of statistics and cells processed for the initial high quality cell filtering per sample.

<u>Table S2: Cluster filtering of our and Dowsett datasets during processing</u>

Compilation of low quality cluster analysis statistics to filter high quality clusters per sample.

<u>Table S3: Singlets retained per dataset after processing</u>

Number of doublets and singlets identified by DoubletFinder[24] after initial filtering of our and Dowset dataset per sample. Doublets were removed from the datasets before sample integration with Harmony[25] (this step was not performed in the rat dataset as it was only one sample, n=2 rat DVC).

<u>Table S4: Murine brain databases used to map our unlabeled mouse dataset</u>

We used four databases included in the scRNAseq v2.16.0[26–28, 31, 55] R package and the celldex v1.12.0 built-in mouse RNA-seq reference (which we called ‘SingleR’ database) using SingleR [30] function to transfer labels to our dataset in R[22]. The name, brain area and reference in which each database was made public are specified.

<u>Table S5: Literature-derived markers per central nervous system cell identity</u>

We thoroughly searched in the literature for established markers at the mRNA and/or protein level. Some markers were found to be described for more than one cell population which are specified in columns ‘notes’ and ‘notes_2’. After initial labeling of our murine dataset with the databases as described, we proceeded to obtain the average expression (on log-normalized counts) of each gene coding for the markers for the labeled cells (i.e. astrocyte, endothelial, microglia, neuron and oligodendrocyte) and unlabeled cells (i.e. unknown). We also calculated the average expression for each cluster in our dataset, established during our processing pipeline, at resolution 1.0.

<u>Table S6: Description of our DVC layer-3 neuronal cell identities established in mice</u>

Neurons were subset and reprocessed before labeling at layer-3 granularity. The MAST algorithm[32] was used through Seurat v5[23]on scaled data with a log fold-change threshold of 0.25 and detection threshold on ≥40% of cells on each neuronal cluster at resolution 1.0. We searched the resulting gene expression markers in the literature and compiled them. We then based the names for our cell identities on these genes.

<u>Table S7: Literature-derived markers per central nervous system cell identity</u>

Breakdown of the cell identities (except neurons at layer 3 of cellular resolution) we established and corroborating using a combination of markers surveyed in Table S2 and additional markers established using the FindConservedMarkers function from Seurat v5[23] package, indicating a cell state derived from the literature. For example, ‘s_fibroblasts’ were named after finding high expression of *Stk39*, a stress-related marker[56].

<u>Table S8: Expression of the murine DVC identity markers by rat DVC clusters</u>

Average log-normalized expression of the top 5 gene expression markers of each of our murine layer-3 cell identities (except duplicated genes n=37, and mouse specific genes n=27; total genes included=186) by rat DVC cluster at resolution 2.0.

<u>Table S9: Description of our DVC layer-3 neuronal cell identities established in rats</u>

Rat neurons were subset and reprocessed before labeling at layers 2 and 3 of granularity. We used neuronal class gene expression markers to identify clusters at resolution 2.0 that belonged to each neuronal cell class. These cell classes were further subclustered and markers from Table S6 and UMAP projections were used to label each population in the dataset.

<u>Table S10: Labeled cells in our murine dataset Number of cell per treatment per identity layer 2.</u>

<u>Table S11: Expression of neurotransmitter-related genes in our murine neurons</u>

Expression of genes coding for neurotransmitter synthesis enzymes TH, DBH, TPH, GABA, NOS, prohormone SST and primary neurotransmitter transporters VMAT2, VGLUT1, VGLUT2, GLYT2 per neuron in our dataset. Expression and glutamate and GABA release are specified as binary variables (i.e. 0=no expression; 1=expression) in the green and blue columns, respectively. The percentile of the log-normalized expression is shown. ∼55% of the neurons display expression of genes related to the synthesis/release of >1 primary neurotransmitter.

<u>Table S12: Receptor expression in magnaclasses 1 and 2 between meal-related treatments in mice</u>

Differential expression analysis (MAST algorithm)[32] results for genes coding for metabolism related receptors between meal-related treatments (i.e. refed versus ad libitum fed, and refed versus overnight fasted) and per neuronal magnaclass. If a receptor is missing for any given comparison for any of the magnaclasses, the log fold-change of that gene between both treatments in that neuronal class was <0.1, therefore it is not reported (see Methods).

## References

[1] L. Gautron, J.K. Elmquist, K.W. Williams, Neural control of energy balance: translating circuits to therapies, Cell. 161 (2015) 133–45. 10.1016/j.cell.2015.02.023.

[2] M.Q. Ludwig, W. Cheng, D. Gordian, J. Lee, S.J. Paulsen, S.N. Hansen, et al., A genetic map of the mouse dorsal vagal complex and its role in obesity, Nat Metab. 3 (2021) 530–45. 10.1038/s42255-021-00363-1.

[3] L. Bai, S. Mesgarzadeh, K.S. Ramesh, E.L. Huey, Y. Liu, L.A. Gray, et al., Genetic Identification of Vagal Sensory Neurons That Control Feeding, Cell. 179 (2019) 1129–43 e23. 10.1016/j.cell.2019.10.031.

[4] W. Han, L.A. Tellez, M.H. Perkins, I.O. Perez, T. Qu, J. Ferreira, et al., A Neural Circuit for Gut-Induced Reward, Cell. 175 (2018) 665–78 e23. 10.1016/j.cell.2018.08.049.

[5] E.K. Williams, R.B. Chang, D.E. Strochlic, B.D. Umans, B.B. Lowell, S.D. Liberles, Sensory Neurons that Detect Stretch and Nutrients in the Digestive System, Cell. 166 (2016) 209–21. 10.1016/j.cell.2016.05.011.

[6] M.W. Schwartz, Central nervous system regulation of food intake, Obesity (Silver Spring). 14 Suppl 1 (2006) 1S–8S. 10.1038/oby.2006.275.

[7] W. Cheng, E. Ndoka, J.N. Maung, W. Pan, A.C. Rupp, C.J. Rhodes, et al., NTS Prlh overcomes orexigenic stimuli and ameliorates dietary and genetic forms of obesity, Nat Commun. 12 (2021) 5175. 10.1038/s41467-021-25525-3.

[8] S.Z. Yanovski, J.A. Yanovski, Long-term drug treatment for obesity: a systematic and clinical review, JAMA. 311 (2014) 74–86. 10.1001/jama.2013.281361.

[9] T. Borner, C.G. Liberini, T.A. Lutz, T. Riediger, Brainstem GLP-1 signalling contributes to cancer anorexia-cachexia syndrome in the rat, Neuropharmacology. 131 (2018) 282–90. 10.1016/j.neuropharm.2017.12.024.

[10] S.M. Advani, P.G. Advani, H.M. VonVille, S.H. Jafri, Pharmacological management of cachexia in adult cancer patients: a systematic review of clinical trials, BMC Cancer. 18 (2018) 1174. 10.1186/s12885-018-5080-4.

[11] K.P. Huang, A.A. Acosta, M.Y. Ghidewon, A.D. McKnight, M.S. Almeida, N.T. Nyema, et al., Dissociable hindbrain GLP1R circuits for satiety and aversion, Nature. 632 (2024) 585–93. 10.1038/s41586-024-07685-6.

[12] W. Cheng, D. Gordian, M.Q. Ludwig, T.H. Pers, R.J. Seeley, M.G. Myers, Jr., Hindbrain circuits in the control of eating behaviour and energy balance, Nat Metab. 4 (2022) 826–35. 10.1038/s42255-022-00606-9.

[13] M.M. Veniant, S.C. Lu, L. Atangan, R. Komorowski, S. Stanislaus, Y. Cheng, et al., A GIPR antagonist conjugated to GLP-1 analogues promotes weight loss with improved metabolic parameters in preclinical and phase 1 settings, Nat Metab. 6 (2024) 290–303. 10.1038/s42255-023-00966-w.

[14] J.M. Adams, H. Pei, D.A. Sandoval, R.J. Seeley, R.B. Chang, S.D. Liberles, et al., Liraglutide Modulates Appetite and Body Weight Through Glucagon-Like Peptide 1 Receptor-Expressing Glutamatergic Neurons, Diabetes. 67 (2018) 1538–48. 10.2337/db17-1385.

[15] I. Medic, S. Gulati, H.-J. Lenz, D. Mahalingam, J. Thomas, J. Luo, et al., Initial results of a cohort of advanced prostate cancer patients in a phase Ia study of NGM120, a first-in-class anti-GDNF family receptor alpha like (GFRAL) antibody, Annals of Oncology. 33 (2022) S1186-S. 10.1016/j.annonc.2022.07.1888.

[16] L. Yang, C.C. Chang, Z. Sun, D. Madsen, H. Zhu, S.B. Padkjaer, et al., GFRAL is the receptor for GDF15 and is required for the anti-obesity effects of the ligand, Nat Med. 23 (2017) 1158–66. 10.1038/nm.4394.

[17] C.W. Roman, V.A. Derkach, R.D. Palmiter, Genetically and functionally defined NTS to PBN brain circuits mediating anorexia, Nat Commun. 7 (2016) 11905. 10.1038/ncomms11905.

[18] G.K.C. Dowsett, B.Y.H. Lam, J.A. Tadross, I. Cimino, D. Rimmington, A.P. Coll, et al., A survey of the mouse hindbrain in the fed and fasted states using single-nucleus RNA sequencing, Mol Metab. 53 (2021) 101240. 10.1016/j.molmet.2021.101240.

[19] C. Zhang, J.A. Kaye, Z. Cai, Y. Wang, S.L. Prescott, S.D. Liberles, Area Postrema Cell Types that Mediate Nausea-Associated Behaviors, Neuron. 109 (2021) 461–72 e5. 10.1016/j.neuron.2020.11.010.

[20] S.M. Fortin, R.K. Lipsky, R. Lhamo, J. Chen, E. Kim, T. Borner, et al., GABA neurons in the nucleus tractus solitarius express GLP-1 receptors and mediate anorectic effects of liraglutide in rats, Sci Transl Med. 12 (2020) 10.1126/scitranslmed.aay8071.

[21] G.X. Zheng, J.M. Terry, P. Belgrader, P. Ryvkin, Z.W. Bent, R. Wilson, et al., Massively parallel digital transcriptional profiling of single cells, Nat Commun. 8 (2017) 14049. 10.1038/ncomms14049.

[22] R.C. Team. R: A Language and Environment for Statistical Computing. Vienna, Austria: R Foundation for Statistical Computing; 2023.

[23] Y. Hao, T. Stuart, M.H. Kowalski, S. Choudhary, P. Hoffman, A. Hartman, et al., Dictionary learning for integrative, multimodal and scalable single-cell analysis, Nat Biotechnol. 42 (2024) 293–304. 10.1038/s41587-023-01767-y.

[24] C.S. McGinnis, L.M. Murrow, Z.J. Gartner, DoubletFinder: Doublet Detection in Single-Cell RNA Sequencing Data Using Artificial Nearest Neighbors, Cell Syst. 8 (2019) 329–37 e4. 10.1016/j.cels.2019.03.003.

[25] I. Korsunsky, N. Millard, J. Fan, K. Slowikowski, F. Zhang, K. Wei, et al., Fast, sensitive and accurate integration of single-cell data with Harmony, Nat Methods. 16 (2019) 1289–96. 10.1038/s41592-019-0619-0.

[26] J.N. Campbell, E.Z. Macosko, H. Fenselau, T.H. Pers, A. Lyubetskaya, D. Tenen, et al., A molecular census of arcuate hypothalamus and median eminence cell types, Nat Neurosci. 20 (2017) 484–96. 10.1038/nn.4495.

[27] R. Chen, X. Wu, L. Jiang, Y. Zhang, Single-Cell RNA-Seq Reveals Hypothalamic Cell Diversity, Cell Rep. 18 (2017) 3227–41. 10.1016/j.celrep.2017.03.004.

[28] R.A. Romanov, A. Zeisel, J. Bakker, F. Girach, A. Hellysaz, R. Tomer, et al., Molecular interrogation of hypothalamic organization reveals distinct dopamine neuronal subtypes, Nat Neurosci. 20 (2017) 176–88. 10.1038/nn.4462.

[29] B. Tasic, V. Menon, T.N. Nguyen, T.K. Kim, T. Jarsky, Z. Yao, et al., Adult mouse cortical cell taxonomy revealed by single cell transcriptomics, Nat Neurosci. 19 (2016) 335–46. 10.1038/nn.4216.

[30] D. Aran, A.P. Looney, L. Liu, E. Wu, V. Fong, A. Hsu, et al., Reference-based analysis of lung single-cell sequencing reveals a transitional profibrotic macrophage, Nat Immunol. 20 (2019) 163–72. 10.1038/s41590-018-0276-y.

[31] D. Risso, M. Cole. scRNAseq: Collection of Public Single-Cell RNA-Seq Datasets. 2.16.0 ed2023.

[32] G. Finak, A. McDavid, M. Yajima, J. Deng, V. Gersuk, A.K. Shalek, et al., MAST: a flexible statistical framework for assessing transcriptional changes and characterizing heterogeneity in single-cell RNA sequencing data, Genome Biol. 16 (2015) 278. 10.1186/s13059-015-0844-5.

[33] R. Cannoodt, W. Saelens, D. Sichien, S. Tavernier, S. Janssens, M. Guilliams, et al., SCORPIUS improves trajectory inference and identifies novel modules in dendritic cell development., Preprint at bioRxiv. (2016) 10.1101/079509.

[34] K. Ushey, J. Allaire, Y. Tang. reticulate: Interface to ’Python’. 1.35.0 ed2024.

[35] R. Cannoodt. anndata: ’anndata’ for R. 0.7.5.6 ed2023.

[36] G. Van Rossum, F.L. Drake. Python 3 Reference Manual. 3.9 ed. Scotts Valley, CA: CreateSpace; 2009.

[37] A. Sathyamurthy, K.R. Johnson, K.J.E. Matson, C.I. Dobrott, L. Li, A.R. Ryba, et al., Massively Parallel Single Nucleus Transcriptional Profiling Defines Spinal Cord Neurons and Their Activity during Behavior, Cell Rep. 22 (2018) 2216–25. 10.1016/j.celrep.2018.02.003.

[38] M.Y. Batiuk, A. Martirosyan, J. Wahis, F. de Vin, C. Marneffe, C. Kusserow, et al., Identification of region-specific astrocyte subtypes at single cell resolution, Nat Commun. 11 (2020) 1220. 10.1038/s41467-019-14198-8.

[39] L. Michielsen, M. Lotfollahi, D. Strobl, L. Sikkema, M.J.T. Reinders, F.J. Theis, et al., Single-cell reference mapping to construct and extend cell-type hierarchies, NAR Genom Bioinform. 5 (2023) lqad070. 10.1093/nargab/lqad070.

[40] A. Gayoso, R. Lopez, G. Xing, P. Boyeau, V. Valiollah Pour Amiri, J. Hong, et al., A Python library for probabilistic analysis of single-cell omics data, Nat Biotechnol. 40 (2022) 163–6. 10.1038/s41587-021-01206-w.

[41] L. Michielsen, M.J.T. Reinders, A. Mahfouz, Hierarchical progressive learning of cell identities in single-cell data, Nat Commun. 12 (2021) 2799. 10.1038/s41467-021-23196-8.

[42] J.D. Hunter, Matplotlib: A 2D graphics environment, Computing in Science & Engineering. 3 (2007) 90–5. 10.1109/MCSE.2007.55.

[43] L. Steuernagel, B.Y.H. Lam, P. Klemm, G.K.C. Dowsett, C.A. Bauder, J.A. Tadross, et al., HypoMap-a unified single-cell gene expression atlas of the murine hypothalamus, Nat Metab. 4 (2022) 1402–19. 10.1038/s42255-022-00657-y.

[44] S. Jessa, A. Blanchet-Cohen, B. Krug, M. Vladoiu, M. Coutelier, D. Faury, et al., Stalled developmental programs at the root of pediatric brain tumors, Nat Genet. 51 (2019) 1702–13. 10.1038/s41588-019-0531-7.

[45] C.E. Vaaga, M. Borisovska, G.L. Westbrook, Dual-transmitter neurons: functional implications of co-release and co-transmission, Curr Opin Neurobiol. 29 (2014) 25–32. 10.1016/j.conb.2014.04.010.

[46] N.X. Tritsch, A.J. Granger, B.L. Sabatini, Mechanisms and functions of GABA co-release, Nat Rev Neurosci. 17 (2016) 139–45. 10.1038/nrn.2015.21.

[47] C.F. Landry, J.A. Ellison, T.M. Pribyl, C. Campagnoni, K. Kampf, A.T. Campagnoni, Myelin basic protein gene expression in neurons: developmental and regional changes in protein targeting within neuronal nuclei, cell bodies, and processes., J Neurosci. 16 (1996) 2452–62. 10.1523/JNEUROSCI.16-08-02452.1996.

[48] D. Lutz, G. Loers, R. Kleene, I. Oezen, H. Kataria, N. Katagihallimath, et al., Myelin basic protein cleaves cell adhesion molecule L1 and promotes neuritogenesis and cell survival, J Biol Chem. 289 (2014) 13503–18. 10.1074/jbc.M113.530238.

[49] V.T. Cheli, D.A. Santiago Gonzalez, V. Spreuer, V. Handley, A.T. Campagnoni, P.M. Paez, Golli Myelin Basic Proteins Modulate Voltage-Operated Ca(++) Influx and Development in Cortical and Hippocampal Neurons, Mol Neurobiol. 53 (2016) 5749–71. 10.1007/s12035-015-9499-1.

[50] M. Melone, S. Ciappelloni, F. Conti, Plasma membrane transporters GAT-1 and GAT-3 contribute to heterogeneity of GABAergic synapses in neocortex, Front Neuroanat. 8 (2014) 72. 10.3389/fnana.2014.00072.

[51] S.M. Theparambil, P.S. Hosford, I. Ruminot, O. Kopach, J.R. Reynolds, P.Y. Sandoval, et al., Astrocytes regulate brain extracellular pH via a neuronal activity-dependent bicarbonate shuttle, Nat Commun. 11 (2020) 5073. 10.1038/s41467-020-18756-3.

[52] Y. Kubo, T.J. Baldwin, Y.N. Jan, L.Y. Jan, Primary structure and functional expression of a mouse inward rectifier potassium channel, Nature. 362 (1993) 127–33. 10.1038/362127a0.

[53] N. Burnashev, A. Khodorova, P. Jonas, P.J. Helm, W. Wisden, H. Monyer, et al., Calcium-Permeable AMPA-Kainate Receptors in Fusiform Cerebellar Glial Cells, Science. 256 (1992) 1566–70. 10.1126/science.1317970.

[54] J.R.P. Geiger, T. Melcher, D.S. Koh, B. Sakmann, S.P. H. P. Jonas, et al., Relative Abundance of Subunit mRNAs Determines Gating and Ca*+ Permeability of AMPA Receptors in Principal Neurons and Interneurons in Rat CNS, Neuron. 15 (1995) 193–204. doi:10.1016/0896-6273(95)90076-4.

[55] S. Marques, A. Zeisel, S. Codeluppi, D. van Bruggen, A. Mendanha Falcao, L. Xiao, et al., Oligodendrocyte heterogeneity in the mouse juvenile and adult central nervous system, Science. 352 (2016) 1326–9. 10.1126/science.aaf6463.

[56] M. Kasai, Y. Omae, Y. Kawai, A. Shibata, A. Hoshino, M. Mizuguchi, et al., GWAS identifies candidate susceptibility loci and microRNA biomarkers for acute encephalopathy with biphasic seizures and late reduced diffusion, Sci Rep. 12 (2022) 1332. 10.1038/s41598-021-04576-y.

[57] A.B. DePaula-Silva, C. Gorbea, D.J. Doty, J.E. Libbey, J.M.S. Sanchez, T.J. Hanak, et al., Differential transcriptional profiles identify microglial- and macrophage-specific gene markers expressed during virus-induced neuroinflammation, J Neuroinflammation. 16 (2019) 152. 10.1186/s12974-019-1545-x.

[58] N. Koutsodendris, J. Blumenfeld, A. Agrawal, M. Traglia, B. Grone, M. Zilberter, et al., Neuronal APOE4 removal protects against tau-mediated gliosis, neurodegeneration and myelin deficits, Nat Aging. 3 (2023) 275–96. 10.1038/s43587-023-00368-3.

[59] L. Schirmer, D. Velmeshev, S. Holmqvist, M. Kaufmann, S. Werneburg, D. Jung, et al., Neuronal vulnerability and multilineage diversity in multiple sclerosis, Nature. 573 (2019) 75–82. 10.1038/s41586-019-1404-z.

[60] J. Creus-Muncunill, J.V. Haure-Mirande, D. Mattei, J. Bons, A.V. Ramirez, B.W. Hamilton, et al., TYROBP/DAP12 knockout in Huntington’s disease Q175 mice cell-autonomously decreases microglial expression of disease-associated genes and non-cell-autonomously mitigates astrogliosis and motor deterioration, J Neuroinflammation. 21 (2024) 66. 10.1186/s12974-024-03052-4.

[61] L.L. Tang, Y.B. Wu, C.Q. Fang, P. Qu, Z.L. Gao, NDRG2 promoted secreted miR-375 in microvesicles shed from M1 microglia, which induced neuron damage, Biochem Biophys Res Commun. 469 (2016) 392–8. 10.1016/j.bbrc.2015.11.098.

[62] W. Cheng, E. Ndoka, C. Hutch, K. Roelofs, A. MacKinnon, B. Khoury, et al., Leptin receptor-expressing nucleus tractus solitarius neurons suppress food intake independently of GLP1 in mice, JCI Insight. 5 (2020) 10.1172/jci.insight.134359.

[63] C.W. Roman, S.R. Sloat, R.D. Palmiter, A tale of two circuits: CCK(NTS) neuron stimulation controls appetite and induces opposing motivational states by projections to distinct brain regions, Neuroscience. 358 (2017) 316–24. 10.1016/j.neuroscience.2017.06.049.

[64] V. Chou, R.V. Pearse, 2nd, A.J. Aylward, N. Ashour, M. Taga, G. Terzioglu, et al., INPP5D regulates inflammasome activation in human microglia, Nat Commun. 14 (2023) 7552. 10.1038/s41467-023-42819-w.

[65] S.O. Kleeman, T.M. Thakir, B. Demestichas, N. Mourikis, D. Loiero, M. Ferrer, et al., Cystatin C is glucocorticoid responsive, directs recruitment of Trem2+ macrophages, and predicts failure of cancer immunotherapy, Cell Genom. 3 (2023) 100347. 10.1016/j.xgen.2023.100347.

[66] S.H. Kang, M. Fukaya, J.K. Yang, J.D. Rothstein, D.E. Bergles, NG2+ CNS glial progenitors remain committed to the oligodendrocyte lineage in postnatal life and following neurodegeneration, Neuron. 68 (2010) 668–81. 10.1016/j.neuron.2010.09.009.

[67] T.P. Bachor, E. Hwang, E. Yulyaningsih, K. Attal, F. Mifsud, V. Pham, et al., Identification of AgRP cells in the murine hindbrain that drive feeding, Mol Metab. 80 (2024) 101886. 10.1016/j.molmet.2024.101886.

[68] M.K. Holt, The ins and outs of the caudal nucleus of the solitary tract: An overview of cellular populations and anatomical connections, J Neuroendocrinol. 34 (2022) e13132. 10.1111/jne.13132.

[69] D.M. Breen, H. Kim, D. Bennett, R.A. Calle, S. Collins, R.M. Esquejo, et al., GDF-15 Neutralization Alleviates Platinum-Based Chemotherapy-Induced Emesis, Anorexia, and Weight Loss in Mice and Nonhuman Primates, Cell Metab. 32 (2020) 938–50 e6. 10.1016/j.cmet.2020.10.023.

[70] K. Tsuzuki, S. Ishiuchi, GluR2 expressed by glial fibrillary acidic protein promoter decreases the number of neurons, Front Biosci (Landmark Ed). 13 (2008) 2784–96. 10.2741/2885.

[71] F. Endo, A. Kasai, J.S. Soto, X. Yu, Z. Qu, H. Hashimoto, et al., Molecular basis of astrocyte diversity and morphology across the CNS in health and disease, Science. 378 (2022) eadc9020. 10.1126/science.adc9020.

[72] R. Lujan, Organisation of potassium channels on the neuronal surface, J Chem Neuroanat. 40 (2010) 1–20. 10.1016/j.jchemneu.2010.03.003.

[73] R. Cabezas, M. Avila, J. Gonzalez, R.S. El-Bacha, E. Baez, L.M. Garcia-Segura, et al., Astrocytic modulation of blood brain barrier: perspectives on Parkinson’s disease, Front Cell Neurosci. 8 (2014) 211. 10.3389/fncel.2014.00211.

[74] W. Cheng, I. Gonzalez, W. Pan, A.H. Tsang, J. Adams, E. Ndoka, et al., Calcitonin Receptor Neurons in the Mouse Nucleus Tractus Solitarius Control Energy Balance via the Non-aversive Suppression of Feeding, Cell Metab. 31 (2020) 301–12 e5. 10.1016/j.cmet.2019.12.012.

[75] A.R. Patel, H. Frikke-Schmidt, P.V. Sabatini, A.C. Rupp, D.A. Sandoval, M.G. Myers, Jr.,et al., Neither GLP-1 receptors nor GFRAL neurons are required for aversive or anorectic response to DON (vomitoxin), Am J Physiol Regul Integr Comp Physiol. 324 (2023) R635–44. 10.1152/ajpregu.00189.2022.

[76] S.C. Robins, E. Trudel, O. Rotondi, X. Liu, T. Djogo, D. Kryzskaya, et al., Evidence for NG2-glia derived, adult-born functional neurons in the hypothalamus, PLoS One. 8 (2013) e78236. 10.1371/journal.pone.0078236.

[77] C.G. Liberini, T. Borner, C.N. Boyle, T.A. Lutz, The satiating hormone amylin enhances neurogenesis in the area postrema of adult rats, Mol Metab. 5 (2016) 834–43. 10.1016/j.molmet.2016.06.015.

[78] N. Wada, S. Hirako, F. Takenoya, H. Kageyama, M. Okabe, S. Shioda, Leptin and its receptors, J Chem Neuroanat. 61-62 (2014) 191-9. 10.1016/j.jchemneu.2014.09.002.

[79] H. Munzberg, J.S. Flier, C. Bjorbaek, Region-specific leptin resistance within the hypothalamus of diet-induced obese mice, Endocrinology. 145 (2004) 4880–9. 10.1210/en.2004-0726.

[80] P.M. Smith, P. Brzezinska, F. Hubert, A. Mimee, D.H. Maurice, A.V. Ferguson, Leptin influences the excitability of area postrema neurons, Am J Physiol Regul Integr Comp Physiol. 310 (2016) R440–8. 10.1152/ajpregu.00326.2015.

[81] C.G. Liberini, C.N. Boyle, C. Cifani, M. Venniro, B.T. Hope, T.A. Lutz, Amylin receptor components and the leptin receptor are co-expressed in single rat area postrema neurons, Eur J Neurosci. 43 (2016) 653–61. 10.1111/ejn.13163.

[82] L. Huo, L. Maeng, C. Bjorbaek, H.J. Grill, Leptin and the control of food intake: neurons in the nucleus of the solitary tract are activated by both gastric distension and leptin, Endocrinology. 148 (2007) 2189–97. 10.1210/en.2006-1572.

[83] J.P.H. Wilding, R.L. Batterham, S. Calanna, M. Davies, L.F. Van Gaal, I. Lingvay, et al., Once-Weekly Semaglutide in Adults with Overweight or Obesity, N Engl J Med. 384 (2021) 989–1002. 10.1056/NEJMoa2032183.

[84] B.B. Lake, R. Ai, G.E. Kaeser, N.S. Salathia, Y.C. Yung, R. Liu, et al., Neuronal subtypes and diversity revealed by single-nucleus RNA sequencing of the human brain, Science. 352 (2016) 1586–90. 10.1126/science.aaf1204.

[85] B.B. Lake, S. Codeluppi, Y.C. Yung, D. Gao, J. Chun, P.V. Kharchenko, et al., A comparative strategy for single-nucleus and single-cell transcriptomes confirms accuracy in predicted cell-type expression from nuclear RNA, Sci Rep. 7 (2017) 6031. 10.1038/s41598-017-04426-w.

[86] N. Thrupp, C. Sala Frigerio, L. Wolfs, N.G. Skene, N. Fattorelli, S. Poovathingal, et al., Single-Nucleus RNA-Seq Is Not Suitable for Detection of Microglial Activation Genes in Humans, Cell Rep. 32 (2020) 108189. 10.1016/j.celrep.2020.108189.

[87] H. Thorvaldsdottir, J.T. Robinson, J.P. Mesirov, Integrative Genomics Viewer (IGV): high-performance genomics data visualization and exploration, Brief Bioinform. 14 (2013) 178–92. 10.1093/bib/bbs017.

[88] M.J. Park, K. Chung, Endogenous lectin (RL-29) expression of the autonomic preganglionic neurons in the rat spinal cord, The Anatomical Record. 254 (1999) 54–60. 10.1002/%28SICI%291097-0185%2819990101%29254%3A1%3C53%3A%3AAID-AR7%3E3.0.CO%3B2-H.

[89] C.O. Miranda, K. Hegedus, H. Wildner, H.U. Zeilhofer, M. Antal, Morphological and neurochemical characterization of glycinergic neurons in laminae I-IV of the mouse spinal dorsal horn, J Comp Neurol. 530 (2022) 607–26. 10.1002/cne.25232.

[90] F. Guo, Y. Wang, TCF7l2, a nuclear marker that labels premyelinating oligodendrocytes and promotes oligodendroglial lineage progression, Glia. 71 (2023) 143–54. 10.1002/glia.24249.

[91] E. Weihe, C. Depboylu, B. Schutz, M.K. Schafer, L.E. Eiden, Three types of tyrosine hydroxylase-positive CNS neurons distinguished by dopa decarboxylase and VMAT2 co-expression, Cell Mol Neurobiol. 26 (2006) 659–78. 10.1007/s10571-006-9053-9.

[92] J. Sun, Y. Song, Z. Chen, J. Qiu, S. Zhu, L. Wu, et al., Heterogeneity and Molecular Markers for CNS Glial Cells Revealed by Single-Cell Transcriptomics, Cell Mol Neurobiol. 42 (2022) 2629–42. 10.1007/s10571-021-01159-3.

[93] M. Dallaporta, M.S. Bonnet, K. Horner, J. Trouslard, A. Jean, J.D. Troadec, Glial cells of the nucleus tractus solitarius as partners of the dorsal hindbrain regulation of energy balance: a proposal for a working hypothesis, Brain Res. 1350 (2010) 35–42. 10.1016/j.brainres.2010.04.025.

[94] J. Kalucka, L. de Rooij, J. Goveia, K. Rohlenova, S.J. Dumas, E. Meta, et al., Single-Cell Transcriptome Atlas of Murine Endothelial Cells, Cell. 180 (2020) 764–79 e20. 10.1016/j.cell.2020.01.015.

[95] N. Dani, R.H. Herbst, C. McCabe, G.S. Green, K. Kaiser, J.P. Head, et al., A cellular and spatial map of the choroid plexus across brain ventricles and ages, Cell. 184 (2021) 3056–74 e21. 10.1016/j.cell.2021.04.003.

[96] B. Ray, J.A. Bailey, S. Sarkar, D.K. Lahiri, Molecular and immunocytochemical characterization of primary neuronal cultures from adult rat brain: Differential expression of neuronal and glial protein markers, J Neurosci Methods. 184 (2009) 294–302. 10.1016/j.jneumeth.2009.08.018.

[97] S.E. Lauri, M. Ryazantseva, E. Orav, A. Vesikansa, T. Taira, Kainate receptors in the developing neuronal networks, Neuropharmacology. 195 (2021) 108585. 10.1016/j.neuropharm.2021.108585.

[98] Y.H. Song, J. Yoon, S.H. Lee, The role of neuropeptide somatostatin in the brain and its application in treating neurological disorders, Exp Mol Med. 53 (2021) 328–38. 10.1038/s12276-021-00580-4.

[99] H.W. Song, K.L. Foreman, B.D. Gastfriend, J.S. Kuo, S.P. Palecek, E.V. Shusta, Transcriptomic comparison of human and mouse brain microvessels, Sci Rep. 10 (2020) 12358. 10.1038/s41598-020-69096-7.

[100] L. Xu, Y. Yao, Central Nervous System Fibroblast-Like Cells in Stroke and Other Neurological Disorders, Stroke. 52 (2021) 2456–64. 10.1161/STROKEAHA.120.033431.

[101] U. Ernsberger, H. Rohrer, Sympathetic tales: subdivisons of the autonomic nervous system and the impact of developmental studies, Neural Dev. 13 (2018) 20. 10.1186/s13064-018-0117-6.

[102] C.R. Gajera, H. Emich, O. Lioubinski, A. Christ, R. Beckervordersandforth-Bonk, K. Yoshikawa, et al., LRP2 in ependymal cells regulates BMP signaling in the adult neurogenic niche, J Cell Sci. 123 (2010) 1922–30. 10.1242/jcs.065912.

[103] A. MacDonald, B. Lu, M. Caron, N. Caporicci-Dinucci, D. Hatrock, K. Petrecca, et al., Single Cell Transcriptomics of Ependymal Cells Across Age, Region and Species Reveals Cilia-Related and Metal Ion Regulatory Roles as Major Conserved Ependymal Cell Functions, Front Cell Neurosci. 15 (2021) 703951. 10.3389/fncel.2021.703951.

[104] H.W. Jeong, R. Dieguez-Hurtado, H. Arf, J. Song, H. Park, K. Kruse, et al., Single-cell transcriptomics reveals functionally specialized vascular endothelium in brain, Elife. 11 (2022) 10.7554/eLife.57520.

[105] K. Huizer, D.A.M. Mustafa, J.C. Spelt, J.M. Kros, A. Sacchetti, Improving the characterization of endothelial progenitor cell subsets by an optimized FACS protocol, PLoS One. 12 (2017) e0184895. 10.1371/journal.pone.0184895.

[106] E. Gerrits, Y. Heng, E. Boddeke, B.J.L. Eggen, Transcriptional profiling of microglia; current state of the art and future perspectives, Glia. 68 (2020) 740–55. 10.1002/glia.23767.

[107] P.T. Shah, J.A. Stratton, M.G. Stykel, S. Abbasi, S. Sharma, K.A. Mayr, et al., Single-Cell Transcriptomics and Fate Mapping of Ependymal Cells Reveals an Absence of Neural Stem Cell Function, Cell. 173 (2018) 1045–57 e9. 10.1016/j.cell.2018.03.063.

[108] C.G. Causing, A. Gloster, R. Aloyz, S.X. Bamji, E. Chang, J. Fawcett, et al., Synaptic Innervation Density Is Regulated by Neuron-Derived BDNF, Neuron. 18 (1997) 257–67. https://www.cell.com/fulltext/S0896-6273(00)80266-4#secd33944447e474.

[109] Y. Zhang, S.A. Sloan, L.E. Clarke, C. Caneda, C.A. Plaza, P.D. Blumenthal, et al., Purification and Characterization of Progenitor and Mature Human Astrocytes Reveals Transcriptional and Functional Differences with Mouse, Neuron. 89 (2016) 37–53. 10.1016/j.neuron.2015.11.013.

[110] F. Langlet, Tanycyte Gene Expression Dynamics in the Regulation of Energy Homeostasis, Front Endocrinol (Lausanne). 10 (2019) 286. 10.3389/fendo.2019.00286.

[111] O. Uriarte Huarte, D. Kyriakis, T. Heurtaux, Y. Pires-Afonso, K. Grzyb, R. Halder, et al., Single-Cell Transcriptomics and In Situ Morphological Analyses Reveal Microglia Heterogeneity Across the Nigrostriatal Pathway, Front Immunol. 12 (2021) 639613. 10.3389/fimmu.2021.639613.

[112] R.E. Pepper, K.A. Pitman, C.L. Cullen, K.M. Young, How Do Cells of the Oligodendrocyte Lineage Affect Neuronal Circuits to Influence Motor Function, Memory and Mood?, Front Cell Neurosci. 12 (2018) 399. 10.3389/fncel.2018.00399.

[113] A. Maegawa, K. Murata, K. Kuroda, S. Fujieda, Y. Fukazawa, Cellular Profiles of Prodynorphin and Preproenkephalin mRNA-Expressing Neurons in the Anterior Olfactory Tubercle of Mice, Front Neural Circuits. 16 (2022) 908964. 10.3389/fncir.2022.908964.

[114] S.T. Hentges, V. Otero-Corchon, R.L. Pennock, C.M. King, M.J. Low, Proopiomelanocortin expression in both GABA and glutamate neurons, J Neurosci. 29 (2009) 13684–90. 10.1523/JNEUROSCI.3770-09.2009.

[115] W. Sun, A. Cornwell, J. Li, S. Peng, M.J. Osorio, N. Aalling, et al., SOX9 Is an Astrocyte-Specific Nuclear Marker in the Adult Brain Outside the Neurogenic Regions, J Neurosci. 37 (2017) 4493–507. 10.1523/JNEUROSCI.3199-16.2017.

[116] A.T. Lee, D. Vogt, J.L. Rubenstein, V.S. Sohal, A class of GABAergic neurons in the prefrontal cortex sends long-range projections to the nucleus accumbens and elicits acute avoidance behavior, J Neurosci. 34 (2014) 11519–25. 10.1523/JNEUROSCI.1157-14.2014.

[117] M. Vanlandewijck, L. He, M.A. Mae, J. Andrae, K. Ando, F. Del Gaudio, et al., A molecular atlas of cell types and zonation in the brain vasculature, Nature. 554 (2018) 475–80. 10.1038/nature25739.

[118] M. Carlen, K. Meletis, C. Goritz, V. Darsalia, E. Evergren, K. Tanigaki, et al., Forebrain ependymal cells are Notch-dependent and generate neuroblasts and astrocytes after stroke, Nat Neurosci. 12 (2009) 259–67. 10.1038/nn.2268.

[119] J. Garcia-Revilla, A. Boza-Serrano, A.M. Espinosa-Oliva, M.S. Soto, T. Deierborg, R. Ruiz, et al., Galectin-3, a rising star in modulating microglia activation under conditions of neurodegeneration, Cell Death Dis. 13 (2022) 628. 10.1038/s41419-022-05058-3.

[120] A.C. Yang, R.T. Vest, F. Kern, D.P. Lee, M. Agam, C.A. Maat, et al., A human brain vascular atlas reveals diverse mediators of Alzheimer’s risk, Nature. 603 (2022) 885–92. 10.1038/s41586-021-04369-3.

[121] S. Chasseigneaux, Y. Moraca, V. Cochois-Guegan, A.C. Boulay, A. Gilbert, S. Le Crom, et al., Isolation and differential transcriptome of vascular smooth muscle cells and mid-capillary pericytes from the rat brain, Sci Rep. 8 (2018) 12272. 10.1038/s41598-018-30739-5.

[122] Z. Gao, K. Ure, J.L. Ables, D.C. Lagace, K.A. Nave, S. Goebbels, et al., Neurod1 is essential for the survival and maturation of adult-born neurons, Nat Neurosci. 12 (2009) 1090–2. 10.1038/nn.2385.

[123] T. Kodama, S. Guerrero, M. Shin, S. Moghadam, M. Faulstich, S. du Lac, Neuronal classification and marker gene identification via single-cell expression profiling of brainstem vestibular neurons subserving cerebellar learning, J Neurosci. 32 (2012) 7819–31. 10.1523/JNEUROSCI.0543-12.2012.

[124] J. Manjarrez-Marmolejo, J. Franco-Pérez, Gap Junction Blockers-An Overview of their Effects on Induced Seizures in Animal Models, Current Neuropharmacology. 14 (2016) 759–71. 10.2174/1570159X14666160603115.

[125] T. Paretkar, E. Dimitrov, Activation of enkephalinergic (Enk) interneurons in the central amygdala (CeA) buffers the behavioral effects of persistent pain, Neurobiol Dis. 124 (2019) 364–72. 10.1016/j.nbd.2018.12.005.

[126] Z. Zhang, Z. Ma, W. Zou, H. Guo, M. Liu, Y. Ma, et al., The Appropriate Marker for Astrocytes: Comparing the Distribution and Expression of Three Astrocytic Markers in Different Mouse Cerebral Regions, Biomed Res Int. 2019 (2019) 9605265. 10.1155/2019/9605265.

[127] A.M. Jurga, M. Paleczna, J. Kadluczka, K.Z. Kuter, Beyond the GFAP-Astrocyte Protein Markers in the Brain, Biomolecules. 11 (2021) 10.3390/biom11091361.

[128] V. Dyachuk, A. Furlan, M.K. Shahidi, M. Giovenco, N. Kaukua, C. Konstantinidou, et al., Parasympathetic neurons originate from nerve-associated peripheral glial progenitors, Science. 345 (2014) 82–7. 10.1126/science.1253281.

[129] D.S. Liu, T.L. Xu, Cell-Type Identification in the Autonomic Nervous System, Neurosci Bull. 35 (2019) 145–55. 10.1007/s12264-018-0284-9.

[130] B.J. Kang, D.A. Chang, D.D. Mackay, G.H. West, T.S. Moreira, A.C. Takakura, et al., Central nervous system distribution of the transcription factor Phox2b in the adult rat, J Comp Neurol. 503 (2007) 627–41. 10.1002/cne.21409.

[131] M. Adler, N. Moriel, A. Goeva, I. Avraham-Davidi, S. Mages, T.S. Adams, et al., Emergence of division of labor in tissues through cell interactions and spatial cues, Cell Rep. 42 (2023) 112412. 10.1016/j.celrep.2023.112412.

[132] A. Steimel, J. Suh, A. Hussainkhel, S. Deheshi, J.M. Grants, R. Zapf, et al., The C. elegans CDK8 Mediator module regulates axon guidance decisions in the ventral nerve cord and during dorsal axon navigation, Dev Biol. 377 (2013) 385–98. 10.1016/j.ydbio.2013.02.009.

[133] T. Daiber, C.J. VanderZwan-Butler, G.J. Bashaw, T.A. Evans, Conserved and divergent aspects of Robo receptor signaling and regulation between Drosophila Robo1 and C. elegans SAX-3, Genetics. 217 (2021) 10.1093/genetics/iyab018.

[134] S. Schmetsdorf, U. Gärtner, T. Arendt, Constitutive Expression of Functionally Active Cyclin-Dependent Kinases and Their Binding Partners Suggests Noncanonical Functions of Cell Cycle Regulators in Differentiated Neurons, Cerebral Cortex. 17 (2007) 1821–9. 10.1093/cercor/bhl091.

[135] K.K. Kummer, M. Zeidler, T. Kalpachidou, M. Kress, Role of IL-6 in the regulation of neuronal development, survival and function, Cytokine. 144 (2021) 155582. 10.1016/j.cyto.2021.155582.

[136] P. März, J.G. Cheng, R.A. Gadient, P.H. Patterson, T. Stoyan, U. Otten, et al., Sympathetic neurons can produce and respond to interleukin 6, Proc Natl Acad Sci. 95 (1998) 3251–6. 10.1073/pnas.95.6.3251.

[137] K.M. McGrail, J.M. Phillips, K.J. Sweadner, Immunofluorescent localization of three Na,K-ATPase isozymes in the rat central nervous system: both neurons and glia can express more than one Na,K-ATPase., J Neurosci. 11 (1991) 381–91. 10.1523/JNEUROSCI.11-02-00381.1991.

[138] H. Poulsen, H. Khandelia, J.P. Morth, M. Bublitz, O.G. Mouritsen, J. Egebjerg, et al., Neurological disease mutations compromise a C-terminal ion pathway in the Na(+)/K(+)-ATPase, Nature. 467 (2010) 99–102. 10.1038/nature09309.

[139] A. Abousaab, J. Warsi, B. Elvira, F. Lang, Caveolin-1 Sensitivity of Excitatory Amino Acid Transporters EAAT1, EAAT2, EAAT3, and EAAT4, J Membr Biol. 249 (2016) 239–49. 10.1007/s00232-015-9863-0.

[140] E.R. Bongarzone, C.W. Campagnoni, K. Kampf, E.C. Jacobs, V.W. Handley, V. Schonmann, et al., Identification of a new exon in the myelin proteolipid protein gene encoding novel protein isoforms that are restricted to the somata of oligodendrocytes and neurons., J Neurosci. 19 (1999) 8349–57. 10.1523/JNEUROSCI.19-19-08349.

[141] L.F. Nunnelly, M. Campbell, D.I. Lee, P. Dummer, G. Gu, V. Menon, et al., St18 specifies globus pallidus projection neuron identity in MGE lineage, Nat Commun. 13 (2022) 7735. 10.1038/s41467-022-35518-5.

[142] C. Plachez, C. Lindwall, N. Sunn, M. Piper, R.X. Moldrich, C.E. Campbell, et al., Nuclear factor I gene expression in the developing forebrain, J Comp Neurol. 508 (2008) 385–401. 10.1002/cne.21645.

[143] J. Fraser, A. Essebier, A.S. Brown, R.A. Davila, D. Harkins, O. Zalucki, et al., Common Regulatory Targets of NFIA, NFIX and NFIB during Postnatal Cerebellar Development, Cerebellum. 19 (2020) 89–101. 10.1007/s12311-019-01089-3.

[144] L. Dubois, A. Vincent, The COE--Collier/Olf1/EBF--transcription factors: structural conservation and diversity of developmental functions, 108. 3–12 (2001) 10.1016/s0925-4773(01)00486-5.

[145] Y. Liu, T. Lear, O. Iannone, S. Shiva, C. Corey, S. Rajbhandari, et al., The Proapoptotic F-box Protein Fbxl7 Regulates Mitochondrial Function by Mediating the Ubiquitylation and Proteasomal Degradation of Survivin, J Biol Chem. 290 (2015) 11843–52. 10.1074/jbc.M114.629931.

[146] H. Kim, N.C. Berens, N.E. Ochandarena, B.D. Philpot, Region and Cell Type Distribution of TCF4 in the Postnatal Mouse Brain, Front Neuroanat. 14 (2020) 42. 10.3389/fnana.2020.00042.

[147] A. Sirp, A. Shubina, J. Tuvikene, L. Tamberg, C.S. Kiir, L. Kranich, et al., Expression of alternative transcription factor 4 mRNAs and protein isoforms in the developing and adult rodent and human tissues, Front Mol Neurosci. 15 (2022) 1033224. 10.3389/fnmol.2022.1033224.

[148] R.D. Hodge, T.E. Bakken, J.A. Miller, K.A. Smith, E.R. Barkan, L.T. Graybuck, et al., Conserved cell types with divergent features in human versus mouse cortex, Nature. 573 (2019) 61–8. 10.1038/s41586-019-1506-7.

[149] T. Kuroda, S. Yasuda, S. Tachi, S. Matsuyama, S. Kusakawa, K. Tano, et al., SALL3 expression balance underlies lineage biases in human induced pluripotent stem cell differentiation, Nat Commun. 10 (2019) 2175. 10.1038/s41467-019-09511-4.

[150] F. Morello, D. Borshagovski, M. Survila, L. Tikker, S. Sadik-Ogli, A. Kirjavainen, et al., Molecular Fingerprint and Developmental Regulation of the Tegmental GABAergic and Glutamatergic Neurons Derived from the Anterior Hindbrain, Cell Rep. 33 (2020) 108268. 10.1016/j.celrep.2020.108268.

[151] L. Wolf, C.S. Gao, K. Gueta, Q. Xie, T. Chevallier, N.R. Podduturi, et al., Identification and characterization of FGF2-dependent mRNA: microRNA networks during lens fiber cell differentiation, G3 (Bethesda). 3 (2013) 2239–55. 10.1534/g3.113.008698.

[152] H. Liu, M. Aramaki, Y. Fu, D. Forrest, Retinoid-Related Orphan Receptor beta and Transcriptional Control of Neuronal Differentiation, Curr Top Dev Biol. 125 (2017) 227–55. 10.1016/bs.ctdb.2016.11.009.

[153] M.G. Del Barrio, S. Bourane, K. Grossmann, R. Schule, S. Britsch, D.D. O’Leary, et al., A transcription factor code defines nine sensory interneuron subtypes in the mechanosensory area of the spinal cord, PLoS One. 8 (2013) e77928. 10.1371/journal.pone.0077928.

[154] Y. Li, A. Osuma, E. Correa, M.A. Okebalama, P. Dao, O. Gaylord, et al., Establishment and maintenance of motor neuron identity via temporal modularity in terminal selector function, Elife. 9 (2020) 10.7554/eLife.59464.

[155] D. Moruzzo, L. Nobbio, B. Sterlini, G.G. Consalez, F. Benfenati, A. Schenone, et al., The Transcription Factors EBF1 and EBF2 Are Positive Regulators of Myelination in Schwann Cells, Mol Neurobiol. 54 (2017) 8117–27. 10.1007/s12035-016-0296-2.

[156] M.A. Jimenez, P. Akerblad, M. Sigvardsson, E.D. Rosen, Critical role for Ebf1 and Ebf2 in the adipogenic transcriptional cascade, Mol Cell Biol. 27 (2007) 743–57. 10.1128/MCB.01557-06.

[157] H. Yu, M. Rubinstein, M.J. Low, Developmental single-cell transcriptomics of hypothalamic POMC neurons reveal the genetic trajectories of multiple neuropeptidergic phenotypes, Elife. 11 (2022) 10.7554/eLife.72883.

[158] I. Singh, L. Wang, B. Xia, J. Liu, A. Tahiri, A. El Ouaamari, et al., Activation of arcuate nucleus glucagon-like peptide-1 receptor-expressing neurons suppresses food intake, Cell Biosci. 12 (2022) 178. 10.1186/s13578-022-00914-3.

[159] L. Xiao, D.P. Merullo, T.M.I. Koch, M. Cao, M. Co, A. Kulkarni, et al., Expression of FoxP2 in the basal ganglia regulates vocal motor sequences in the adult songbird, Nat Commun. 12 (2021) 2617. 10.1038/s41467-021-22918-2.

[160] T.T. Du, J.B. Dewey, E.L. Wagner, R. Cui, J. Heo, J.J. Park, et al., LMO7 deficiency reveals the significance of the cuticular plate for hearing function, Nat Commun. 10 (2019) 1117. 10.1038/s41467-019-09074-4.

[161] M.A. Wozniak, B.M. Baker, C.S. Chen, K.L. Wilson, The emerin-binding transcription factor Lmo7 is regulated by association with p130Cas at focal adhesions, PeerJ. 1 (2013) e134. 10.7717/peerj.134.

[162] T. Kawaue, A. Shitamukai, A. Nagasaka, Y. Tsunekawa, T. Shinoda, K. Saito, et al., Lzts1 controls both neuronal delamination and outer radial glial-like cell generation during mammalian cerebral development, Nat Commun. 10 (2019) 2780. 10.1038/s41467-019-10730-y.

[163] O. Randlett, L. Poggi, F.R. Zolessi, W.A. Harris, The oriented emergence of axons from retinal ganglion cells is directed by laminin contact in vivo, Neuron. 70 (2011) 266–80. 10.1016/j.neuron.2011.03.013.

[164] N. Ichikawa-Tomikawa, J. Ogawa, V. Douet, Z. Xu, Y. Kamikubo, T. Sakurai, et al., Laminin alpha1 is essential for mouse cerebellar development, Matrix Biol. 31 (2012) 17–28. 10.1016/j.matbio.2011.09.002.

[165] E.S. Liau, S. Jin, Y.C. Chen, W.S. Liu, M. Calon, S. Nedelec, et al., Single-cell transcriptomic analysis reveals diversity within mammalian spinal motor neurons, Nat Commun. 14 (2023) 46. 10.1038/s41467-022-35574-x.

