## Supplemental Figures for "A unified rodent atlas reveals the cellular complexity and evolutionary divergence of the dorsal vagal complex"

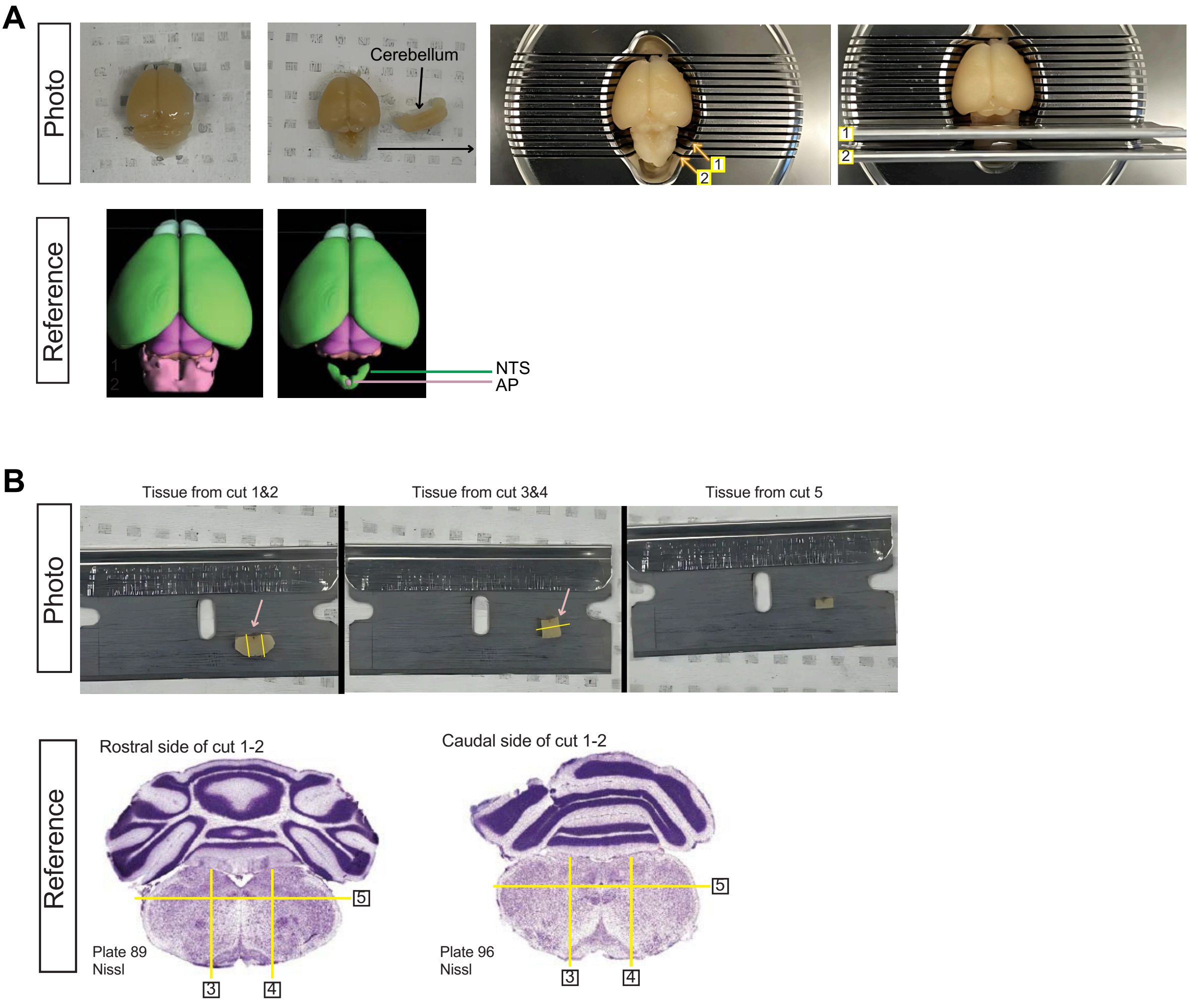
A
Photo
Cerebellum
1
2
Reference
NTS
AP
B
Tissue from cut 1&2
Tissue from cut 3&4
Tissue from cut 5
Photo
Reference
Rostral side of cut 1-2
Caudal side of cut 1-2
Plate 89
Nissl
Plate 96
Nissl
3
4
5

Figure S1

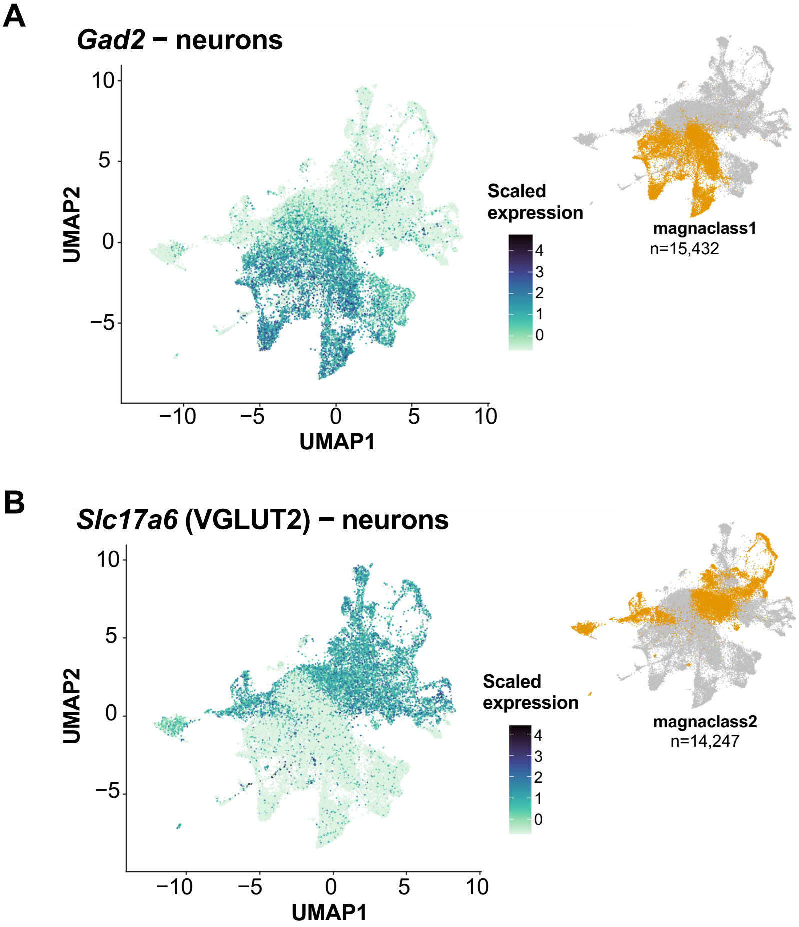
A
Gad2 − neurons
UMAP2
UMAP1
Scaled expression
magnaclass1
n=15,432
B
Slc17a6 (VGLUT2) − neurons
UMAP2
UMAP1
Scaled expression
magnaclass2
n=14,247

Figure S2

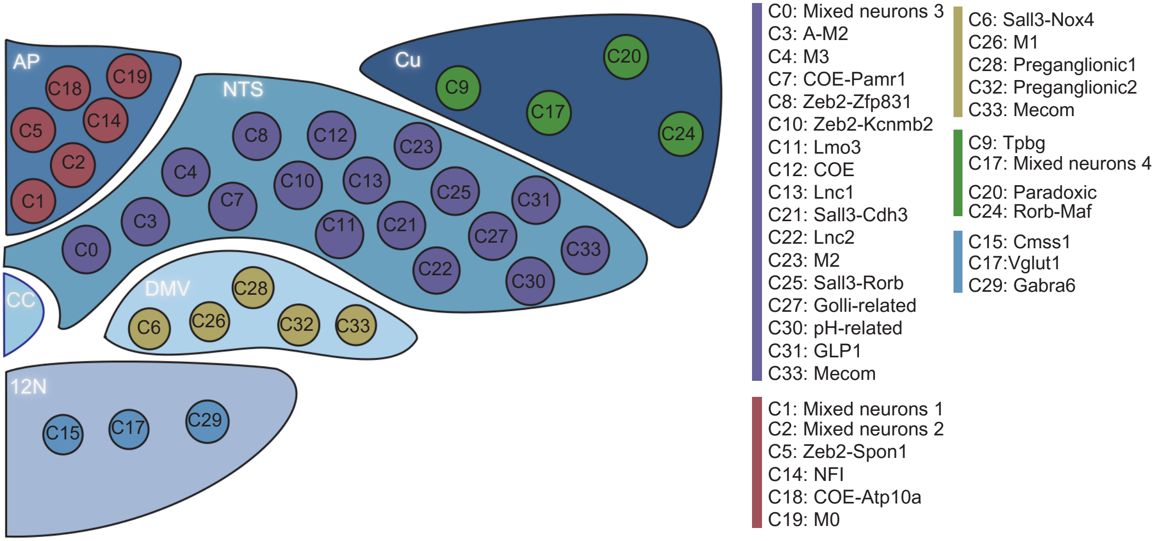
AP
C18
C19
C5
C14
C2
C1
NTS
C8
C12
C23
C4
C10
C13
C25
C31
C7
C3
C11
C21
C27
C33
C0
C22
C30
Cu
C9
C20
C17
C24
CC
DMV
C28
C6
C26
C32
C33
12N
C15
C17
C29
C0: Mixed neurons 3
C3: A-M2
C4: M3
C7: COE-Pamr1
C8: Zeb2-Zfp831
C10: Zeb2-Kcnmb2
C11: Lmo3
C12: COE
C13: Lnc1
C21: Sall3-Cdh3
C22: Lnc2
C23: M2
C25: Sall3-Rorb
C27: Golli-related
C30: pH-related
C31: GLP1
C33: Mecom
C1: Mixed neurons 1
C2: Mixed neurons 2
C5: Zeb2-Spon1
C14: NFI
C18: COE-Atp10a
C19: M0
C6: Sall3-Nox4
C26: M1
C28: Preganglionic1
C32: Preganglionic2
C33: Mecom
C9: Tpbg
C17: Mixed neurons 4
C20: Paradoxic
C24: Rorb-Maf
C15: Cmss1
C17:Vglut1
C29: Gabra6

**Figure S3**

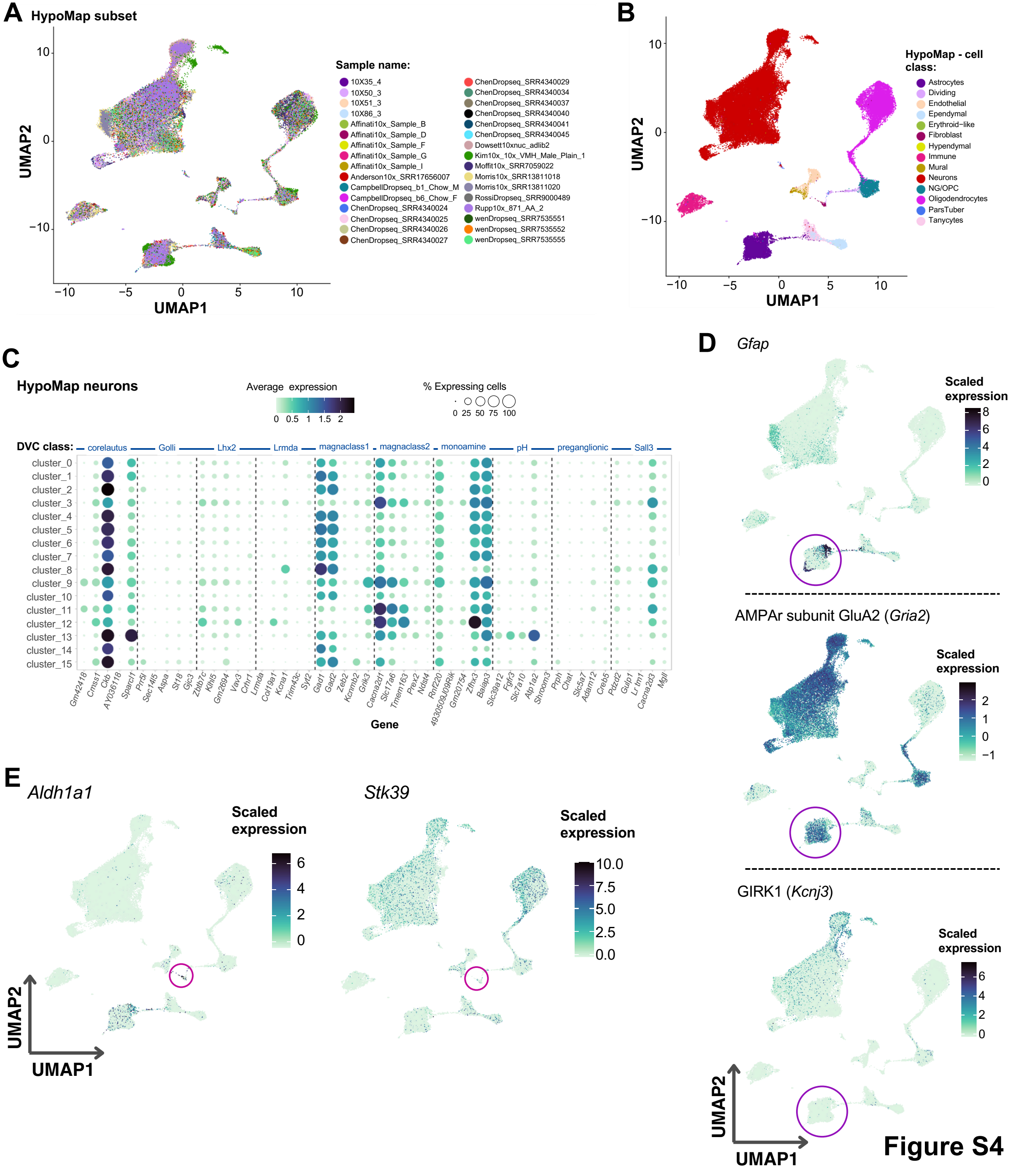
A
HypoMap subset
UMAP1
UMAP2
Sample name:
10X35_4
10X50_3
10X51_3
10X86_3
Affinati10x_Sample_B
Affinati10x_Sample_D
Affinati10x_Sample_F
Affinati10x_Sample_G
Affinati10x_Sample_I
Anderson10x_SRR17656007
CampbellDropseq_b1_Chow_M
CampbellDropseq_b6_Chow_F
ChenDropseq_SRR4340024
ChenDropseq_SRR4340025
ChenDropseq_SRR4340026
ChenDropseq_SRR4340027
ChenDropseq_SRR4340029
ChenDropseq_SRR4340034
ChenDropseq_SRR4340037
ChenDropseq_SRR4340040
ChenDropseq_SRR4340041
ChenDropseq_SRR4340045
Dowsett10xnuc_adlib2
Kim10x_10x_VMH_Male_Plain_1
Moffit10x_SRR7059022
Morris10x_SRR13811018
Morris10x_SRR13811020
RossiDropseq_SRR9000489
Rupp10x_871_AA_2
wenDropseq_SRR7535551
wenDropseq_SRR7535552
wenDropseq_SRR7535555
B
HypoMap - cell class:
Astrocytes
Dividing
Endothelial
Ependymal
Erythroid−like
Fibroblast
Hypendymal
Immune
Mural
Neurons
NG/OPC
Oligodendrocytes
ParsTuber
Tanycytes
C
HypoMap neurons
Average expression
% Expressing cells
DVC class:
corelautus
Golli
Lhx2
Lrmda
magnaclass1
magnaclass2
monoamine
pH
preganglionic
Sall3
cluster_0
cluster_1
cluster_2
cluster_3
cluster_4
cluster_5
cluster_6
cluster_7
cluster_8
cluster_9
cluster_10
cluster_11
cluster_12
cluster_13
cluster_14
cluster_15
Gm42418 Cmss1 Ckb AY036118 Sparcl1 Prr5l Sec14l5 Aspa St18 Gjc3 Zbtb7c Klhl5 Gm2694 Vav3 Crhr1 Lrmda Col19a1 Kcna1 Trim43c Syt2 Gad1 Gad2 Zeb2 Kcnmb2 Grik3 Cacna2d1 Slc17a6 Tmem163 Prex2 Ndst4 Rnf220 4930509J09Rik Gm20754 Zfhx3 Baiap3 Slc39a12 Fgfr3 Slc7a10 Atp1a2 Shroom3 Prph Chat Slc5a7 Adam12 Creb5 Pdzd2 Gulp1 Lrtm1 Cacna2d3 Mgll
Gene
D
Gfap
Scaled expression
AMPAr subunit GluA2 (Gria2)
GIRK1 (Kcnj3)
E
Aldh1a1
Stk39
UMAP1
UMAP2

Figure S4

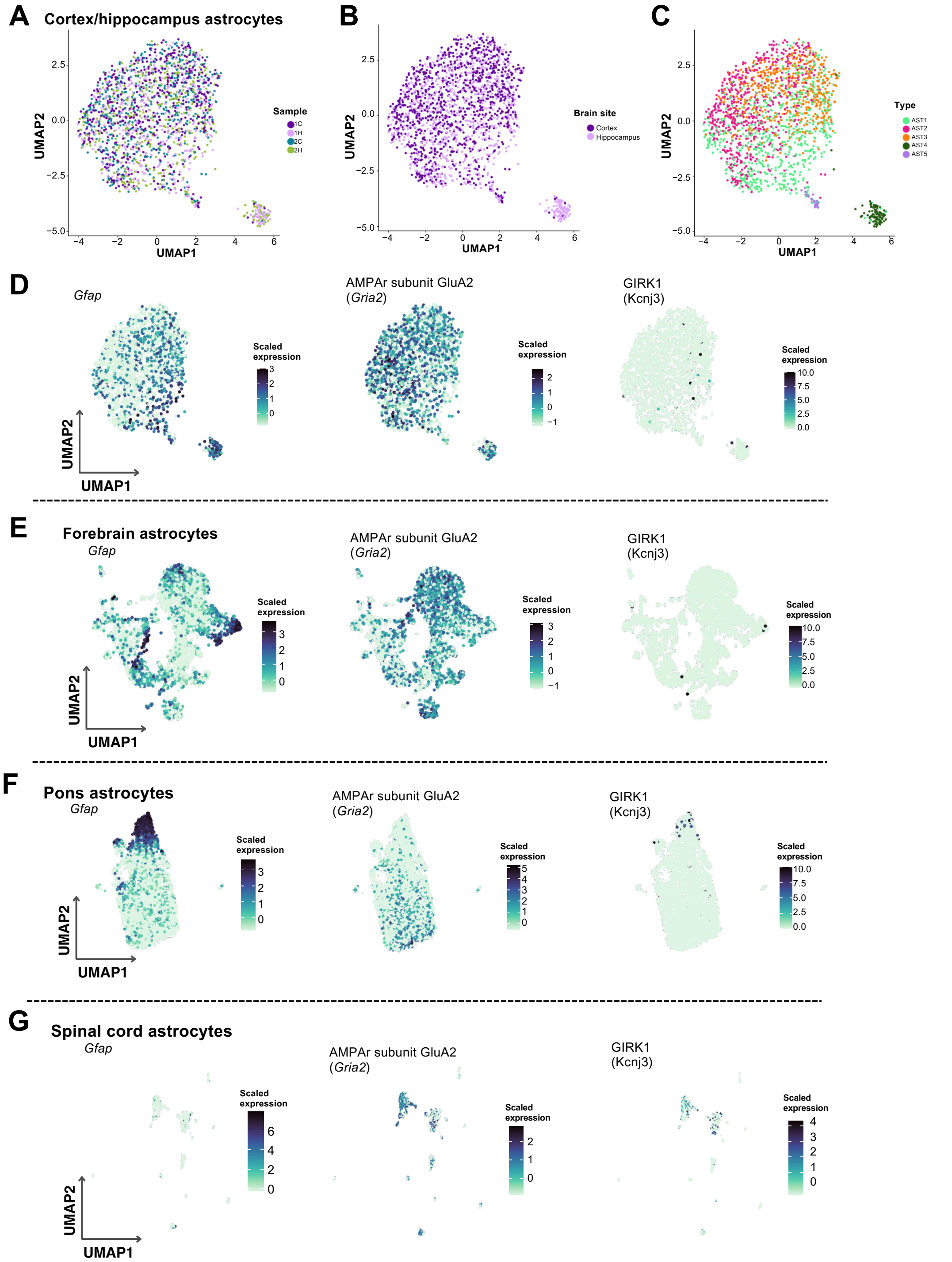
A
Cortex/hippocampus astrocytes
UMAP2
UMAP1
Sample
1C
1H
2C
2H
B
Brain site
Cortex
Hippocampus
C
Type
AST1
AST2
AST3
AST4
AST5
D
Gfap
AMPAr subunit GluA2
(Gria2)
GIRK1
(Kcnj3)
Scaled expression
E
Forebrain astrocytes
F
Pons astrocytes
G
Spinal cord astrocytes

**Figure S5**

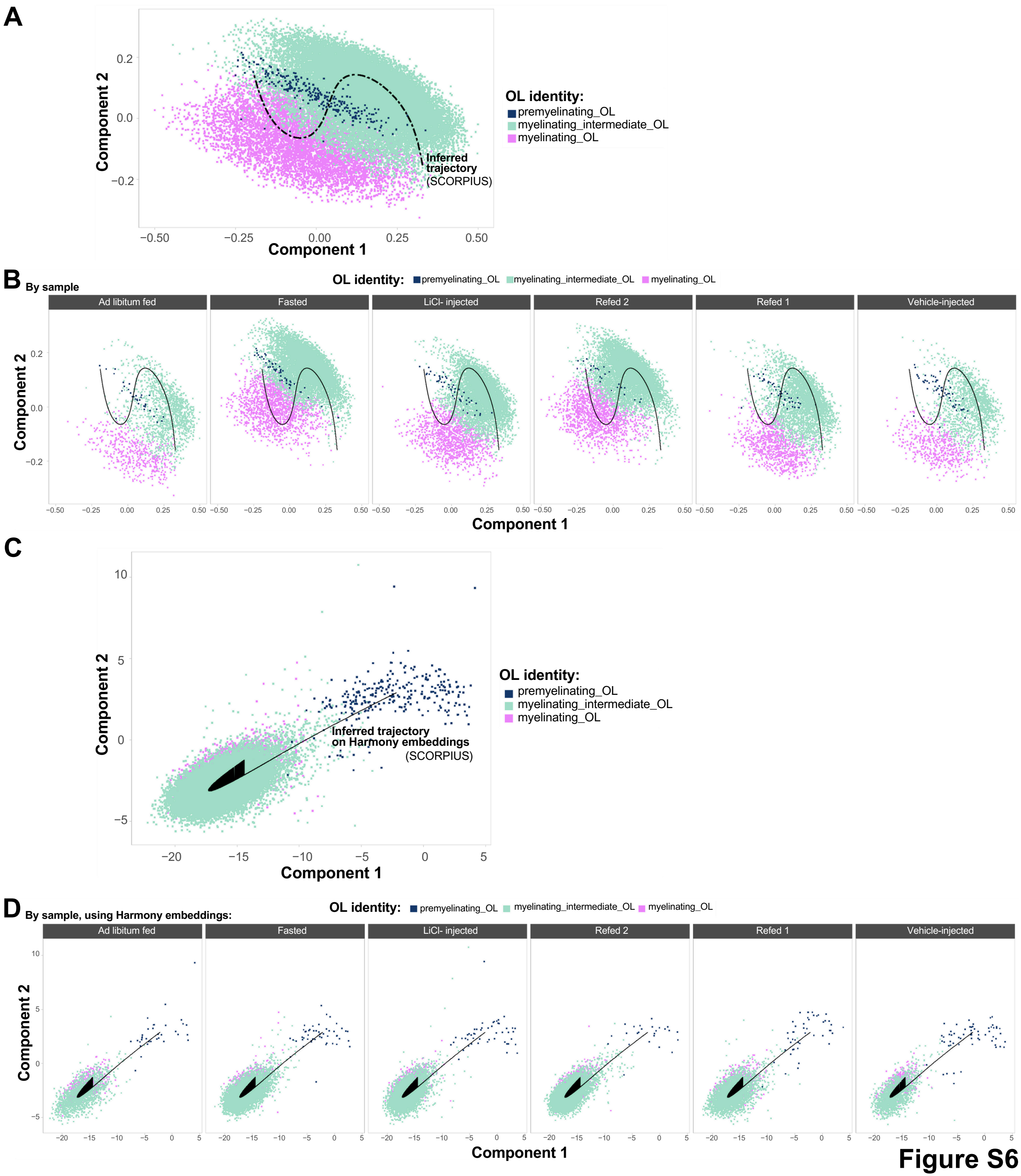
A
Component 2
Component 1
OL identity:
premyelinating_OL
myelinating_intermediate_OL
myelinating_OL
Inferred trajectory (SCORPIUS)
B By sample
OL identity: premyelinating_OL myelinating_intermediate_OL myelinating_OL
Ad libitum fed
Fasted
LiCl- injected
Refed 2
Refed 1
Vehicle-injected
C
Inferred trajectory on Harmony embeddings (SCORPIUS)
D By sample, using Harmony embeddings:
Figure S6

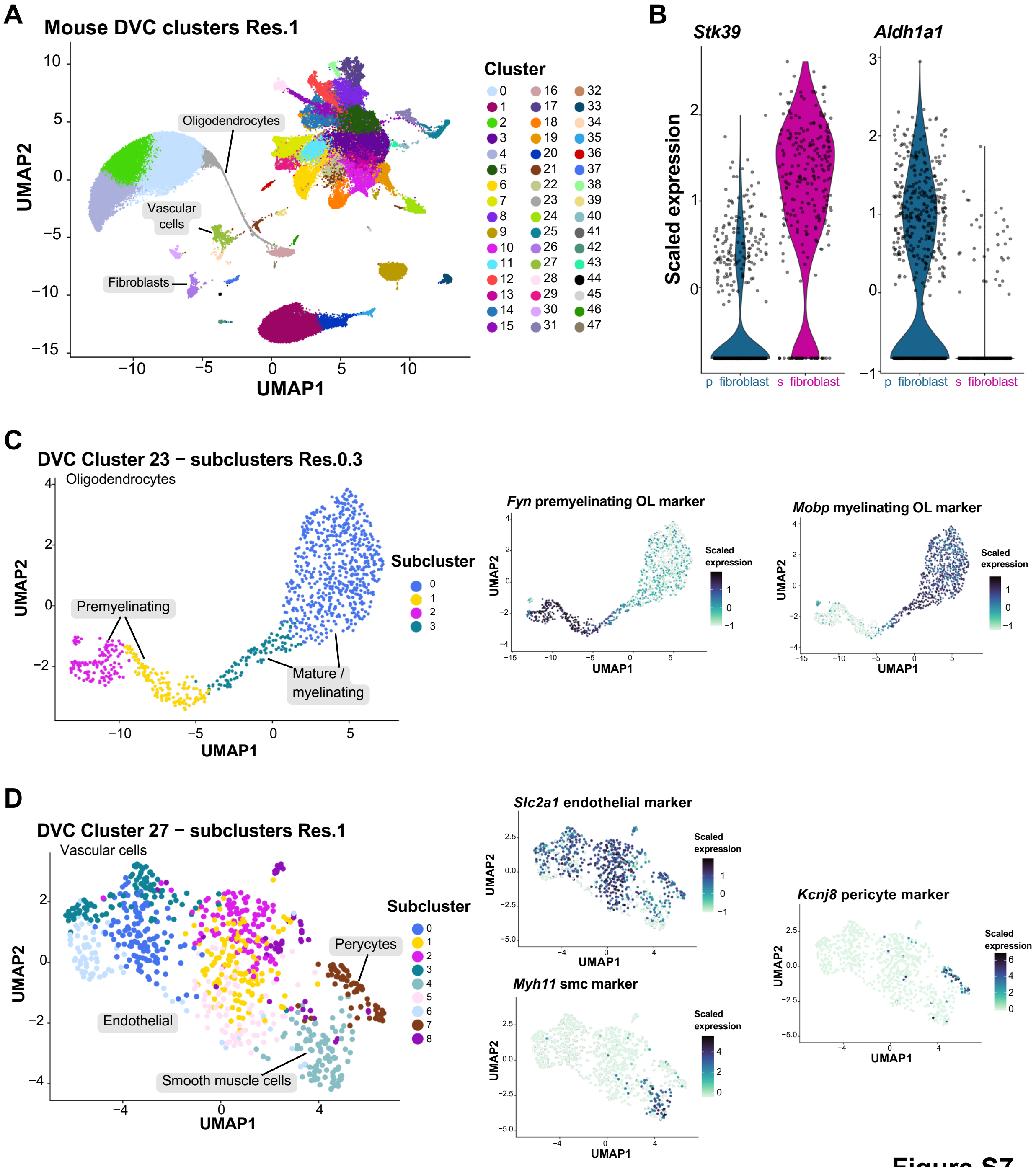

A
Mouse DVC clusters Res.1
Oligodendrocytes
Vascular cells
Fibroblasts
UMAP1
UMAP2
Cluster
B
Stk39
Aldh1a1
Scaled expression
p_fibroblast
s_fibroblast
C
DVC Cluster 23 − subclusters Res.0.3
Oligodendrocytes
Premyelinating
Mature / myelinating
Subcluster
Fyn premyelinating OL marker
Mobp myelinating OL marker
Scaled expression
D
DVC Cluster 27 − subclusters Res.1
Vascular cells
Endothelial
Perycytes
Smooth muscle cells
Subcluster
Slc2a1 endothelial marker
Myh11 smc marker
Kcnj8 pericyte marker
Scaled expression


Figure S7

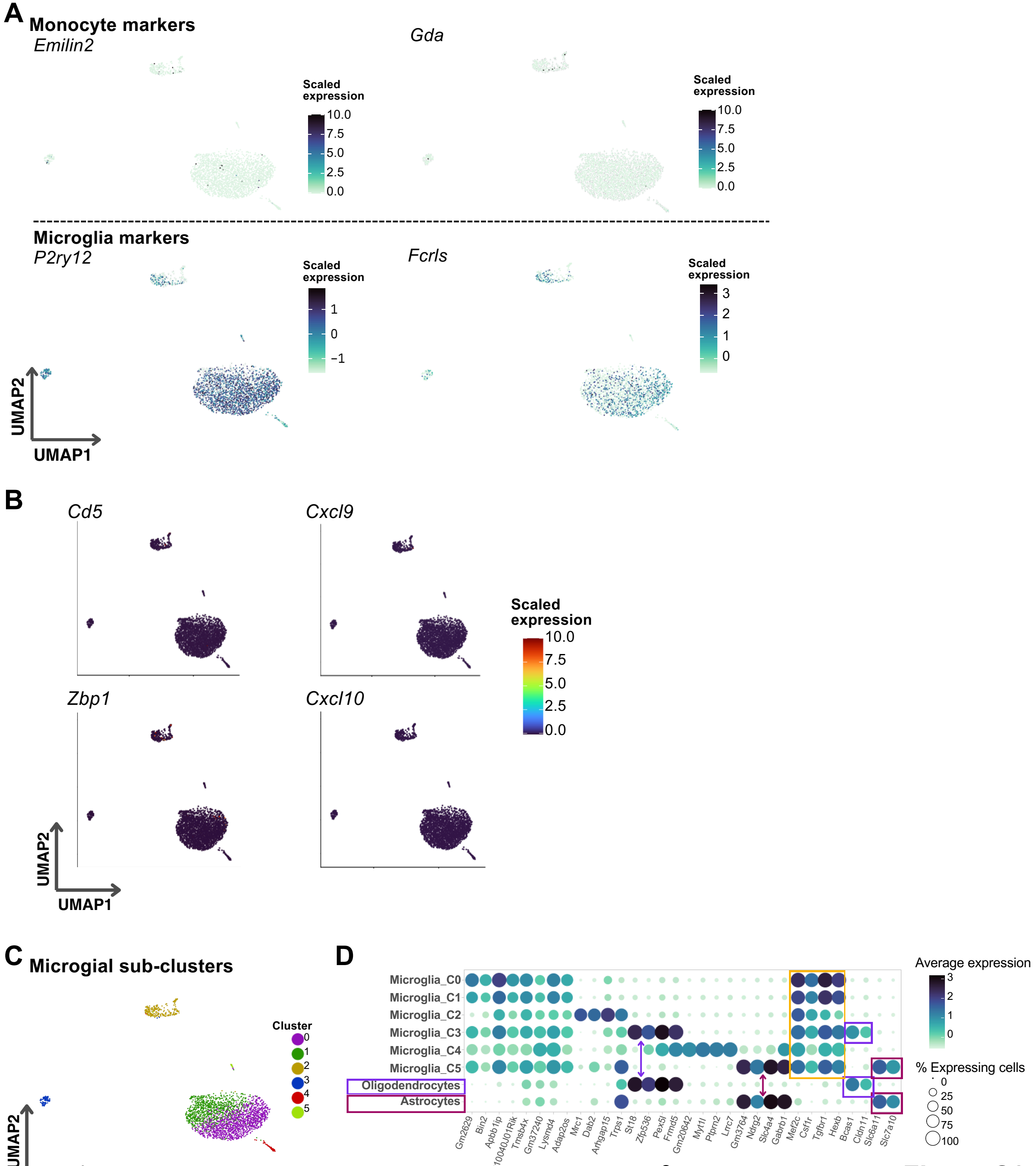
A
Monocyte markers
Emilin2
Gda
Scaled expression
10.0
7.5
5.0
2.5
0.0
Microglia markers
P2ry12
Fcrls
Scaled expression
1
0
−1
3
2
1
0
UMAP2
UMAP1
B
Cd5
Cxcl9
Zbp1
Cxcl10
Scaled expression
10.0
7.5
5.0
2.5
0.0
C
Microgial sub-clusters
Cluster
0
1
2
3
4
5
D
Microglia_C0
Microglia_C1
Microglia_C2
Microglia_C3
Microglia_C4
Microglia_C5
Oligodendrocytes
Astrocytes
Gm2629
Bin2
Apbb1ip
0610040J01Rik
Tmsb4x
Gm37240
Lysmd4
Adap2os
Mrc1
Dab2
Arhgap15
Trps1
St18
Zfp536
Pex5l
Frmd5
Gm20642
Myt1l
Ptprn2
Lrrc7
Gm3764
Ndrg2
Slc4a4
Gabrb1
Mef2c
Csf1r
Tgfbr1
Hexb
Bcas1
Cldn11
Slc6a11
Slc7a10
Gene
Average expression
% Expressing cells
25
50
75
100

**Figure S8**

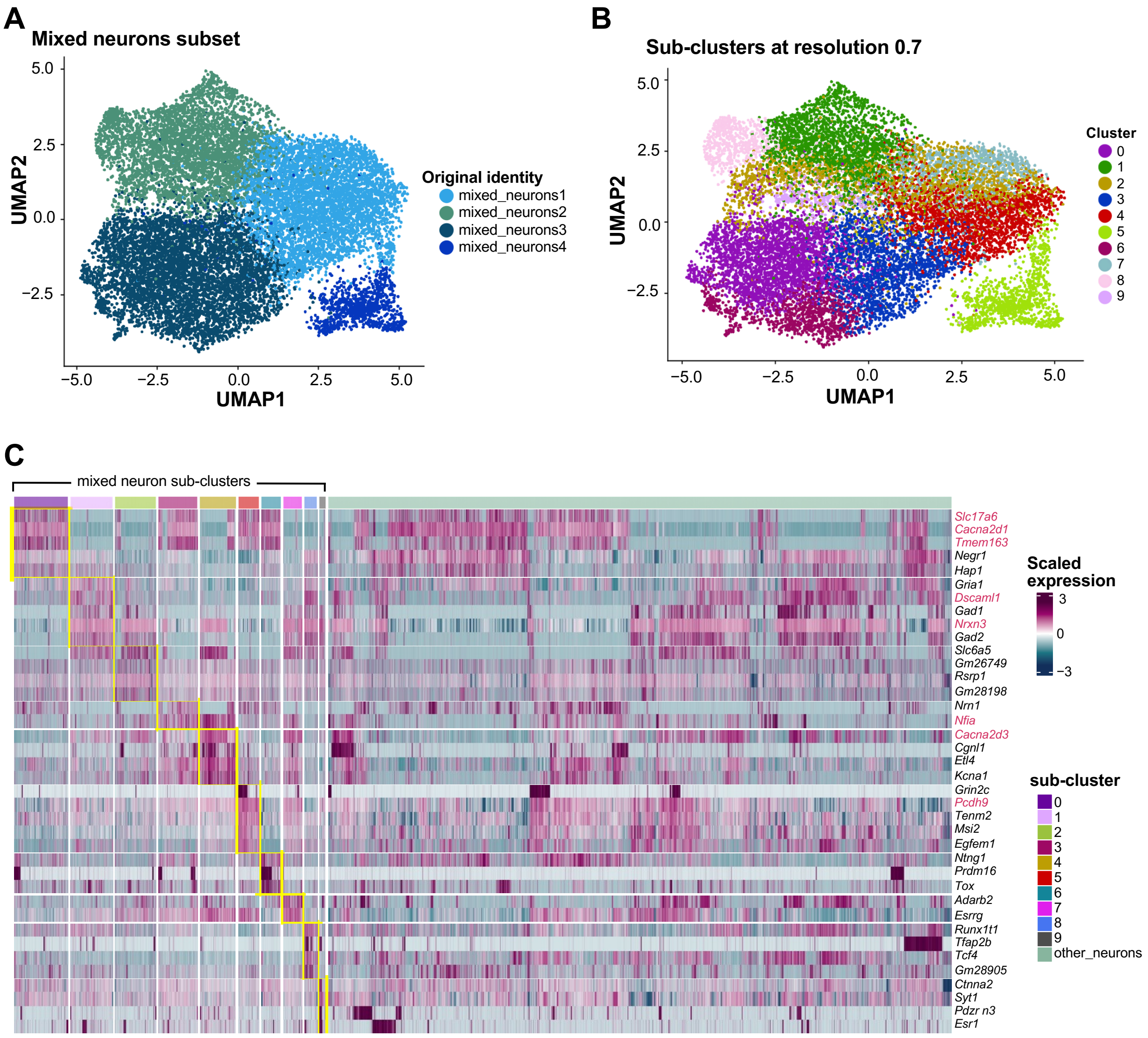
A
Mixed neurons subset
UMAP2
UMAP1
Original identity
mixed_neurons1
mixed_neurons2
mixed_neurons3
mixed_neurons4
B
Sub-clusters at resolution 0.7
UMAP2
UMAP1
Cluster
C
mixed neuron sub-clusters
Slc17a6
Cacna2d1
Tmem163
Negr1
Hap1
Gria1
Dscaml1
Gad1
Nrxn3
Gad2
Slc6a5
Gm26749
Rsrp1
Gm28198
Nrn1
Nfia
Cacna2d3
Cgnl1
Etl4
Kcna1
Grin2c
Pcdh9
Tenm2
Msi2
Egfem1
Ntng1
Prdm16
Tox
Adarb2
Esrrg
Runx1t1
Tfap2b
Tcf4
Gm28905
Ctnna2
Syt1
Pdzr n3
Esr1
Scaled expression
sub-cluster
other_neurons

Figure S9

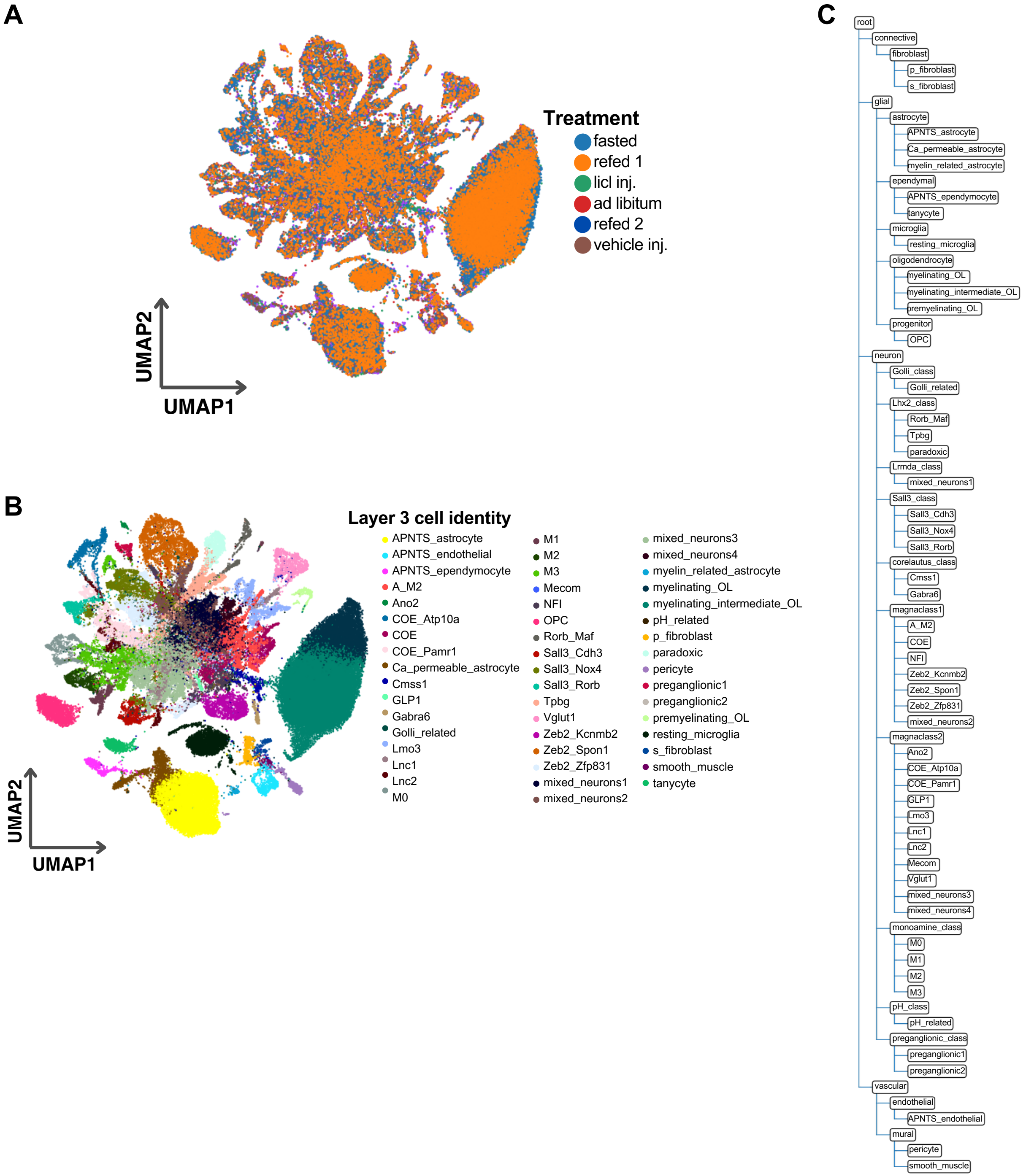
A
Treatment
fasted
refed 1
licl inj.
ad libitum
refed 2
vehicle inj.
UMAP2
UMAP1
B
Layer 3 cell identity
APNTS_astrocyte
APNTS_endothelial
APNTS_ependymocyte
A_M2
Ano2
COE_Atp10a
COE
COE_Pamr1
Ca_permeable_astrocyte
Cmss1
GLP1
Gabra6
Golli_related
Lmo3
Lnc1
Lnc2
M0
M1
M2
M3
Mecom
NFI
OPC
Rorb_Maf
Sall3_Cdh3
Sall3_Nox4
Sall3_Rorb
Tpbg
Vglut1
Zeb2_Kcnmb2
Zeb2_Spon1
Zeb2_Zfp831
mixed_neurons1
mixed_neurons2
mixed_neurons3
mixed_neurons4
myelin_related_astrocyte
myelinating_OL
myelinating_intermediate_OL
pH_related
p_fibroblast
paradoxic
pericyte
preganglionic1
preganglionic2
premyelinating_OL
resting_microglia
s_fibroblast
smooth_muscle
tanycyte
UMAP2
UMAP1
C
root
connective
fibroblast
p_fibroblast
s_fibroblast
glial
astrocyte
APNTS_astrocyte
Ca_permeable_astrocyte
myelin_related_astrocyte
ependymal
APNTS_ependymocyte
tanycyte
microglia
resting_microglia
oligodendrocyte
myelinating_OL
myelinating_intermediate_OL
premyelinating_OL
progenitor
OPC
neuron
Golli_class
Golli_related
Lhx2_class
Rorb_Maf
Tpbg
paradoxic
Lrmda_class
mixed_neurons1
Sall3_class
Sall3_Cdh3
Sall3_Nox4
Sall3_Rorb
corelautus_class
Cmss1
Gabra6
magnaclass1
A_M2
COE
NFI
Zeb2_Kcnmb2
Zeb2_Spon1
Zeb2_Zfp831
mixed_neurons2
magnaclass2
Ano2
COE_Atp10a
COE_Pamr1
GLP1
Lmo3
Lnc1
Lnc2
Mecom
Vglut1
mixed_neurons3
mixed_neurons4
monoamine_class
M0
M1
M2
M3
pH_class
pH_related
preganglionic_class
preganglionic1
preganglionic2
vascular
endothelial
APNTS_endothelial
mural
pericyte
smooth_muscle

**Figure S10**

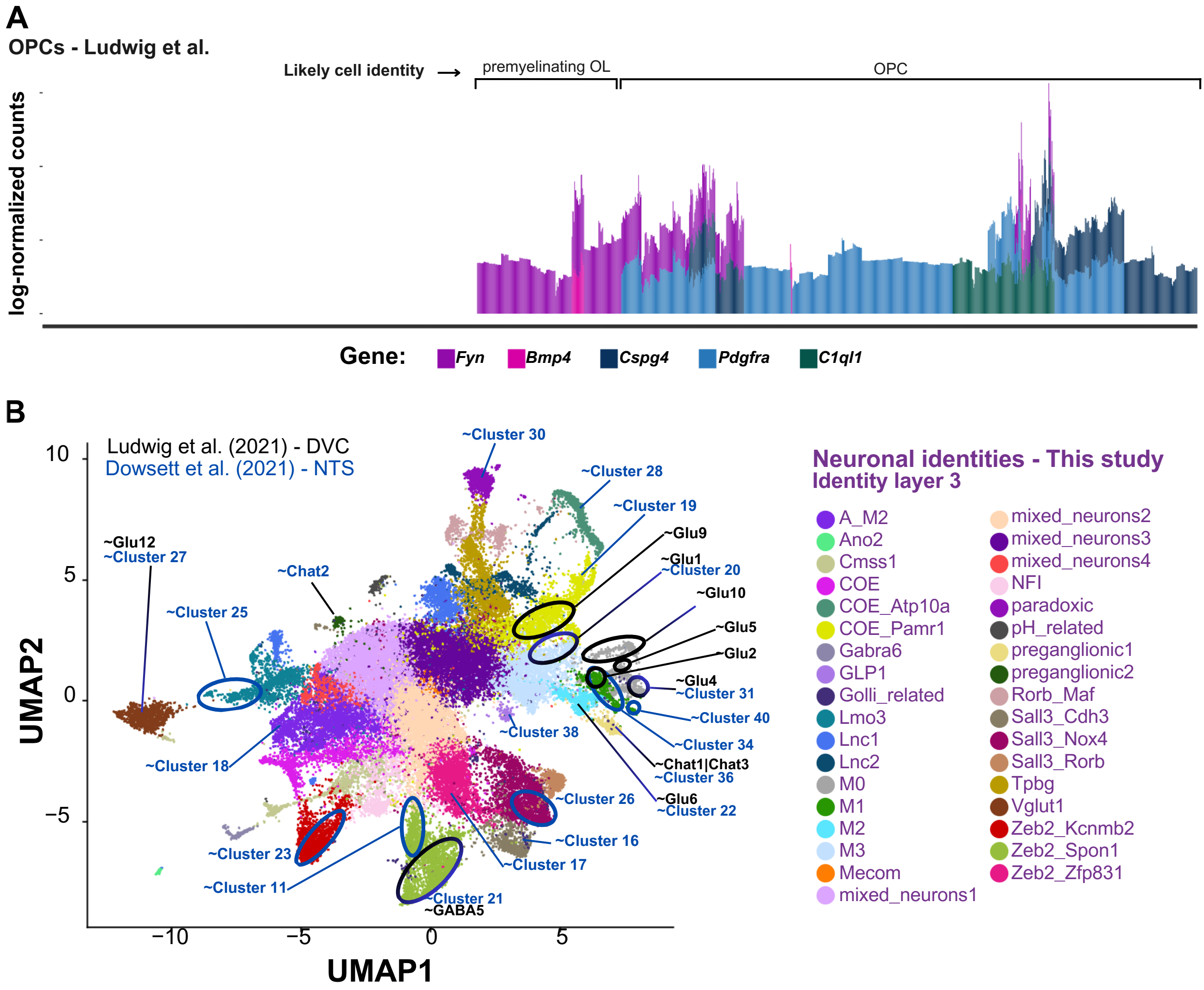
A
OPCs - Ludwig et al.
Likely cell identity
premyelinating OL
OPC
log-normalized counts
Gene:
Fyn
Bmp4
Cspg4
Pdgfra
C1ql1
B
Ludwig et al. (2021) - DVC
Dowsett et al. (2021) - NTS
UMAP2
UMAP1
~Cluster 30
~Cluster 28
~Cluster 19
~Glu9
~Glu1
~Cluster 20
~Glu10
~Glu5
~Glu2
~Glu4
~Cluster 31
~Cluster 40
~Cluster 34
~Chat1|Chat3
~Cluster 36
~Glu6
~Cluster 22
~Cluster 38
~Cluster 26
~Cluster 16
~Cluster 17
~Cluster 21
~GABA5
~Cluster 11
~Cluster 23
~Cluster 18
~Cluster 25
~Chat2
~Glu12
~Cluster 27
Neuronal identities - This study
Identity layer 3
A_M2
Ano2
Cmss1
COE
COE_Atp10a
COE_Pamr1
Gabra6
GLP1
Golli_related
Lmo3
Lnc1
Lnc2
M0
M1
M2
M3
Mecom
mixed_neurons1
mixed_neurons2
mixed_neurons3
mixed_neurons4
NFI
paradoxic
pH_related
preganglionic1
preganglionic2
Rorb_Maf
Sall3_Cdh3
Sall3_Nox4
Sall3_Rorb
Tpbg
Vglut1
Zeb2_Kcnmb2
Zeb2_Spon1
Zeb2_Zfp831

**Figure S11**

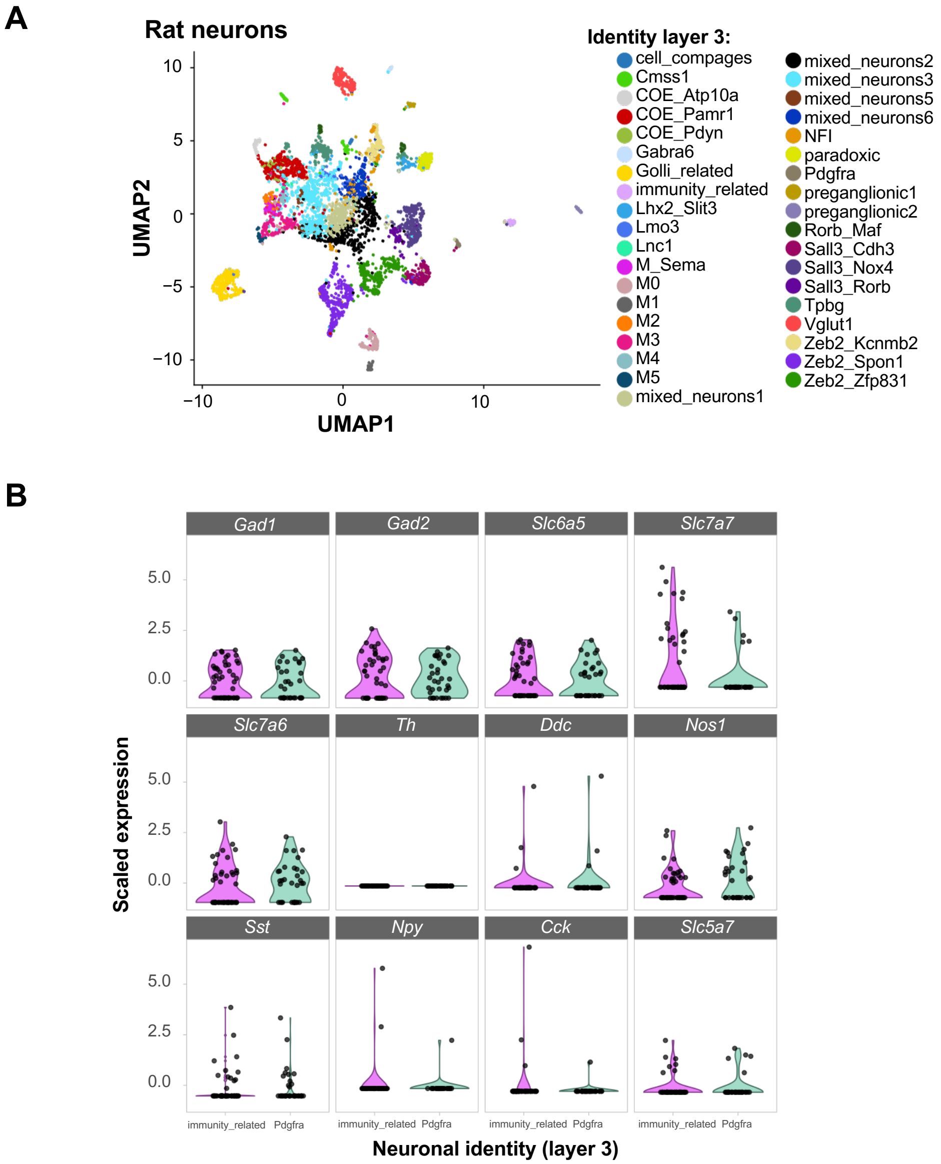
A
Rat neurons
UMAP1
UMAP2
Identity layer 3:
cell_compages
Cmss1
COE_Atp10a
COE_Pamr1
COE_Pdyn
Gabra6
Golli_related
immunity_related
Lhx2_Slit3
Lmo3
Lnc1
M_Sema
M0
M1
M2
M3
M4
M5
mixed_neurons1
mixed_neurons2
mixed_neurons3
mixed_neurons5
mixed_neurons6
NFI
paradoxic
Pdgfra
preganglionic1
preganglionic2
Rorb_Maf
Sall3_Cdh3
Sall3_Nox4
Sall3_Rorb
Tpbg
Vglut1
Zeb2_Kcnmb2
Zeb2_Spon1
Zeb2_Zfp831
B
Gad1
Gad2
Slc6a5
Slc7a7
Slc7a6
Th
Ddc
Nos1
Sst
Npy
Cck
Slc5a7
Scaled expression
immunity_related
Pdgfra
Neuronal identity (layer 3)

Figure S12

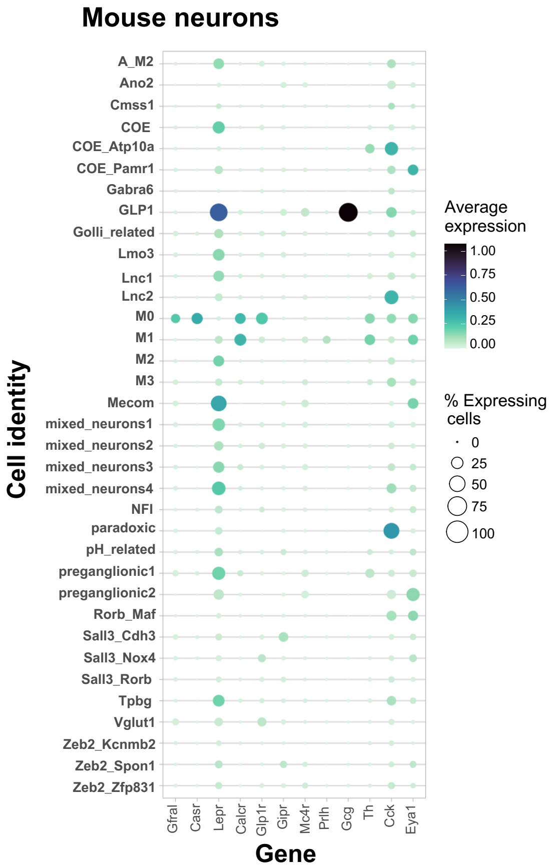

Mouse neurons
Cell identity
A_M2
Ano2
Cmss1
COE
COE_Atp10a
COE_Pamr1
Gabra6
GLP1
Golli_related
Lmo3
Lnc1
Lnc2
M0
M1
M2
M3
Mecom
mixed_neurons1
mixed_neurons2
mixed_neurons3
mixed_neurons4
NFI
paradoxic
pH_related
preganglionic1
preganglionic2
Rorb_Maf
Sall3_Cdh3
Sall3_Nox4
Sall3_Rorb
Tpbg
Vglut1
Zeb2_Kcnmb2
Zeb2_Spon1
Zeb2_Zfp831
Gfral
Casr
Lepr
Calcr
Glp1r
Gipr
Mc4r
Prlh
Gcg
Th
Cck
Eya1
Gene
Average expression
1.00
0.75
0.50
0.25
0.00
% Expressing cells
0
25
50
75
100


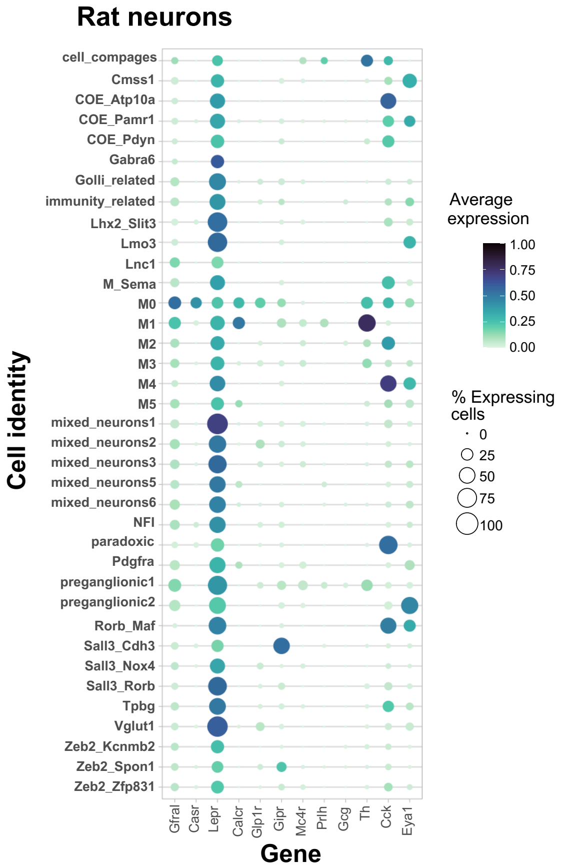

Rat neurons
Cell identity
cell_compages
Cmss1
COE_Atp10a
COE_Pamr1
COE_Pdyn
Gabra6
Golli_related
immunity_related
Lhx2_Slit3
Lmo3
Lnc1
M_Sema
M0
M1
M2
M3
M4
M5
mixed_neurons1
mixed_neurons2
mixed_neurons3
mixed_neurons5
mixed_neurons6
NFI
paradoxic
Pdgfra
preganglionic1
preganglionic2
Rorb_Maf
Sall3_Cdh3
Sall3_Nox4
Sall3_Rorb
Tpbg
Vglut1
Zeb2_Kcnmb2
Zeb2_Spon1
Zeb2_Zfp831
Gfral
Casr
Lepr
Calcr
Glp1r
Gipr
Mc4r
Prlh
Gcg
Th
Cck
Eya1
Gene
Average expression
1.00
0.75
0.50
0.25
0.00
% Expressing cells
0
25
50
75
100


**Figure S13**

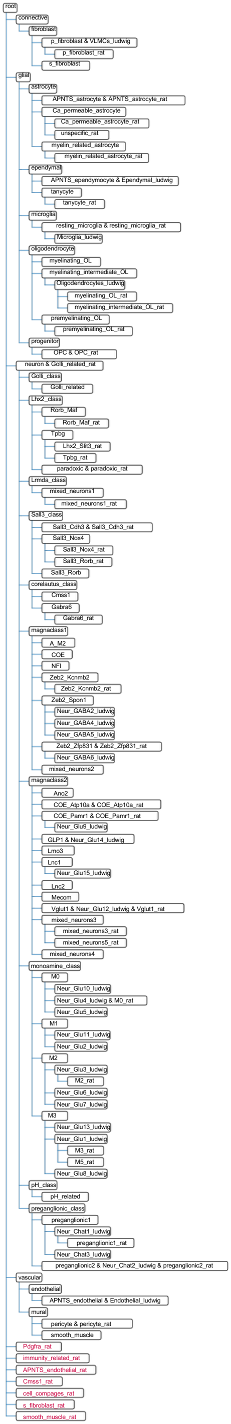
root
connective
fibroblast
p_fibroblast & VLMCs_ludwig
p_fibroblast_rat
s_fibroblast
glial
astrocyte
APNTS_astrocyte & APNTS_astrocyte_rat
Ca_permeable_astrocyte
Ca_permeable_astrocyte_rat
unspecific_rat
myelin_related_astrocyte
myelin_related_astrocyte_rat
ependymal
APNTS_ependymocyte & Ependymal_ludwig
tanycyte
tanycyte_rat
microglia
resting_microglia & resting_microglia_rat
Microglia_ludwig
oligodendrocyte
myelinating_OL
myelinating_intermediate_OL
Oligodendrocytes_ludwig
myelinating_OL_rat
myelinating_intermediate_OL_rat
premyelinating_OL
premyelinating_OL_rat
progenitor
OPC & OPC_rat
neuron & Golli_related_rat
Golli_class
Golli_related
Lhx2_class
Rorb_Maf
Rorb_Maf_rat
Tpbg
Lhx2_Slit3_rat
Tpbg_rat
paradoxic & paradoxic_rat
Lrmda_class
mixed_neurons1
mixed_neurons1_rat
Sall3_class
Sall3_Cdh3 & Sall3_Cdh3_rat
Sall3_Nox4
Sall3_Nox4_rat
Sall3_Rorb_rat
Sall3_Rorb
corelautus_class
Cmss1
Gabra6
Gabra6_rat
magnaclass1
A_M2
COE
NFI
Zeb2_Kcnmb2
Zeb2_Kcnmb2_rat
Zeb2_Spon1
Neur_GABA2_ludwig
Neur_GABA4_ludwig
Neur_GABA5_ludwig
Zeb2_Zfp831 & Zeb2_Zfp831_rat
Neur_GABA6_ludwig
mixed_neurons2
magnaclass2
Ano2
COE_Atp10a & COE_Atp10a_rat
COE_Pamr1 & COE_Pamr1_rat
Neur_Glu9_ludwig
GLP1 & Neur_Glu14_ludwig
Lmo3
Lnc1
Neur_Glu15_ludwig
Lnc2
Mecom
Vglut1 & Neur_Glu12_ludwig & Vglut1_rat
mixed_neurons3
mixed_neurons3_rat
mixed_neurons5_rat
mixed_neurons4
monoamine_class
M0
Neur_Glu10_ludwig
Neur_Glu4_ludwig & M0_rat
Neur_Glu5_ludwig
M1
Neur_Glu11_ludwig
Neur_Glu2_ludwig
M2
Neur_Glu3_ludwig
M2_rat
Neur_Glu6_ludwig
Neur_Glu7_ludwig
M3
Neur_Glu13_ludwig
Neur_Glu1_ludwig
M3_rat
M5_rat
Neur_Glu8_ludwig
pH_class
pH_related
preganglionic_class
preganglionic1
Neur_Chat1_ludwig
preganglionic1_rat
Neur_Chat3_ludwig
preganglionic2 & Neur_Chat2_ludwig & preganglionic2_rat
vascular
endothelial
APNTS_endothelial & Endothelial_ludwig
mural
pericyte & pericyte_rat
smooth_muscle
Pdgfra_rat
immunity_related_rat
APNTS_endothelial_rat
Cmss1_rat
cell_compages_rat
s_fibroblast_rat
smooth_muscle_rat

**Figure S14**

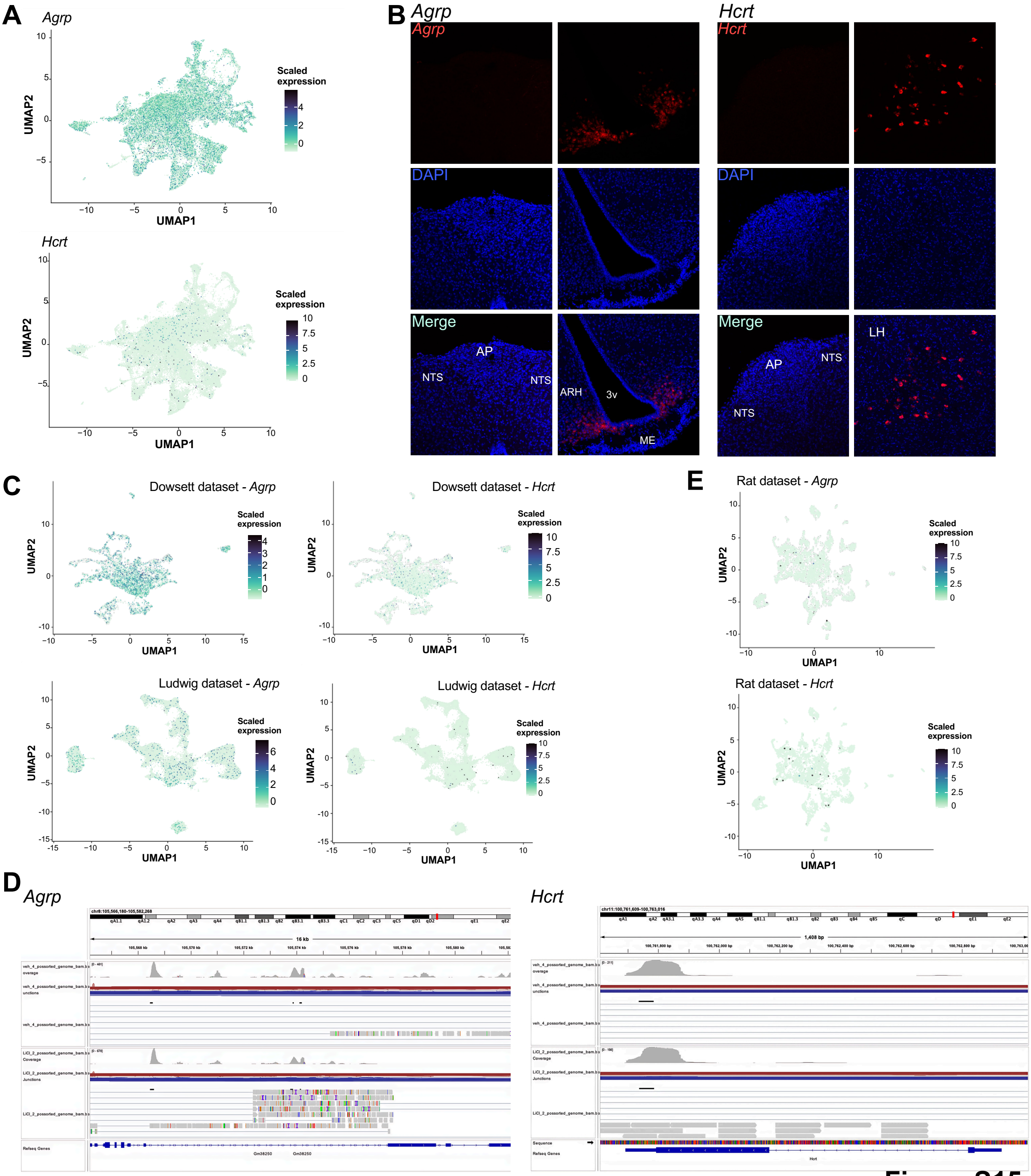
A
Agrp
Scaled expression
UMAP1
UMAP2
Hcrt
B
Agrp
Hcrt
DAPI
Merge
AP
NTS
ARH
3v
ME
LH
C
Dowsett dataset - Agrp
Dowsett dataset - Hcrt
Ludwig dataset - Agrp
Ludwig dataset - Hcrt
E
Rat dataset - Agrp
Rat dataset - Hcrt
D
Agrp
Hcrt
Refseq Genes
Sequence

Figure S15

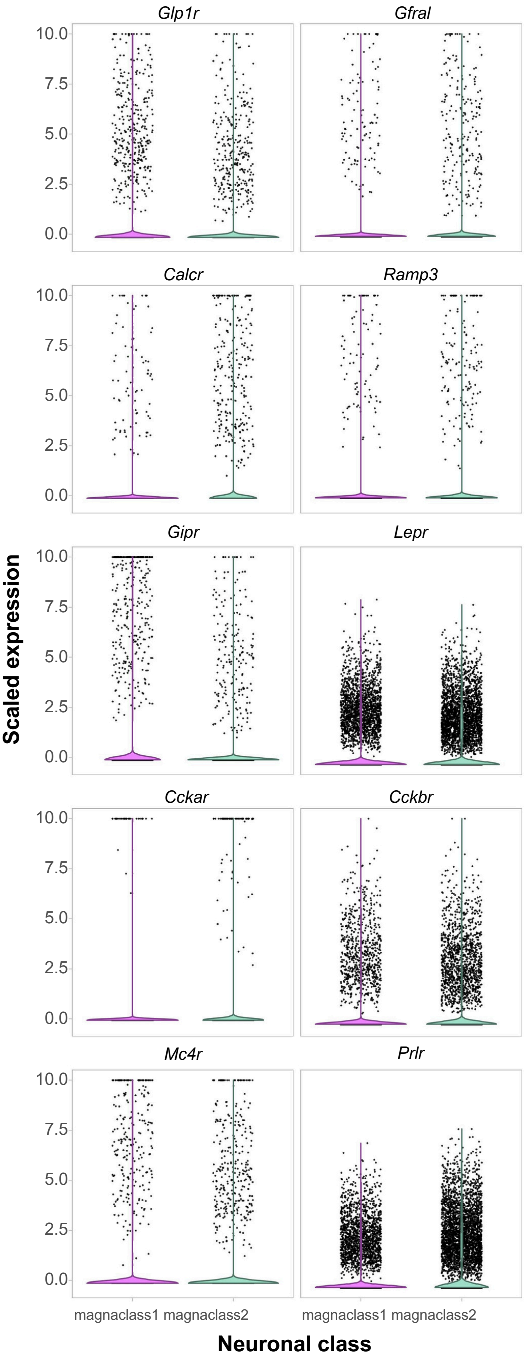
Glp1r
Gfral
Calcr
Ramp3
Gipr
Lepr
Cckar
Cckbr
Mc4r
Prlr
10.0
7.5
5.0
2.5
0.0
Scaled expression
magnaclass1 magnaclass2
magnaclass1 magnaclass2
Neuronal class

**Figure S16**
